# GWAS identifies nine nephrolithiasis susceptibility loci related with metabolic metabolic and crystallization pathways

**DOI:** 10.1101/519553

**Authors:** Chizu Tanikawa, Yoichiro Kamatani, Chikashi Terao, Masayuki Usami, Atsushi Takahashi, Yukihide Momozawa, Kichiya Suzuki, Soichi Ogishima, Atsushi Shimizu, Mamoru Satoh, Keitaro Matsuo, Haruo Mikami, Mariko Naito, Kenji Wakai, Taiki Yamaji, Norie Sawada, Motoki Iwasaki, Shoichiro Tsugane, Kenjiro Kohri, Takahiro Yasui, Yoshinori Murakami, Michiaki Kubo, Koichi Matsuda

## Abstract

Nephrolithiasis is a common urological trait disorder with acute pain. Although previous studies have identified various genetic variations associated with nephrolithiasis, the host genetic factors remain largely unidentified. To identify novel nephrolithiasis loci in the Japanese population, we performed large-scale GWAS (Genome wide association study) using 11,130 cases and 187,639 controls, followed by a replication analysis using 2,289 cases and 3,817 controls. The analysis identified 14 significant loci, including 9 novel loci on 2p23.2-3, 6p21.2, 6p12.3, 6q23.2, 16p12.3, 16q12.2, 17q23.2, 19p13.12, and 20q13.2. Interestingly, 10 of the 14 regions showed a significant association with any of 16 quantitative traits, including metabolic, kidney-related, and electrolyte traits, suggesting a common genetic background among nephrolithiasis patients and these quantitative traits. Four novel loci are related to the metabolic pathway, while the remaining 10 loci are associated with the crystallization pathway. Our findings demonstrate the crucial roles of genetic variations in the development of nephrolithiasis.

**SIGNIFICANCE STATEMENT:** Nephrolithiasis is a common urothelial disorders with frequent recurrence rate, but its genetic background is largely remained unidentified. Previous GWAS identified 6 genetic factors in total. Here we performed a GWAS using more than 200,000 samples in the Japanese populations, and identified 14 significant loci and nine of them are novel. We also found that 10 of the 14 loci showed a significant association with any of 16 quantitative traits, including metabolic, kidney-related, and electrolyte traits (BMI, eGFR, UA, Ca etc). All 14 significant loci are associate with either metabolic or crystallization pathways. Thus, our findings elucidated the underlying molecular pathogenesis of nephrolithiasis.

## INTRODUCTION

Nephrolithiasis is a common health problem that causes severe, acute back pain. The prevalence of nephrolithiasis is estimated to be 10-15% in men and 7% in women, and its prevalence increased by 70-85% between the 1980s and the 2000s in both the USA and Japan ^1, 2^. The recurrence rate of nephrolithiasis is 30-50% within 10 years after the initial episode ^3^ and occasionally leads to severe complications, such as pyelonephritis or acute renal failure. Therefore, the identification of risk factors is important for the management of nephrolithiasis. Environmental factors, such as lifestyle, obesity ^4, 5^, hypertension ^6^, and diabetes ^7^, are associated with an increased risk of nephrolithiasis. In addition, a positive family history increases the disease risk by 2.57-fold among the Japanese population ^8^, and a twin study estimated the heritability of nephrolithiasis to be 56% ^9^, suggesting a pivotal role for host genetic factors. Monogenic diseases causing hypercalciuria (CLCN5, SLC34A1, NKCC2, ROMK, and CaSR9), distal tubular acidosis (SLC4A1), hyperoxaluria (AGTXT), and cysteinuria (SLC3A1 and SLC7A9) are associated with familial nephrolithiasis syndrome ^10^. In addition, three previous GWAS in European and Japanese populations identified 6 common genetic factors on 21q22.13 (*CLDN14*) ^11^, 5q35.3, 7p14.3, 13q14.1 ^12^, 1p36.12 (*ALPL*), and 3q21.1 (*CASR*) ^13^. These loci were shown to be quantitative loci for ALP and serum phosphate (1p36.12), PTH and urinary Mg/Ca ratio (5q35.3 and 21q22.13), bone mineral density (7p14.3 and 21q22.13), serum Ca (13q14.1 and 3q21.1) and kidney function (5q35.3), suggesting the underlying mechanism of these traits in the regulation of cations, kidney function, and mineralization. To further elucidate the genetic factors related to nephrolithiasis, we conducted GWAS using more than 13,000 Japanese cases and 190,000 control samples.

## METHODS

### Study participants

The characteristics of each cohort are shown in **Table 1**. The diagnosis of nephrolithiasis in patients enrolled in screening 1 was confirmed by physicians, while we selected cases for screening 2 based on medical history obtained by medical coordinators in each hospital. DNA samples of 12,846 nephrolithiasis patients and 162,394 controls were obtained from BioBank Japan ^14, 15^. The 28,867 controls were from four population-based cohorts, including the JPHC (Japan Public Health Center)-based prospective study ^16^, the J-MICC (Japan Multi-Institutional Collaborative Cohort) study ^17^, IMM (Iwate Tohoku Medical Megabank Organization) and ToMMo (Tohoku Medical Megabank Organization) ^18^. A total of 573 nephrolithiasis patients and 195 healthy controls were recruited at the Nagoya City University ^19^. The diagnosis of nephrolithiasis in patients from Nagoya City University (n = 573) and BioBank Japan in the screening 1 (n = 6,246) was confirmed by a clinician. Cases in screening 2 (n = 4,884) and the replication (n = 1,716) were selected from samples in BioBank Japan and were based on medical history obtained by questionnaire or medical records. Genomic DNA was extracted from peripheral blood leukocytes using a standard method. All participants provided written informed consent, and the project was approved by the ethical committees at each institute.

**Table 1.** Characteristics of study population

| Stage | Sample type | Source | Platform | Number of samples | Female (%) | Age (mean +/- SD) |
| --- | --- | --- | --- | --- | --- | --- |
| screening 1 | kidney stone | BBJ | OmniExpressExome or<br>OmniExpress+HumanExome | 6,246 | 1,531 (24.5%) | 54.7 +/- 13.8 |
|  | control | JPHC, J-MICC,<br>ToMMo | OmniExpressExome | 28,867 | 17,490 (60.6%) | 56.3 +/- 10.0 |
| screening 2 | kidney stone | BBJ | OmniExpressExome or<br>OmniExpress+HumanExome | 4,884 | 1,303 (26.7%) | 62.5 +/- 12.0 |
|  | control | BBJ | OmniExpressExome or<br>OmniExpress+HumanExome | 158,772 | 75,209 (47.4%) | 63.0 +/- 14.4 |
| replication | kidney stone | BBJ, NCU | Invader assay | 2,289 | 416 (18.3%) | 63.3 +/- 13.9 |
|  | control | BBJ, NCU | Invader assay | 3,817 | 1,869 (49.0%) | 63.7 +/- 13.9 |

Note: BBJ:Biobank Japan, JPHC: Japan Public Health Center-based Prospective Study, J-MICC: Japan Multi-Institutional Collaborative Cohort study, ToMMo:Tohoku Medical Megabank Organization, NCU: Nagoya City University

### Genotyping and imputation

The strategy of our screening is shown in **Supplemental Figure 1**. In screening 1, we used clinically confirmed nephrolithiasis cases and healthy controls. In screening 2, we selected 4,884 nephrolithiasis cases and 158,772 controls from BioBank Japan based on medical records or self-reporting. In previous studies ^20-22^, all 11,130 nephrolithiasis cases and 187,639 controls (screening 1 and 2) were genotyped with an Illumina HumanOmniExpressExome BeadChip or a combination of the Illumina HumanOmniExpress and HumanExome BeadChips (**Table 1**). We excluded (i) samples with a call rate < 0.98, (ii) samples from closely related individuals identified by identity-by-descent analysis, (iii) sex-mismatched samples with a lack of information, and (iv) samples from non–East Asian outliers identified by principal component analysis of the studied samples and the three major reference populations (Africans, Europeans, and East Asians) in the International HapMap Project ^23^. We then applied standard quality-control criteria for variants, excluding those with (i) SNP call rate < 0.99, (ii) minor allele frequency < 1%, and (iii) Hardy–Weinberg equilibrium P value < 1.0 × 10^−6^. We prephased the genotypes with MACH ^24^ and imputed dosages with minimac and the 1000 Genomes Project Phase 1 (version 3) East Asian reference haplotypes ^25^. Imputed SNPs with an imputation quality Rsq < 0.4 were excluded from the subsequent association analysis.

**Figure 1.**
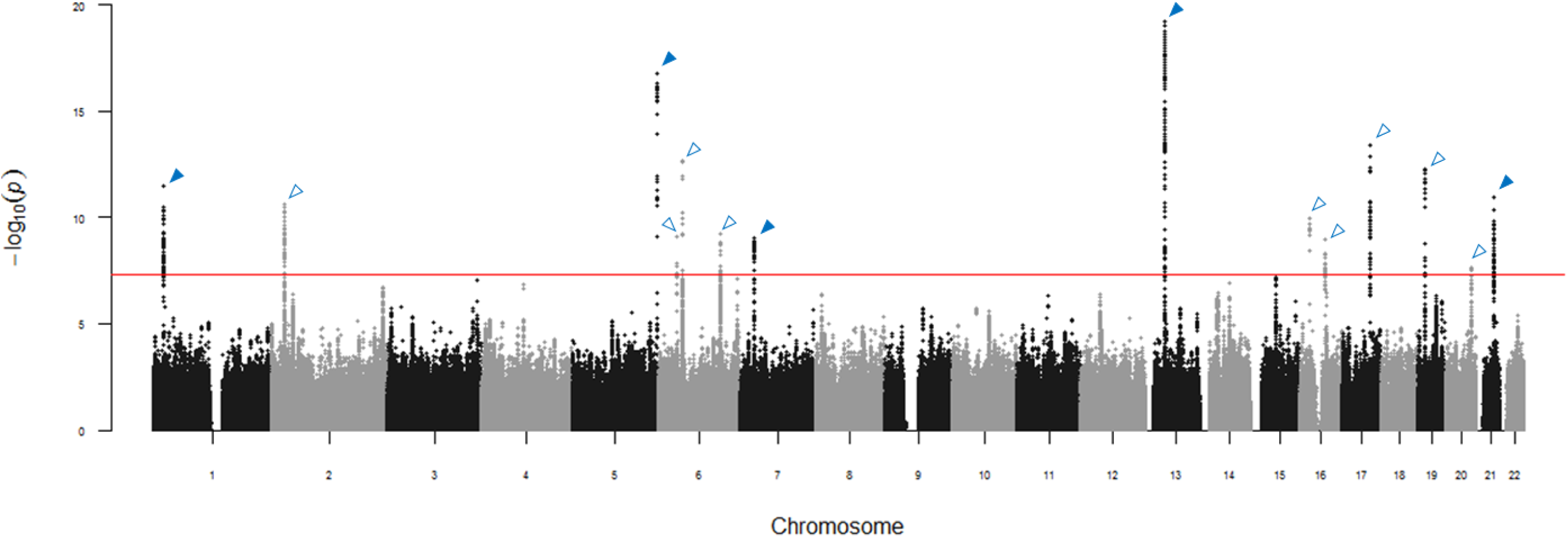
Manhattan plot showing the genome-wide *P* values of association. The genome-wide *P*-values of 6,603,247 autosomal SNPs in 11,130 cases and 187,639 controls from the meta-analysis of screening 1 and screening 2 are shown. Fourteen loci are identified risk loci, and open arrowheads indicate newly identified loci. The red horizontal line represents the genome-wide significance threshold of *P* = 5.0 × 10^−8^.

### Genome-wide association analysis

We conducted a GWAS using a logistic regression model by incorporating age, sex, and the top 10 PCs as covariates. Meta-analysis of screening 1 and screening 2 was conducted by using METAL ^26^. Heterogeneity between studies was examined using Cochran’s Q test ^27^. To estimate the genetic correlation, a bivariate LD score regression ^28^ was conducted using the results from the GWAS of screening 1 and screening 2 with the LD scores for the East Asian population ^22^. We calculated the genomic inflation factor λGC in R. λGC adjusted to a sample size of 1,000 (*λ*_1,000_) was calculated using the following formula ^29^, as large sample sizes cause inflated λGC values ^30^): *λ*_1,000_ = 1+(1-*λ*_obs_) × (1/*n*_cases_+1/*n*_controls_)/(1/1,000_cases_+1/1,000_controls_). A quantile-quantile plot was drawn using R.

A Manhattan plot of the associations was constructed by plotting -log10 (P values) against chromosome position using R. We generated regional plots with LocusZoom (v. 1.3) ^31^. A forest plot was drawn using R.

### Replication study

Among the genome-wide significant SNPs with P < 5 × 10^−8^ from the meta-analysis of screening 1 and screening 2, we selected 17 SNPs within 14 loci by linkage disequilibrium (LD) analysis (r2 < 0.2). We genotyped 17 SNPs using 2,320 nephrolithiasis cases and 3,961 controls by the multiplex PCR-based Invader assay (Third Wave Technologies). The meta-analysis of screening 1, screening 2, and the replication was conducted using METAL in the same way as in the screening step. The threshold of heterogeneity was P < 0.05/17.

### Pleiotropy analysis

The GWAS results for metabolic traits (BMI, TC, HDL-C, LDL-C, TG, BS, and HbA1c), kidney-related traits (BUN, sCr, eGFR, and UA), and electrolytes (Na, K, Cl, Ca, and P) were used in the pleiotropy analysis ^22^. To assess the colocalization of causal SNPs, we conducted a conditional logistic regression analysis with conditioning of the top SNPs of the Japanese QTL GWAS. The tested SNPs were selected to be located within 1 Mb of the typing SNP. We downloaded the NHGRI-EBI GWAS catalog (v1.0.1).

### eQTL analysis

Expression data for specific tissues obtained from the GTEx Portal were used to evaluate whether the variants in the genomic loci identified in this study affect gene expression (eQTL analysis).

### Subgroup analysis

We used the subjects in screening 1 for the subgroup analysis. Each subgroup was analyzed using a logistic regression model with an adjustment for sex and age. Control samples were the same as those used in screening 1. Then, comparisons between subgroups were performed using a logistic regression model with an adjustment for sex and age.

### Weighted genetic risk score

A total of 17 associated SNPs (P < 5.0 × 10^−8^ from the meta-analysis of screening 1 and screening 2) were used. The wGRS (weighted genetic risk score) model was established to incorporate the estimate (weight) from the meta-analysis of screening 1 and screening 2 for each of the 17 associated SNPs. The cumulative risk scores were calculated by multiplying the weight of each SNP by the frequency of the risk alleles for the SNP carried by the individual, and the sum across the total number of SNPs was considered. Subsequently, the risk scores were classified into five quantiles on the basis of wGRS. P values, odds ratios, and 95% confidence intervals were evaluated using the first quantile as the reference.

### Statistical Analysis

We conducted GWAS and subgroup analysis using a logistic regression model by using plink. Meta-analysis was conducted by using METAL. The wGRS (weighted genetic risk score) model was evaluated by Fisher’s exact test by using R. Summary statistics and primary genotyping data that support the findings of this study can be found at NDBC with the accession code hum0014 (http://humandbs.biosciencedbc.jp/), after acceptance of this paper.

## RESULTS

### Genome wide association study of nephrolithiasis

A total of 6,246 nephrolithiasis patients and 28,867 controls (screening 1) and 4,884 nephrolithiasis cases and 158,772 controls (screening 2) were analyzed in the screening stage. All samples were genotyped using the Illumina OmniExpressExome or OmniExpress+HumanExome BeadChip in the previous analyses (**Table 1** and **Supplemental Figure 1**) ^20-22^. We excluded samples after performing a standard quality control (QC) procedure. Then, we selected 509,872 SNPs for genome-wide imputation (MAF ≥ 0.01, HWE ≥ 1×10^−6^, and call rate ≥ 0.99) and obtained imputed dosages of 6,603,247 SNPs (RSQR ≥ 0.4). We tested the association with nephrolithiasis susceptibility using a logistic regression analysis with age, sex, and the top 10 PCs as covariates. A meta-analysis of screening 1 and screening 2 indicated that 822 SNPs in 14 genomic regions including five previously reported loci showed significant association with p-value of less than 5 x10^−8^ (**Figure 1** and **Supplemental Table 1**). The genomic inflation factor *λ* and *λ*_1000_ was 1.164 and 1.008, respectively (**Supplemental Figure 2**) ^29^. The LD score regression analysis ^32^ of screening 1 and screening 2 revealed a genetic correlation score of 0.92, indicating that the results were consistent between the two studies.

**Figure 2.**
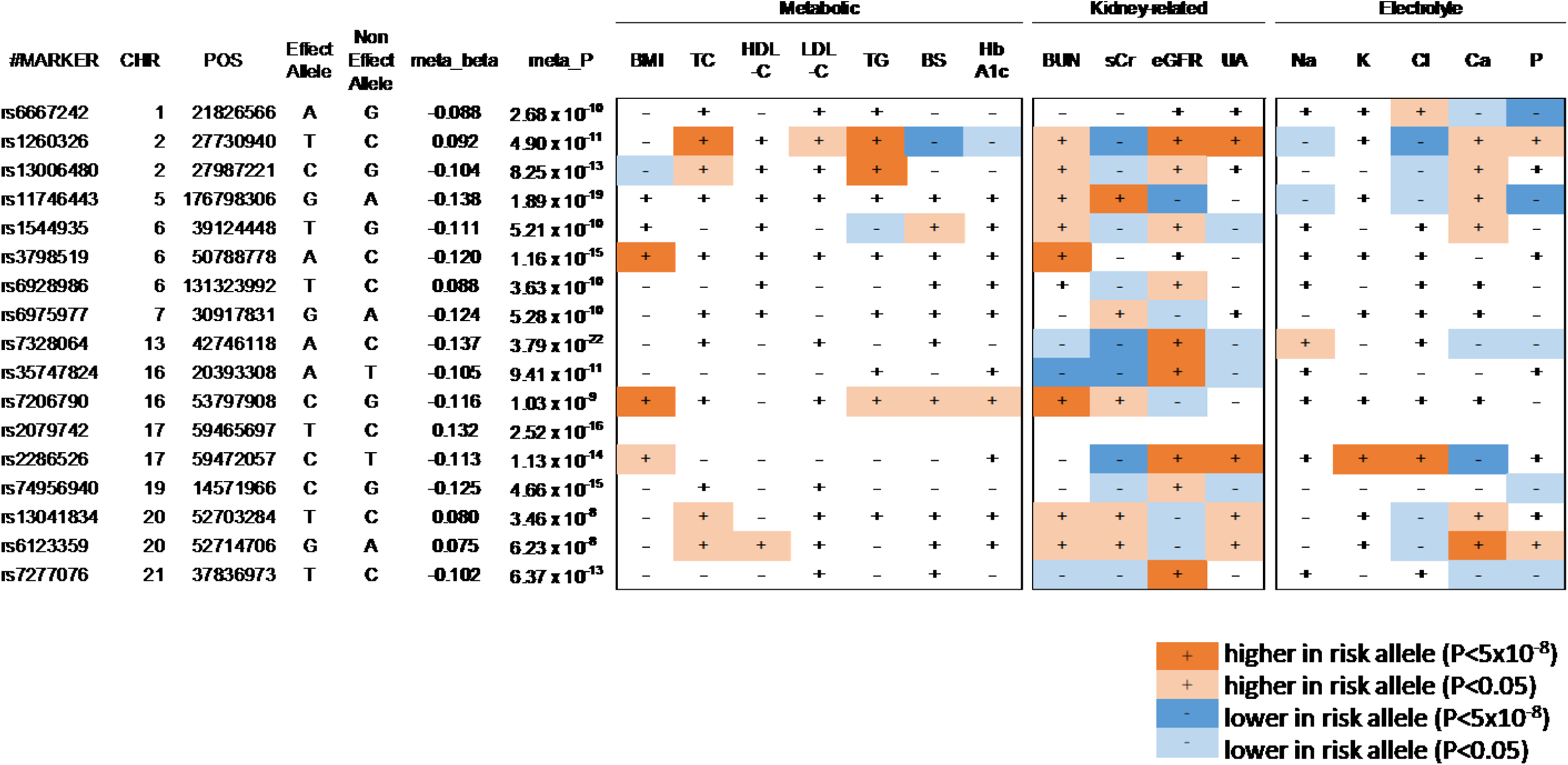
Summary of pleiotropy analysis. The association of 17 loci with 16 quantitative traits in three categories, including metabolic, kidney-related, and electrolyte traits, are shown.

We selected 17 SNPs in 14 genomic regions with significant associations (P < 5 × 10^−8^) in the replication analysis based on linkage disequilibrium (LD, *r*^2^< 0.2) and conditioned analysis adjusted by lead SNPs in each region (**Supplemental Table 2** and **Supplemental Figure 3**). We substituted the top-ranked SNPs rs6667242, rs11746443, rs3798519, and rs74956940 on 1p36.12, 5q35.3, 6p12.3, and 19p13.12, respectively, for rs1697420, rs10866705, rs62405419, and rs2241358 because we could not design probes for invader assay. These 17 SNPs were analyzed using independent Japanese samples consisting of 2,289 cases and 3,817 controls ^19^ by a multiplex-polymerase chain reaction-based Invader assay ^33^. As a result, the risk alleles were consistent, and the effect sizes were similar among the three studies (screening 1, screening 2, and replication) for 17 SNPs. A meta-analysis revealed that 16 SNPs within 14 regions including nine novel regions exceeded the genome-wide significant threshold (P < 5 × 10^−8^) (**Table 2**). These SNPs included: *GCKR-C2orf16-ZNF512-CCDC121-GPN1-SUPT7L-SLC4A1AP-MRPL33-RBKS* (2p23.2-3), *SAYSD1-KCNK5* (6p21.2), *TFAP2D-TFAP2B* (6p12.3), *EPB41L2* (6q23.2), *PDILT* (16p12.3), *FTO* (16q12.2), *BCAS3-TBX2-C17orf82* (17q23.2), *PKN1-PTGER1-GIPC1* (19p13.12), and *BCAS1* (20q13.2). In addition, all 6 previously identified SNPs showed a significant association (P = 0.0297 – 2.76 × 10^−18^, **Supplemental Table 3**).

**Table 2.** The result of association analysis of kidney stone in each stage

| SNP | Chr | Position | Region | Effect Allele | Non Effect Allele | RSQR | gene | relative loc | screening 1 |  | screening 2 |  | meta_screening |  |  |  | replication |  | meta |  |  |  |
| --- | --- | --- | --- | --- | --- | --- | --- | --- | --- | --- | --- | --- | --- | --- | --- | --- | --- | --- | --- | --- | --- | --- |
|  |  |  |  |  |  |  |  |  | OR<sup>a</sup> | P | OR<sup>a</sup> | P | OR<sup>a</sup> | P | Q<sup>b</sup> | I<sup>c</sup> | OR<sup>a</sup> | P | OR<sup>a</sup> | P | Q<sup>b</sup> | I<sup>c</sup> |
| rs6667242 | 1 | 21826566 | 1p36.12 | A | G | 1.000 | ALPL | -9292 | 0.915 | 2.87 x 10<sup>-5</sup> | 0.898 | 3.07 x 10<sup>-7</sup> | 0.907 | 4.12 x 10<sup>-11</sup> | 0.545 | 0.0 | 0.987 | 7.47 x 10<sup>-1</sup> | 0.916 | 2.68 x 10<sup>-10</sup> | 0.114 | 54.0 |
| rs1260326 | 2 | 27730940 | 2p23.3 | T | C | 0.997 | GCKR | 0 | 1.121 | 6.39 x 10<sup>-8</sup> | 1.084 | 1.03 x 10<sup>-4</sup> | 1.102 | 5.17 x 10<sup>-11</sup> | 0.267 | 19.0 | 1.050 | 2.27 x 10<sup>-1</sup> | 1.096 | 4.90 x 10<sup>-11</sup> | 0.275 | 22.5 |
| rs13006480 | 2 | 27987221 | 2p23.2 | C | G | 0.903 | MRPL33 | -7363 | 0.873 | 9.81 x 10<sup>-10</sup> | 0.931 | 1.12 x 10<sup>-3</sup> | 0.901 | 2.30 x 10<sup>-11</sup> | 0.040 | 76.4 | 0.900 | 8.98 x 10<sup>-3</sup> | 0.901 | 8.25 x 10<sup>-13</sup> | 0.118 | 53.2 |
| rs11746443 | 5 | 176798306 | 5q35.3 | G | A | 1.000 | RGS14 | 0 | 0.848 | 1.59 x 10<sup>-12</sup> | 0.899 | 4.50 x 10<sup>-6</sup> | 0.873 | 7.98 x 10<sup>-17</sup> | 0.070 | 69.6 | 0.857 | 5.46 x 10<sup>-4</sup> | 0.871 | 1.89 x 10<sup>-19</sup> | 0.185 | 40.8 |
| rs1544935 | 6 | 39124448 | 6p21.2 | T | G | 1.000 | KCNK5 | 32299 | 0.862 | 3.45 x 10<sup>-8</sup> | 0.916 | 1.07 x 10<sup>-3</sup> | 0.889 | 7.53 x 10<sup>-10</sup> | 0.110 | 60.8 | 0.941 | 2.39 x 10<sup>-1</sup> | 0.895 | 5.21 x 10<sup>-10</sup> | 0.166 | 44.2 |
| rs3798519 | 6 | 50788778 | 6p12.3 | A | C | 0.981 | TFAP2B | 0 | 0.906 | 2.01 x 10<sup>-5</sup> | 0.875 | 2.19 x 10<sup>-9</sup> | 0.890 | 2.60 x 10<sup>-13</sup> | 0.272 | 17.3 | 0.869 | 1.28 x 10<sup>-3</sup> | 0.887 | 1.16 x 10<sup>-15</sup> | 0.483 | 0.0 |
| rs6928986 | 6 | 131323992 | 6q23.2 | T | C | 0.998 | EPB41L2 | 0 | 1.103 | 4.91 x 10<sup>-6</sup> | 1.090 | 5.16 x 10<sup>-5</sup> | 1.096 | 5.81 x 10<sup>-10</sup> | 0.686 | 0.0 | 1.056 | 2.23 x 10<sup>-1</sup> | 1.092 | 3.63 x 10<sup>-10</sup> | 0.665 | 0.0 |
| rs6975977 | 7 | 30917831 | 7p14.3 | G | A | 0.983 | INMT-FAM188B | 0 | 0.879 | 1.82 x 10<sup>-5</sup> | 0.877 | 1.09 x 10<sup>-5</sup> | 0.878 | 8.89 x 10<sup>-10</sup> | 0.962 | 0.0 | 0.927 | 1.91 x 10<sup>-1</sup> | 0.884 | 5.28 x 10<sup>-10</sup> | 0.681 | 0.0 |
| rs7328064 | 13 | 42746118 | 13q14.11 | A | C | 0.937 | DGKH | 0 | 0.840 | 5.48 x 10<sup>-16</sup> | 0.898 | 5.19 x 10<sup>-7</sup> | 0.870 | 5.87 x 10<sup>-20</sup> | 0.028 | 79.4 | 0.886 | 1.95 x 10<sup>-3</sup> | 0.872 | 3.79 x 10<sup>-22</sup> | 0.082 | 60.0 |
| rs35747824 | 16 | 20393308 | 16p12.3 | A | T | 0.988 | PDILT | 0 | 0.851 | 4.34 x 10<sup>-11</sup> | 0.944 | 1.95 x 10<sup>-2</sup> | 0.894 | 1.15 x 10<sup>-10</sup> | 0.003 | 88.7 | 0.943 | 2.23 x 10<sup>-1</sup> | 0.900 | 9.41 x 10<sup>-11</sup> | 0.007 | 80.1 |
| rs7206790 | 16 | 53797908 | 16q12.2 | C | G | 0.741 | FTO | 0 | 0.884 | 5.07 x 10<sup>-5</sup> | 0.877 | 9.14 x 10<sup>-6</sup> | 0.881 | 1.08 x 10<sup>-9</sup> | 0.848 | 0.0 | 0.942 | 2.00 x 10<sup>-1</sup> | 0.890 | 1.03 x 10<sup>-9</sup> | 0.414 | 0.0 |
| rs2079742 | 17 | 59465697 | 17q23.2 | T | C | 0.685 | BCAS3 | 0 | 1.175 | 1.89 x 10<sup>-10</sup> | 1.110 | 2.76 x 10<sup>-5</sup> | 1.143 | 4.29 x 10<sup>-14</sup> | 0.107 | 61.5 | 1.137 | 1.08 x 10<sup>-3</sup> | 1.141 | 2.52 x 10<sup>-16</sup> | 0.272 | 23.1 |
| rs2286526 | 17 | 59472057 | 17q23.2 | C | T | 0.976 | LOC645722 | 0 | 0.879 | 4.00 x 10<sup>-9</sup> | 0.910 | 1.40 x 10<sup>-5</sup> | 0.894 | 6.04 x 10<sup>-13</sup> | 0.247 | 25.3 | 0.887 | 4.30 x 10<sup>-3</sup> | 0.894 | 1.13 x 10<sup>-14</sup> | 0.529 | 0.0 |
| rs74956940 | 19 | 14571966 | 19p13.12 | C | G | 0.966 | PKN1 | 0 | 0.876 | 5.12 x 10<sup>-8</sup> | 0.894 | 2.75 x 10<sup>-6</sup> | 0.885 | 6.53 x 10<sup>-13</sup> | 0.556 | 0.0 | 0.869 | 1.81 x 10<sup>-3</sup> | 0.883 | 4.66 x 10<sup>-15</sup> | 0.777 | 0.0 |
| rs13041834 | 20 | 52703284 | 20q13.2 | T | C | 0.998 | BCAS1 | -15980 | 1.113 | 1.21 x 10<sup>-6</sup> | 1.068 | 2.41 x 10<sup>-3</sup> | 1.090 | 2.69 x 10<sup>-8</sup> | 0.188 | 42.4 | 1.038 | 3.68 x 10<sup>-1</sup> | 1.084 | 3.46 x 10<sup>-8</sup> | 0.223 | 33.5 |
| rs6123359 | 20 | 52714706 | 20q13.2 | G | A | 0.980 | BCAS1 | -27402 | 1.102 | 4.85 x 10<sup>-6</sup> | 1.072 | 8.92 x 10<sup>-4</sup> | 1.087 | 2.28 x 10<sup>-8</sup> | 0.346 | 0.0 | 1.018 | 6.52 x 10<sup>-1</sup> | 1.078 | 6.23 x 10<sup>-8</sup> | 0.195 | 38.8 |
| rs7277076 | 21 | 37836973 | 21q22.13 | T | C | 0.928 | CLDN14 | 0 | 0.894 | 2.24 x 10<sup>-7</sup> | 0.909 | 7.65 x 10<sup>-6</sup> | 0.902 | 1.14 x 10<sup>-11</sup> | 0.576 | 0.0 | 0.912 | 1.84 x 10<sup>-2</sup> | 0.903 | 6.37 x 10<sup>-13</sup> | 0.834 | 0.0 |

<sup>a</sup>Non-effect alleles were considered as references. <sup>b</sup>p-value for Cochran's Q statistic. <sup>c</sup>I<sup>2</sup> heterogeneity index.

**Figure 3.**
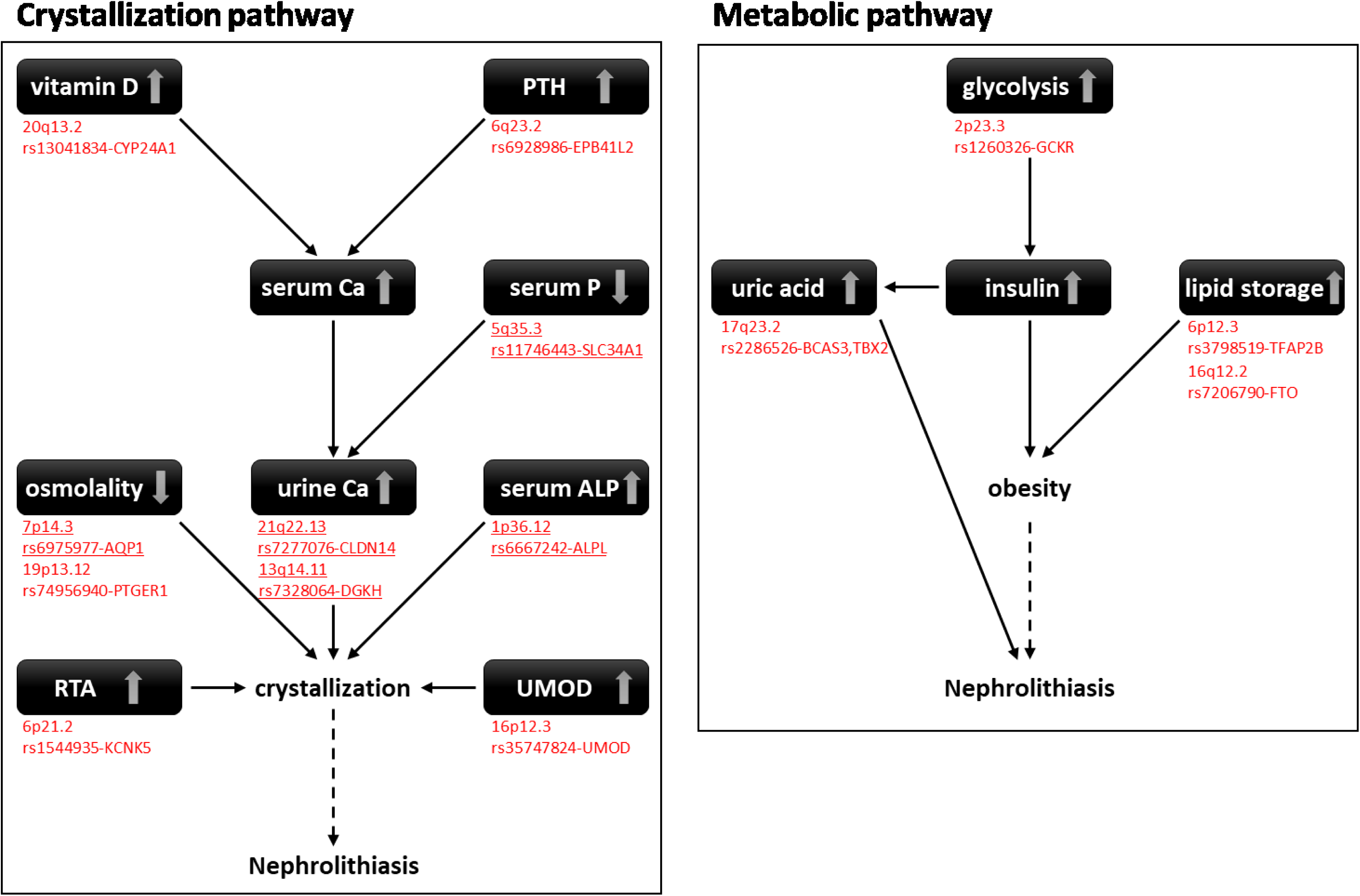
Putative effector genes and functions. Schematic of some established components of the crystallization pathway and metabolic pathway, with the putative effector genes and corresponding lead SNPs. The underlined regions have been reported previously.

Nephrolithiasis is a complex disease that is regulated by many factors, such as serum/urinary cations, urate, kidney function, obesity, humoral factors, dietary factors, and hydration. To further investigate the roles of these genetic factors in the pathogenesis of nephrolithiasis, we analyzed the association of 17 variations with 16 quantitative traits in three categories, including metabolic (n = 7), kidney-related (n = 4), and electrolyte (n = 5) (**Supplemental Table 4**), which were analyzed in our previous studies ^22^. Interestingly, 10 of the 14 regions showed a significant association with any of 16 quantitative traits, including metabolic (2p23.2-3, 6p12.3, and 16q12.2), kidney-related (2p23.3, 5q35.3, 6p12.3, 13q14.11, 16p12.3, 16q12.2, 17q23.2, and 21q22.13), and electrolyte traits (1p36.12, 2p23.3, 5q35.3, 17q23.2, and 20q13.2) (P < 5 × 10^−8^) (**Figure 2**, **Supplemental Figure 4** and **Supplemental Table 5-7**). Among the associations, 2p23.3 was associated with 7 traits across all three categories (TC, TG, BS, sCr, eGFR, UA, and Cl). Moreover, 6p12.3 (BMI and BUN) and 16q12.2 (BMI and BUN) showed pleiotropy across metabolic and kidney-related traits, while 5q35.3 (sCr, eGFR, and P) and 17q23.2 (sCr, eGFR, UA, K, Cl, and Ca) showed pleiotropy across kidney-related traits and electrolytes (**Supplemental Figure 5**). The regional plots of 10 pleiotropic loci are shown in **Supplemental Figure 6**. All 10 regions exhibited a similar pattern of association between the QTLs and nephrolithiasis. The results of a conditioned analysis adjusted by lead SNPs based on our previous QTL analysis ^22^ revealed the colocalization of causal variants between nephrolithiasis and quantitative traits (**Supplemental Figure 6**). We conducted a subgroup analysis using clinical information, such as recurrent stones, stone location, positive family history, and history of gout (**Supplemental Figure 7**). As a result, rs35747824 and rs7277076 were significantly associated with recurrent stones, while rs35747824 and rs13006480 were associated with kidney stones and history of gout.

To evaluate the cumulative effects of genetic variants on nephrolithiasis risk, a weighted-genetic risk score (wGRS) was used. We used the 17 significant SNPs that cleared the GWAS significance threshold in the screening stage and their corresponding weights from the meta-analysis of the screening stage. Individuals were partitioned into quintile groups based on the wGRS. As a result, the highest risk group (top quantile) showed an OR of 2.130 (95% C.I.: 1.999-2.271) and 1.710 (95% C.I.: 1.422-2.059) compared with the reference group (first quantile) in the screening stage and replication stage, respectively (**Supplemental Table 8**). We also evaluated the association of these variations with the mRNA expression of nearby genes. rs1106357, rs1260326, and rs3798519 were associated with ALPL, GCKR, and TFAP2B, respectively (**Supplemental Figure 8**). In addition to rs6667242, rs1106357 was also associated with nephrolithiasis, the serum ALP level, and serum P levels (**Supplemental Figure 9**). These results suggested an underlying molecular mechanism whereby these variations regulate the risk of nephrolithiasis.

## DISCUSSION

We found that nine novel loci were associated with the risk of nephrolithiasis in the Japanese population. To the best of our knowledge, this study is the largest genomic analysis of nephrolithiasis. Carbohydrate intake increases the absorption of calcium in the intestine and the urinary excretion of calcium via insulin secretion ^34 35^. In addition, obesity is associated with higher uric acid in the serum ^36^, a major component of nephrolithiasis ^37^, due to increased uric acid production and decreased renal excretion. Four of nine novel risk loci were associated with higher BMI or triglycerides (2p23.3, 6p12.3, and 16q12.2) and/or higher uric acid content (2p23.3 and 17q23.2) (**Figure 2** and **Supplemental Figure 4**) ^38, 39^, which are well-established risk factors for nephrolithiasis ^5^. *FTO* variations on 16q12.2 suppress the expression of *IRX3* and *IRX5* and subsequently increase body weight by promoting lipid storage ^40^. The risk allele on 6p12.3 was associated with higher expression of *TFAP2B* (**Supplemental Figure 8c**), and TFAP2B promotes the enlargement of adipocytes ^41^. GCKR functions as a negative regulator of GCK, and the risk allele on 2p23.3 would suppress GCKR function via amino acid substitution and/or its transcriptional repression (**Supplemental Figure 8b**) and subsequently enhance the activity of GCK ^42^, which promotes insulin secretion and triglyceride synthesis ^43^ and suppresses uric acid excretion ^44^. Previously, 17q23.2 was shown to be associated with hyperuricemia ^22, 39, 45^, although the underlying molecular mechanism has not been elucidated. Thus, these 4 loci promote the formation of nephrolithiasis through the regulation of metabolic traits, such as obesity and uric acid production (**Figure 3**).

The majority of patients with nephrolithiasis have calcium-containing stones. Four previously reported loci were shown to be associated with higher urine Ca concentration (5q35.3, 7q14.3, 13q14.11, and 21q22.13)^46-48^ and familial nephrolithiasis (*SLC34A1* on 5q35.3) ^49^. Vitamin D and PTH are major regulators of the serum/urine Ca level through the regulation of intestinal Ca absorption, renal excretion/reabsorption, and bone mineralization ^50, 51^. rs13041834 on 20q13.2 is located near *CYP24A1,* which encodes an enzyme that inactivates vitamin D, and its risk allele was associated with higher vitamin D ^52, 53^ and serum Ca ^54^. rs6928986 on 6q23.2 is located within *EPB41L2,* which regulates the localization and activity of PTHR ^55^. rs74956940 on 19p13.12 is located within the *PTGER1* gene that was shown to regulate urine osmotic pressure ^56^. Thus, seven of 14 nephrolithiasis loci were likely to regulate serum/uric electrolyte concentration. Moreover, 1p36.12, 16p12.3 and 6p21.2 would regulate the crystallization step. The risk allele of rs6667242 on 1p36.12 was associated with a high ALP level and low serum P level (**Supplemental Figure 8a and 9**). ALPL hydrates and reduces pyrophosphate and acts as an inhibiter of CaOX and CaP crystallization, subsequently increasing the nephrolithiasis risk ^57, 58^. rs35747824 on 16p12.3 is located within *UMOD*, which encodes UMOD, the most abundant protein in urine that suppresses the crystallization of Ca in urine ^59^. rs1544935 on 6p21.2 is located within *KCNK5*, which encodes a potassium channel. Because KCNK5-knockout mice exhibit distal tubular acidosis and alkali urine that reduces the solubility of calcium phosphate products ^60^, this variation regulates stone formation by regulating urine pH. Taken together, all 14 loci are associated with the regulation of either the metabolic or crystallization pathway (**Figure 3**). We hope our findings will contribute to the elucidation of the molecular pathology of nephrolithiasis and the implementation of personalized medical care for this disease.

## AUTHOR CONTRIBUTIONS

C.Ta., Y.K., M.K., and Ko.M. conceived and planned the experiments. C.Ta., M.U., and Y.Mo. carried out the experiments. C.Ta., Y.K., C.Te., and A.T. carried out statistical analysis. K.S., S.O., A.S., M.S., Ke.M., H.M., M.N., K.W. T.Y., N.S., M.I., S.T., K.K., Y.Y., Y.Mu., M.K., and Ko.M. contributed to sample preparation and data analyasis. C.T., Y.K., and Ko.M. contributed to the interpretation of the results. C.T. and K.M. took the lead in writing the manuscript. All authors provided critical feedback and helped shape the research, analysis and manuscript.

## ACKNOWLEDGMENTS

We thank all the participants in this study. We are grateful for the staff of BioBank Japan, Tohoku Medical Megabank, Iwate Tohoku Medical Megabank, J-MICC, and JPHC for their outstanding assistance. We also thank Satoyo Oda and Yoshiyuki Yukawa for their technical assistance. This study was partially supported by the BioBank Japan project and the Tohoku Medical Megabank project, which is supported by the Ministry of Education, Culture, Sports, Sciences and Technology Japan and the Japan Agency for Medical Research and Development. The JPHC Study was supported by the National Cancer Research and Development Fund since 2010 and was supported by a Grant-in-Aid for Cancer Research from the Ministry of Health, Labour and Welfare of Japan from 1989 to 2010. The J-MICC Study was supported by Grants-in-Aid for Scientific Research for Priority Areas of Cancer (17015018) and Innovative Areas (221S0001) and the JSPS KAKENHI Grant (16H06277) from the Japan Ministry of Education, Science, Sports, Culture and Technology. This study was also supported by JSPS KAKENHI Grants (25293168 to K. M.).

## DISCLOSURES

None

**Supplementary material table of contents**

Supplemental Figure 1. Scheme for study design and screening results

Supplemental Figure 2. Quantile quantile plot for imputed GWAS of nephrolithiasis in the Japanese population

Supplemental Figure 3. Regional plots of 17 loci for nephrolithiasis before and after condition on read SNPs

Supplemental Figure 4. Forest plots for risk variants in the 17 nephrolithiasis risk loci

Supplemental Figure 5. Summary of pleiotropy analysis

Supplemental Figure 6. Regional plots of 10 pleiotropic loci before and after condition

Supplemental Figure 7. Forest plots for risk variants in the 17 nephrolithiasis risk loci

Supplemental Figure 8. eQTL analysis

Supplemental Figure 9. Forest plots for risk variants of rs6667242 and rs1106357

Supplemental Table 1. List of 822 SNPs with p-value of less than 5 × 10^−8^ in meta-analysis of screening 1 and screening 2

Supplemental Table 2. The result of association analysis of kidney stone in the replication stage

Supplemental Table 3. Association of reported SNPs in our GWAS data set

Supplemental Table 4. List of quantitative traits

Supplemental Table 5. Overview of the identified loci and their pleiotropy in metabolic traits

Supplemental Table 6. Overview of the identified loci and their pleiotropy in kidney-related traits

Supplemental Table 7. Overview of the identified loci and their pleiotropy in electrolyte traits

Supplemental Table 8. wGRS analysis of 17 associated SNPs

## REFERENCES

1. Yasui, T, Iguchi, M, Suzuki, S, Kohri, K: Prevalence and epidemiological characteristics of urolithiasis in Japan: national trends between 1965 and 2005. Urology, 71: 209–213, 2008.

2. Scales, CD, Jr., Smith, AC, Hanley, JM, Saigal, CS, Urologic Diseases in America, P: Prevalence of kidney stones in the United States. European urology, 62: 160–165, 2012.

3. Strohmaier, WL: Course of calcium stone disease without treatment. What can we expect? European urology, 37: 339–344, 2000.

4. Scales, CD, Smith, AC, Hanley, JM, Saigal, CS: Prevalence of kidney stones in the United States. Eur Urol, 62: 160–165, 2012.

5. Taylor, EN, Stampfer, MJ, Curhan, GC: Obesity, weight gain, and the risk of kidney stones. JAMA: the journal of the American Medical Association, 293: 455–462, 2005.

6. Borghi, L, Meschi, T, Guerra, A, Briganti, A, Schianchi, T, Allegri, F, Novarini, A: Essential arterial hypertension and stone disease. Kidney international, 55: 2397–2406, 1999.

7. Taylor, EN, Stampfer, MJ, Curhan, GC: Diabetes mellitus and the risk of nephrolithiasis. Kidney international, 68: 1230–1235, 2005.

8. Curhan, GC, Willett, WC, Rimm, EB, Stampfer, MJ: Family history and risk of kidney stones. J Am Soc Nephrol, 8: 1568–1573, 1997.

9. Goldfarb, DS, Fischer, ME, Keich, Y, Goldberg, J: A twin study of genetic and dietary influences on nephrolithiasis: a report from the Vietnam Era Twin (VET) Registry. Kidney Int, 67: 1053–1061, 2005.

10. Botzenhart, E, Vester, U, Schmidt, C, Hesse, A, Halber, M, Wagner, C, Lang, F, Hoyer, P, Zerres, K, Eggermann, T, Arbeitsgemeinschaft fur Padiatrische, N: Cystinuria in children: distribution and frequencies of mutations in the SLC3A1 and SLC7A9 genes. Kidney Int, 62: 1136–1142, 2002.

11. Thorleifsson, G, Holm, H, Edvardsson, V, Walters, GB, Styrkarsdottir, U, Gudbjartsson, DF, Sulem, P, Halldorsson, BV, de Vegt, F, d’Ancona, FC, den Heijer, M, Franzson, L, Christiansen, C, Alexandersen, P, Rafnar, T, Kristjansson, K, Sigurdsson, G, Kiemeney, LA, Bodvarsson, M, Indridason, OS, Palsson, R, Kong, A, Thorsteinsdottir, U, Stefansson, K: Sequence variants in the CLDN14 gene associate with kidney stones and bone mineral density. Nat Genet, 41: 926–930, 2009.

12. Urabe, Y, Tanikawa, C, Takahashi, A, Okada, Y, Morizono, T, Tsunoda, T, Kamatani, N, Kohri, K, Chayama, K, Kubo, M, Nakamura, Y, Matsuda, K: A genome-wide association study of nephrolithiasis in the Japanese population identifies novel susceptible Loci at 5q35.3, 7p14.3, and 13q14.1. PLoS Genet, 8: e1002541, 2012.

13. Oddsson, A, Sulem, P, Helgason, H, Edvardsson, VO, Thorleifsson, G, Sveinbjornsson, G, Haraldsdottir, E, Eyjolfsson, GI, Sigurdardottir, O, Olafsson, I, Masson, G, Holm, H, Gudbjartsson, DF, Thorsteinsdottir, U, Indridason, OS, Palsson, R, Stefansson, K: Common and rare variants associated with kidney stones and biochemical traits. Nature communications, 6: 7975, 2015.

14. Nagai, A, Hirata, M, Kamatani, Y, Muto, K, Matsuda, K, Kiyohara, Y, Ninomiya, T, Tamakoshi, A, Yamagata, Z, Mushiroda, T, Murakami, Y, Yuji, K, Furukawa, Y, Zembutsu, H, Tanaka, T, Ohnishi, Y, Nakamura, Y, BioBank Japan Cooperative Hospital, G, Kubo, M: Overview of the BioBank Japan Project: Study design and profile. J Epidemiol, 2017.

15. Hirata, M, Nagai, A, Kamatani, Y, Ninomiya, T, Tamakoshi, A, Yamagata, Z, Kubo, M, Muto, K, Kiyohara, Y, Mushiroda, T, Murakami, Y, Yuji, K, Furukawa, Y, Zembutsu, H, Tanaka, T, Ohnishi, Y, Nakamura, Y, BioBank Japan Cooperative Hospital, G, Matsuda, K: Overview of BioBank Japan follow-up data in 32 diseases. J Epidemiol, 2017.

16. Tsugane, S, Sobue, T: Baseline survey of JPHC study--design and participation rate. Japan Public Health Center-based Prospective Study on Cancer and Cardiovascular Diseases. J Epidemiol, 11: S24–29, 2001.

17. Hamajima, N, Group, JMS: The Japan Multi-Institutional Collaborative Cohort Study (J-MICC Study) to detect gene-environment interactions for cancer. Asian Pac J Cancer Prev, 8: 317–323, 2007.

18. Kuriyama, S, Yaegashi, N, Nagami, F, Arai, T, Kawaguchi, Y, Osumi, N, Sakaida, M, Suzuki, Y, Nakayama, K, Hashizume, H, Tamiya, G, Kawame, H, Suzuki, K, Hozawa, A, Nakaya, N, Kikuya, M, Metoki, H, Tsuji, I, Fuse, N, Kiyomoto, H, Sugawara, J, Tsuboi, A, Egawa, S, Ito, K, Chida, K, Ishii, T, Tomita, H, Taki, Y, Minegishi, N, Ishii, N, Yasuda, J, Igarashi, K, Shimizu, R, Nagasaki, M, Koshiba, S, Kinoshita, K, Ogishima, S, Takai-Igarashi, T, Tominaga, T, Tanabe, O, Ohuchi, N, Shimosegawa, T, Kure, S, Tanaka, H, Ito, S, Hitomi, J, Tanno, K, Nakamura, M, Ogasawara, K, Kobayashi, S, Sakata, K, Satoh, M, Shimizu, A, Sasaki, M, Endo, R, Sobue, K, Tohoku Medical Megabank Project Study Group, T, Yamamoto, M: The Tohoku Medical Megabank Project: Design and Mission. J Epidemiol, 26: 493–511, 2016.

19. Yasui, T, Okada, A, Urabe, Y, Usami, M, Mizuno, K, Kubota, Y, Tozawa, K, Sasaki, S, Higashi, Y, Sato, Y, Kubo, M, Nakamura, Y, Matsuda, K, Kohri, K: A replication study for three nephrolithiasis loci at 5q35.3, 7p14.3 and 13q14.1 in the Japanese population. J Hum Genet, 58: 588–593, 2013.

20. Akiyama, M, Okada, Y, Kanai, M, Takahashi, A, Momozawa, Y, Ikeda, M, Iwata, N, Ikegawa, S, Hirata, M, Matsuda, K, Iwasaki, M, Yamaji, T, Sawada, N, Hachiya, T, Tanno, K, Shimizu, A, Hozawa, A, Minegishi, N, Tsugane, S, Yamamoto, M, Kubo, M, Kamatani, Y: Genome-wide association study identifies 112 new loci for body mass index in the Japanese population. Nat Genet, 49: 1458–1467, 2017.

21. Ishigaki, K, Akiyama, M, Kanai, M, Takahashi, A, Kawakami, E, Sugishita, H, Sakaue, S, Siew-Kee, L, Okada, Y, Kochi, Y, Horikoshi, M, Ito, K, Momozawa, Y, Hirata, M, Matsuda, K, Ikeda, M, Iwata, N, Ikegawa, S, Kou, I, Tanaka, T, Nakagawa, H, Suzuki, A, Hirota, T, Tamari, M, Chayama, K, Miki, D, Mori, M, Nagayama, S, Daigo, Y, Miki, Y, Katagiri, T, Ogawa, O, Obara, W, Ito, H, Yoshida, T, Imoto, I, Takahashi, T, Tanikawa, C, Minegishi, N, Suzuki, K, Tanno, K, Shimizu, A, Yamaji, T, Iwasaki, M, Sawada, N, Uemura, H, Tanaka, K, Naito, M, Sasaki, M, Wakai, K, Tsugane, S, Yamamoto, M, Yamamoto, K, Murakami, Y, Nakamura, Y, Inazawa, J, Yamauchi, T, Kadowaki, T, Kubo, M, Kamatani, Y: Genetics identifies core genes in the biology of common diseases. under review.

22. Kanai, M, Akiyama, M, Takahashi, A, Matoba, N, Momozawa, Y, Ikeda, M, Iwata, N, Ikegawa, S, Hirata, M, Matsuda, K, Kubo, M, Okada, Y, Kamatani, Y: Genetic analysis of quantitative traits in the Japanese population links cell types to complex human diseases. Nat Genet, 50: 390–400, 2018.

23. Altshuler, DM, Gibbs, RA, Peltonen, L, Altshuler, DM, Gibbs, RA, Peltonen, L, Dermitzakis, E, Schaffner, SF, Yu, F, Peltonen, L, Dermitzakis, E, Bonnen, PE, Altshuler, DM, Gibbs, RA, de Bakker, PI, Deloukas, P, Gabriel, SB, Gwilliam, R, Hunt, S, Inouye, M, Jia, X, Palotie, A, Parkin, M, Whittaker, P, Yu, F, Chang, K, Hawes, A, Lewis, LR, Ren, Y, Wheeler, D, Gibbs, RA, Muzny, DM, Barnes, C, Darvishi, K, Hurles, M, Korn, JM, Kristiansson, K, Lee, C, McCarrol, SA, Nemesh, J, Dermitzakis, E, Keinan, A, Montgomery, SB, Pollack, S, Price, AL, Soranzo, N, Bonnen, PE, Gibbs, RA, Gonzaga-Jauregui, C, Keinan, A, Price, AL, Yu, F, Anttila, V, Brodeur, W, Daly, MJ, Leslie, S, McVean, G, Moutsianas, L, Nguyen, H, Schaffner, SF, Zhang, Q, Ghori, MJ, McGinnis, R, McLaren, W, Pollack, S, Price, AL, Schaffner, SF, Takeuchi, F, Grossman, SR, Shlyakhter, I, Hostetter, EB, Sabeti, PC, Adebamowo, CA, Foster, MW, Gordon, DR, Licinio, J, Manca, MC, Marshall, PA, Matsuda, I, Ngare, D, Wang, VO, Reddy, D, Rotimi, CN, Royal, CD, Sharp, RR, Zeng, C, Brooks, LD, McEwen, JE: Integrating common and rare genetic variation in diverse human populations. Nature, 467: 52–58, 2010.

24. Li, Y, Willer, CJ, Ding, J, Scheet, P, Abecasis, GR: MaCH: using sequence and genotype data to estimate haplotypes and unobserved genotypes. Genet Epidemiol, 34: 816–834, 2010.

25. Auton, A, Brooks, LD, Durbin, RM, Garrison, EP, Kang, HM, Korbel, JO, Marchini, JL, McCarthy, S, McVean, GA, Abecasis, GR: A global reference for human genetic variation. Nature, 526: 68–74, 2015.

26. Willer, CJ, Li, Y, Abecasis, GR: METAL: fast and efficient meta-analysis of genomewide association scans. Bioinformatics, 26: 2190–2191, 2010.

27. Breslow, NE, Day, NE: Statistical methods in cancer research. Volume II--The design and analysis of cohort studies. IARC Sci Publ: 1–406, 1987.

28. Bulik-Sullivan, B, Finucane, HK, Anttila, V, Gusev, A, Day, FR, Loh, PR, Duncan, L, Perry, JR, Patterson, N, Robinson, EB, Daly, MJ, Price, AL, Neale, BM: An atlas of genetic correlations across human diseases and traits. Nat Genet, 47: 1236–1241, 2015.

29. Freedman, ML, Reich, D, Penney, KL, McDonald, GJ, Mignault, AA, Patterson, N, Gabriel, SB, Topol, EJ, Smoller, JW, Pato, CN, Pato, MT, Petryshen, TL, Kolonel, LN, Lander, ES, Sklar, P, Henderson, B, Hirschhorn, JN, Altshuler, D: Assessing the impact of population stratification on genetic association studies. Nat Genet, 36: 388–393, 2004.

30. Yang, J, Weedon, MN, Purcell, S, Lettre, G, Estrada, K, Willer, CJ, Smith, AV, Ingelsson, E, O’Connell, JR, Mangino, M, Magi, R, Madden, PA, Heath, AC, Nyholt, DR, Martin, NG, Montgomery, GW, Frayling, TM, Hirschhorn, JN, McCarthy, MI, Goddard, ME, Visscher, PM: Genomic inflation factors under polygenic inheritance. Eur J Hum Genet, 19: 807–812, 2011.

31. Pruim, RJ, Welch, RP, Sanna, S, Teslovich, TM, Chines, PS, Gliedt, TP, Boehnke, M, Abecasis, GR, Willer, CJ: LocusZoom: regional visualization of genome-wide association scan results. Bioinformatics, 26: 2336–2337, 2010.

32. Bulik-Sullivan, B, Finucane, HK, Anttila, V, Gusev, A, Day, FR, Loh, PR, ReproGen, C, Psychiatric Genomics, C, Genetic Consortium for Anorexia Nervosa of the Wellcome Trust Case Control, C, Duncan, L, Perry, JR, Patterson, N, Robinson, EB, Daly, MJ, Price, AL, Neale, BM: An atlas of genetic correlations across human diseases and traits. Nat Genet, 47: 1236–1241, 2015.

33. Ohnishi, Y, Tanaka, T, Ozaki, K, Yamada, R, Suzuki, H, Nakamura, Y: A high-throughput SNP typing system for genome-wide association studies. Journal of human genetics, 46: 471–477, 2001.

34. Lemann, J, Jr., Piering, WF, Lennon, EJ: Possible role of carbohydrate-induced calciuria in calcium oxalate kidney-stone formation. The New England journal of medicine, 280: 232–237, 1969.

35. Wood, RJ, Allen, LH: Evidence for insulin involvement in arginine- and glucose-induced hypercalciuria in the rat. J Nutr, 113: 1561–1567, 1983.

36. Powell, CR, Stoller, ML, Schwartz, BF, Kane, C, Gentle, DL, Bruce, JE, Leslie, SW: Impact of body weight on urinary electrolytes in urinary stone formers. Urology, 55: 825–830, 2000.

37. Yamashita, S, Matsuzawa, Y, Tokunaga, K, Fujioka, S, Tarui, S: Studies on the impaired metabolism of uric acid in obese subjects: marked reduction of renal urate excretion and its improvement by a low-calorie diet. International journal of obesity, 10: 255–264, 1986.

38. Kolz, M, Johnson, T, Sanna, S, Teumer, A, Vitart, V, Perola, M, Mangino, M, Albrecht, E, Wallace, C, Farrall, M, Johansson, A, Nyholt, DR, Aulchenko, Y, Beckmann, JS, Bergmann, S, Bochud, M, Brown, M, Campbell, H, Connell, J, Dominiczak, A, Homuth, G, Lamina, C, McCarthy, MI, Meitinger, T, Mooser, V, Munroe, P, Nauck, M, Peden, J, Prokisch, H, Salo, P, Salomaa, V, Samani, NJ, Schlessinger, D, Uda, M, Volker, U, Waeber, G, Waterworth, D, Wang-Sattler, R, Wright, AF, Adamski, J, Whitfield, JB, Gyllensten, U, Wilson, JF, Rudan, I, Pramstaller, P, Watkins, H, Doering, A, Wichmann, HE, Spector, TD, Peltonen, L, Volzke, H, Nagaraja, R, Vollenweider, P, Caulfield, M, Illig, T, Gieger, C: Meta-analysis of 28,141 individuals identifies common variants within five new loci that influence uric acid concentrations. PLoS Genet, 5: e1000504, 2009.

39. Kottgen, A, Albrecht, E, Teumer, A, Vitart, V, Krumsiek, J, Hundertmark, C, Pistis, G, Ruggiero, D, O’Seaghdha, CM, Haller, T, Yang, Q, Tanaka, T, Johnson, AD, Kutalik, Z, Smith, AV, Shi, J, Struchalin, M, Middelberg, RP, Brown, MJ, Gaffo, AL, Pirastu, N, Li, G, Hayward, C, Zemunik, T, Huffman, J, Yengo, L, Zhao, JH, Demirkan, A, Feitosa, MF, Liu, X, Malerba, G, Lopez, LM, van der Harst, P, Li, X, Kleber, ME, Hicks, AA, Nolte, IM, Johansson, A, Murgia, F, Wild, SH, Bakker, SJ, Peden, JF, Dehghan, A, Steri, M, Tenesa, A, Lagou, V, Salo, P, Mangino, M, Rose, LM, Lehtimaki, T, Woodward, OM, Okada, Y, Tin, A, Muller, C, Oldmeadow, C, Putku, M, Czamara, D, Kraft, P, Frogheri, L, Thun, GA, Grotevendt, A, Gislason, GK, Harris, TB, Launer, LJ, McArdle, P, Shuldiner, AR, Boerwinkle, E, Coresh, J, Schmidt, H, Schallert, M, Martin, NG, Montgomery, GW, Kubo, M, Nakamura, Y, Tanaka, T, Munroe, PB, Samani, NJ, Jacobs, DR, Jr., Liu, K, D’Adamo, P, Ulivi, S, Rotter, JI, Psaty, BM, Vollenweider, P, Waeber, G, Campbell, S, Devuyst, O, Navarro, P, Kolcic, I, Hastie, N, Balkau, B, Froguel, P, Esko, T, Salumets, A, Khaw, KT, Langenberg, C, Wareham, NJ, Isaacs, A, Kraja, A, Zhang, Q, Wild, PS, Scott, RJ, Holliday, EG, Org, E, Viigimaa, M, Bandinelli, S, Metter, JE, Lupo, A, Trabetti, E, Sorice, R, Doring, A, Lattka, E, Strauch, K, Theis, F, Waldenberger, M, Wichmann, HE, Davies, G, Gow, AJ, Bruinenberg, M, Stolk, RP, Kooner, JS, Zhang, W, Winkelmann, BR, Boehm, BO, Lucae, S, Penninx, BW, Smit, JH, Curhan, G, Mudgal, P, Plenge, RM, Portas, L, Persico, I, Kirin, M, Wilson, JF, Mateo Leach, I, van Gilst, WH, Goel, A, Ongen, H, Hofman, A, Rivadeneira, F, Uitterlinden, AG, Imboden, M, von Eckardstein, A, Cucca, F, Nagaraja, R, Piras, MG, Nauck, M, Schurmann, C, Budde, K, Ernst, F, Farrington, SM, Theodoratou, E, Prokopenko, I, Stumvoll, M, Jula, A, Perola, M, Salomaa, V, Shin, SY, Spector, TD, Sala, C, Ridker, PM, Kahonen, M, Viikari, J, Hengstenberg, C, Nelson, CP, Meschia, JF, Nalls, MA, Sharma, P, Singleton, AB, Kamatani, N, Zeller, T, Burnier, M, Attia, J, Laan, M, Klopp, N, Hillege, HL, Kloiber, S, Choi, H, Pirastu, M, Tore, S, Probst-Hensch, NM, Volzke, H, Gudnason, V, Parsa, A, Schmidt, R, Whitfield, JB, Fornage, M, Gasparini, P, Siscovick, DS, Polasek, O, Campbell, H, Rudan, I, Bouatia-Naji, N, Metspalu, A, Loos, RJ, van Duijn, CM, Borecki, IB, Ferrucci, L, Gambaro, G, Deary, IJ, Wolffenbuttel, BH, Chambers, JC, Marz, W, Pramstaller, PP, Snieder, H, Gyllensten, U, Wright, AF, Navis, G, Watkins, H, Witteman, JC, Sanna, S, Schipf, S, Dunlop, MG, Tonjes, A, Ripatti, S, Soranzo, N, Toniolo, D, Chasman, DI, Raitakari, O, Kao, WH, Ciullo, M, Fox, CS, Caulfield, M, Bochud, M, Gieger, C: Genome-wide association analyses identify 18 new loci associated with serum urate concentrations. Nat Genet, 45: 145–154, 2013.

40. Claussnitzer, M, Dankel, SN, Kim, KH, Quon, G, Meuleman, W, Haugen, C, Glunk, V, Sousa, IS, Beaudry, JL, Puviindran, V, Abdennur, NA, Liu, J, Svensson, PA, Hsu, YH, Drucker, DJ, Mellgren, G, Hui, CC, Hauner, H, Kellis, M: FTO Obesity Variant Circuitry and Adipocyte Browning in Humans. The New England journal of medicine, 373: 895–907, 2015.

41. Tao, Y, Maegawa, H, Ugi, S, Ikeda, K, Nagai, Y, Egawa, K, Nakamura, T, Tsukada, S, Nishio, Y, Maeda, S, Kashiwagi, A: The transcription factor AP-2beta causes cell enlargement and insulin resistance in 3T3-L1 adipocytes. Endocrinology, 147: 1685–1696, 2006.

42. Rees, MG, Wincovitch, S, Schultz, J, Waterstradt, R, Beer, NL, Baltrusch, S, Collins, FS, Gloyn, AL: Cellular characterisation of the GCKR P446L variant associated with type 2 diabetes risk. Diabetologia, 55: 114–122, 2012.

43. Beer, NL, Tribble, ND, McCulloch, LJ, Roos, C, Johnson, PR, Orho-Melander, M, Gloyn, AL: The P446L variant in GCKR associated with fasting plasma glucose and triglyceride levels exerts its effect through increased glucokinase activity in liver. Hum Mol Genet, 18: 4081– 4088, 2009.

44. Ter Maaten, JC, Voorburg, A, Heine, RJ, Ter Wee, PM, Donker, AJ, Gans, RO: Renal handling of urate and sodium during acute physiological hyperinsulinaemia in healthy subjects. Clinical science (London, England: 1979), 92: 51–58, 1997.

45. Li, C, Li, Z, Liu, S, Wang, C, Han, L, Cui, L, Zhou, J, Zou, H, Liu, Z, Chen, J, Cheng, X, Zhou, Z, Ding, C, Wang, M, Chen, T, Cui, Y, He, H, Zhang, K, Yin, C, Wang, Y, Xing, S, Li, B, Ji, J, Jia, Z, Ma, L, Niu, J, Xin, Y, Liu, T, Chu, N, Yu, Q, Ren, W, Wang, X, Zhang, A, Sun, Y, Wang, H, Lu, J, Li, Y, Qing, Y, Chen, G, Wang, Y, Zhou, L, Niu, H, Liang, J, Dong, Q, Li, X, Mi, QS, Shi, Y: Genome-wide association analysis identifies three new risk loci for gout arthritis in Han Chinese. Nature communications, 6: 7041, 2015.

46. Corre, T, Olinger, E, Harris, SE, Traglia, M, Ulivi, S, Lenarduzzi, S, Belge, H, Youhanna, S, Tokonami, N, Bonny, O, Houillier, P, Polasek, O, Deary, IJ, Starr, JM, Toniolo, D, Gasparini, P, Vollenweider, P, Hayward, C, Bochud, M, Devuyst, O: Common variants in CLDN14 are associated with differential excretion of magnesium over calcium in urine. Pflugers Archiv: European journal of physiology, 469: 91–103, 2017.

47. Xu, Y, Zeng, G, Mai, Z, Ou, L: Association study of DGKH gene polymorphisms with calcium oxalate stone in Chinese population. Urolithiasis, 42: 379–385, 2014.

48. Wang, L, Feng, C, Ding, G, Lin, X, Gao, P, Jiang, H, Xu, J, Ding, Q, Wu, Z: Association Study of Reported Significant Loci at 5q35.3, 7p14.3, 13q14.1 and 16p12.3 with Urolithiasis in Chinese Han Ethnicity. Scientific reports, 7: 45766, 2017.

49. Prie, D, Huart, V, Bakouh, N, Planelles, G, Dellis, O, Gerard, B, Hulin, P, Benque-Blanchet, F, Silve, C, Grandchamp, B, Friedlander, G: Nephrolithiasis and osteoporosis associated with hypophosphatemia caused by mutations in the type 2a sodium-phosphate cotransporter. The New England journal of medicine, 347: 983–991, 2002.

50. Moe, SM: Disorders involving calcium, phosphorus, and magnesium. Prim Care, 35: 215-237, v-vi, 2008.

51. Weiser, MM, Bloor, JH, Dasmahapatra, A: Intestinal calcium absorption and vitamin D metabolism. J Clin Gastroenterol, 4: 75–86, 1982.

52. Manousaki, D, Dudding, T, Haworth, S, Hsu, YH, Liu, CT, Medina-Gomez, C, Voortman, T, van der Velde, N, Melhus, H, Robinson-Cohen, C, Cousminer, DL, Nethander, M, Vandenput, L, Noordam, R, Forgetta, V, Greenwood, CMT, Biggs, ML, Psaty, BM, Rotter, JI, Zemel, BS, Mitchell, JA, Taylor, B, Lorentzon, M, Karlsson, M, Jaddoe, VVW, Tiemeier, H, Campos-Obando, N, Franco, OH, Utterlinden, AG, Broer, L, van Schoor, NM, Ham, AC, Ikram, MA, Karasik, D, de Mutsert, R, Rosendaal, FR, den Heijer, M, Wang, TJ, Lind, L, Orwoll, ES, Mook-Kanamori, DO, Michaelsson, K, Kestenbaum, B, Ohlsson, C, Mellstrom, D, de Groot, L, Grant, SFA, Kiel, DP, Zillikens, MC, Rivadeneira, F, Sawcer, S, Timpson, NJ, Richards, JB: Low-Frequency Synonymous Coding Variation in CYP2R1 Has Large Effects on Vitamin D Levels and Risk of Multiple Sclerosis. American journal of human genetics, 101: 227–238, 2017.

53. Jiang, X, O’Reilly, PF, Aschard, H, Hsu, YH, Richards, JB, Dupuis, J, Ingelsson, E, Karasik, D, Pilz, S, Berry, D, Kestenbaum, B, Zheng, J, Luan, J, Sofianopoulou, E, Streeten, EA, Albanes, D, Lutsey, PL, Yao, L, Tang, W, Econs, MJ, Wallaschofski, H, Volzke, H, Zhou, A, Power, C, McCarthy, MI, Michos, ED, Boerwinkle, E, Weinstein, SJ, Freedman, ND, Huang, WY, Van Schoor, NM, van der Velde, N, Groot, L, Enneman, A, Cupples, LA, Booth, SL, Vasan, RS, Liu, CT, Zhou, Y, Ripatti, S, Ohlsson, C, Vandenput, L, Lorentzon, M, Eriksson, JG, Shea, MK, Houston, DK, Kritchevsky, SB, Liu, Y, Lohman, KK, Ferrucci, L, Peacock, M, Gieger, C, Beekman, M, Slagboom, E, Deelen, J, Heemst, DV, Kleber, ME, Marz, W, de Boer, IH, Wood, AC, Rotter, JI, Rich, SS, Robinson-Cohen, C, den Heijer, M, Jarvelin, MR, Cavadino, A, Joshi, PK, Wilson, JF, Hayward, C, Lind, L, Michaelsson, K, Trompet, S, Zillikens, MC, Uitterlinden, AG, Rivadeneira, F, Broer, L, Zgaga, L, Campbell, H, Theodoratou, E, Farrington, SM, Timofeeva, M, Dunlop, MG, Valdes, AM, Tikkanen, E, Lehtimaki, T, Lyytikainen, LP, Kahonen, M, Raitakari, OT, Mikkila, V, Ikram, MA, Sattar, N, Jukema, JW, Wareham, NJ, Langenberg, C, Forouhi, NG, Gundersen, TE, Khaw, KT, Butterworth, AS, Danesh, J, Spector, T, Wang, TJ, Hypponen, E, Kraft, P, Kiel, DP: Genome-wide association study in 79,366 European-ancestry individuals informs the genetic architecture of 25-hydroxyvitamin D levels. Nature communications, 9: 260, 2018.

54. O’Seaghdha, CM, Wu, H, Yang, Q, Kapur, K, Guessous, I, Zuber, AM, Kottgen, A, Stoudmann, C, Teumer, A, Kutalik, Z, Mangino, M, Dehghan, A, Zhang, W, Eiriksdottir, G, Li, G, Tanaka, T, Portas, L, Lopez, LM, Hayward, C, Lohman, K, Matsuda, K, Padmanabhan, S, Firsov, D, Sorice, R, Ulivi, S, Brockhaus, AC, Kleber, ME, Mahajan, A, Ernst, FD, Gudnason, V, Launer, LJ, Mace, A, Boerwinckle, E, Arking, DE, Tanikawa, C, Nakamura, Y, Brown, MJ, Gaspoz, JM, Theler, JM, Siscovick, DS, Psaty, BM, Bergmann, S, Vollenweider, P, Vitart, V, Wright, AF, Zemunik, T, Boban, M, Kolcic, I, Navarro, P, Brown, EM, Estrada, K, Ding, J, Harris, TB, Bandinelli, S, Hernandez, D, Singleton, AB, Girotto, G, Ruggiero, D, d’Adamo, AP, Robino, A, Meitinger, T, Meisinger, C, Davies, G, Starr, JM, Chambers, JC, Boehm, BO, Winkelmann, BR, Huang, J, Murgia, F, Wild, SH, Campbell, H, Morris, AP, Franco, OH, Hofman, A, Uitterlinden, AG, Rivadeneira, F, Volker, U, Hannemann, A, Biffar, R, Hoffmann, W, Shin, SY, Lescuyer, P, Henry, H, Schurmann, C, Munroe, PB, Gasparini, P, Pirastu, N, Ciullo, M, Gieger, C, Marz, W, Lind, L, Spector, TD, Smith, AV, Rudan, I, Wilson, JF, Polasek, O, Deary, IJ, Pirastu, M, Ferrucci, L, Liu, Y, Kestenbaum, B, Kooner, JS, Witteman, JC, Nauck, M, Kao, WH, Wallaschofski, H, Bonny, O, Fox, CS, Bochud, M: Meta-analysis of genome-wide association studies identifies six new Loci for serum calcium concentrations. PLoS Genet, 9: e1003796, 2013.

55. Saito, M, Sugai, M, Katsushima, Y, Yanagisawa, T, Sukegawa, J, Nakahata, N: Increase in cell-surface localization of parathyroid hormone receptor by cytoskeletal protein 4.1G. The Biochemical journal, 392: 75–81, 2005.

56. Kennedy, CR, Xiong, H, Rahal, S, Vanderluit, J, Slack, RS, Zhang, Y, Guan, Y, Breyer, MD, Hebert, RL: Urine concentrating defect in prostaglandin EP1-deficient mice. American journal of physiology Renal physiology, 292: F868–875, 2007.

57. Moochhala, SH, Sayer, JA, Carr, G, Simmons, NL: Renal calcium stones: insights from the control of bone mineralization. Experimental physiology, 93: 43–49, 2008.

58. Basavaraj, DR, Biyani, CS, Browning, AJ, Cartledge, JJ: The Role of Urinary Kidney Stone Inhibitors and Promoters in the Pathogenesis of Calcium Containing Renal Stones. EAU-EBU Update Series, 5: 126–136, 2007.

59. Chen, WC, Lin, HS, Chen, HY, Shih, CH, Li, CW: Effects of Tamm-Horsfall protein and albumin on calcium oxalate crystallization and importance of sialic acids. Molecular urology, 5: 1–5, 2001.

60. Wagner, CA, Mohebbi, N: Urinary pH and stone formation. Journal of nephrology, 23 Suppl 16: S165–169, 2010.

