## Supplementary data1 for "GWAS identifies nine nephrolithiasis susceptibility loci related with metabolic metabolic and crystallization pathways"

**Supplemental Table 1. List of 822 SNPs with p-value of less than 5 x 10-8 in meta-analysis of screening 1 and screening 2**

| MarkerName | CHR | POS | Allele1 | Allele2 | Effect | StdErr | P-value | Direction[a] | HetISq[b] | HetChiSq[c] | HetDf[d] | HetPVal[e] | num_of_gene | gene | relativeloc | gene | relativeloc |
|---|---|---|---|---|---|---|---|---|---|---|---|---|---|---|---|---|---|
| rs3820297 | 1 | 21758355 | t | c | 0.159 | 0.028 | 8.14E-09 | ++ | 0 | 0.258 | 1 | 0.612 | 1 | HS6ST1P1 | 0 | | |
| rs10916988 | 1 | 21759020 | a | g | -0.128 | 0.021 | 1.85E-09 | -- | 0 | 0.245 | 1 | 0.621 | 1 | HS6ST1P1 | 293 | | |
| rs10916989 | 1 | 21759128 | a | g | -0.107 | 0.019 | 3.39E-08 | -- | 0 | 0.169 | 1 | 0.681 | 1 | HS6ST1P1 | 401 | | |
| rs12759170 | 1 | 21759839 | t | g | 0.121 | 0.020 | 1.16E-09 | ++ | 0 | 0.184 | 1 | 0.668 | 1 | LOC100421114 | -965 | | |
| rs12734589 | 1 | 21760127 | a | g | -0.117 | 0.019 | 8.89E-10 | -- | 0 | 0.176 | 1 | 0.675 | 1 | LOC100421114 | -677 | | |
| rs4079441 | 1 | 21761309 | t | g | -0.104 | 0.018 | 1.54E-08 | -- | 0 | 0.145 | 1 | 0.703 | 1 | LOC100421114 | 0 | | |
| rs10916990 | 1 | 21763132 | a | t | -0.116 | 0.019 | 1.23E-09 | -- | 0 | 0.176 | 1 | 0.675 | 1 | LOC100421114 | 0 | | |
| rs6680628 | 1 | 21765211 | a | g | 0.104 | 0.018 | 1.81E-08 | ++ | 0 | 0.166 | 1 | 0.683 | 1 | LOC100652932 | 1371 | | |
| rs6668141 | 1 | 21765268 | c | g | 0.110 | 0.019 | 9.73E-09 | ++ | 0 | 0.056 | 1 | 0.814 | 1 | LOC100652932 | 1314 | | |
| rs6694671 | 1 | 21765925 | t | g | -0.120 | 0.020 | 1.58E-09 | -- | 0 | 0.052 | 1 | 0.820 | 1 | LOC100652932 | 657 | | |
| rs1566523 | 1 | 21766663 | c | g | -0.116 | 0.019 | 1.45E-09 | -- | 0 | 0.154 | 1 | 0.694 | 2 | LOC100652932 | 0 | NBPF3 | 0 |
| rs4654745 | 1 | 21767096 | t | g | -0.119 | 0.020 | 2.16E-09 | -- | 0 | 0.184 | 1 | 0.668 | 2 | LOC100652932 | 0 | NBPF3 | 0 |
| rs12047493 | 1 | 21768191 | c | g | -0.118 | 0.019 | 1.37E-09 | -- | 0 | 0.149 | 1 | 0.700 | 2 | LOC100652932 | 0 | NBPF3 | 0 |
| rs10916993 | 1 | 21770035 | t | c | -0.124 | 0.021 | 1.48E-09 | -- | 0 | 0.193 | 1 | 0.661 | 2 | LOC100652932 | 0 | NBPF3 | 0 |
| rs10799691 | 1 | 21773362 | c | g | 0.125 | 0.020 | 4.87E-10 | ++ | 32.3 | 1.478 | 1 | 0.224 | 1 | NBPF3 | 0 | | |
| rs10799692 | 1 | 21775943 | t | c | -0.107 | 0.018 | 6.92E-09 | -- | 0 | 0.089 | 1 | 0.765 | 1 | NBPF3 | 0 | | |
| rs10737456 | 1 | 21776121 | a | g | -0.110 | 0.019 | 9.73E-09 | -- | 0 | 0.083 | 1 | 0.773 | 1 | NBPF3 | 0 | | |
| rs12565322 | 1 | 21776335 | a | c | 0.103 | 0.018 | 2.11E-08 | ++ | 0 | 0.074 | 1 | 0.786 | 1 | NBPF3 | 0 | | |
| rs10737457 | 1 | 21776392 | a | c | 0.106 | 0.018 | 8.14E-09 | ++ | 0 | 0.074 | 1 | 0.786 | 1 | NBPF3 | 0 | | |
| rs6685071 | 1 | 21776884 | a | g | 0.106 | 0.018 | 8.14E-09 | ++ | 0 | 0.074 | 1 | 0.786 | 1 | NBPF3 | 0 | | |
| rs4654747 | 1 | 21777426 | a | c | 0.098 | 0.018 | 2.96E-08 | ++ | 0 | 0.157 | 1 | 0.692 | 1 | NBPF3 | 0 | | |
| rs4654936 | 1 | 21777452 | a | g | 0.107 | 0.018 | 5.88E-09 | ++ | 0 | 0.074 | 1 | 0.786 | 1 | NBPF3 | 0 | | |
| rs4311854 | 1 | 21777751 | a | g | 0.098 | 0.018 | 2.96E-08 | ++ | 0 | 0.157 | 1 | 0.692 | 1 | NBPF3 | 0 | | |
| rs10737458 | 1 | 21777960 | t | c | -0.108 | 0.018 | 4.24E-09 | -- | 0 | 0.189 | 1 | 0.664 | 1 | NBPF3 | 0 | | |
| rs4654937 | 1 | 21778035 | a | g | 0.097 | 0.018 | 4.09E-08 | ++ | 0 | 0.157 | 1 | 0.692 | 1 | NBPF3 | 0 | | |
| rs1814736 | 1 | 21778204 | a | t | -0.098 | 0.018 | 2.96E-08 | -- | 0 | 0.115 | 1 | 0.734 | 1 | NBPF3 | 0 | | |
| rs2016464 | 1 | 21778965 | t | c | -0.108 | 0.019 | 1.54E-08 | -- | 0 | 0.044 | 1 | 0.834 | 1 | NBPF3 | 0 | | |
| rs4399126 | 1 | 21779090 | t | c | 0.101 | 0.018 | 1.96E-08 | ++ | 0 | 0.077 | 1 | 0.782 | 1 | NBPF3 | 0 | | |
| rs2016413 | 1 | 21779465 | a | g | -0.104 | 0.018 | 8.40E-09 | -- | 0 | 0.077 | 1 | 0.782 | 1 | NBPF3 | 0 | | |
| rs2016203 | 1 | 21779737 | t | c | -0.107 | 0.019 | 1.18E-08 | -- | 0 | 0.071 | 1 | 0.790 | 1 | NBPF3 | 0 | | |
| rs2010202 | 1 | 21780651 | a | g | 0.106 | 0.018 | 8.14E-09 | ++ | 0 | 0.074 | 1 | 0.786 | 1 | NBPF3 | 0 | | |
| rs10737459 | 1 | 21780834 | t | c | -0.102 | 0.018 | 7.93E-09 | -- | 0 | 0.051 | 1 | 0.821 | 1 | NBPF3 | 0 | | |
| rs4414015 | 1 | 21784060 | c | g | -0.097 | 0.017 | 1.09E-08 | -- | 0 | 0.087 | 1 | 0.768 | 1 | NBPF3 | 0 | | |
| rs4417036 | 1 | 21784069 | t | g | 0.099 | 0.018 | 2.52E-08 | ++ | 0 | 0.097 | 1 | 0.756 | 1 | NBPF3 | 0 | | |
| rs4495665 | 1 | 21784070 | a | t | -0.098 | 0.018 | 2.96E-08 | -- | 0 | 0.115 | 1 | 0.734 | 1 | NBPF3 | 0 | | |
| rs4515756 | 1 | 21784093 | a | g | -0.092 | 0.016 | 1.54E-08 | -- | 0 | 0.185 | 1 | 0.667 | 1 | NBPF3 | 0 | | |
| rs4385671 | 1 | 21784299 | c | g | 0.092 | 0.016 | 1.54E-08 | ++ | 0 | 0.185 | 1 | 0.667 | 1 | NBPF3 | 0 | | |
| rs6675223 | 1 | 21785229 | a | g | -0.092 | 0.016 | 1.54E-08 | -- | 0 | 0.185 | 1 | 0.667 | 1 | NBPF3 | 0 | | |
| rs6666997 | 1 | 21785287 | t | c | 0.092 | 0.016 | 1.54E-08 | ++ | 0 | 0.185 | 1 | 0.667 | 1 | NBPF3 | 0 | | |
| rs4393124 | 1 | 21785489 | a | c | -0.096 | 0.017 | 1.54E-08 | -- | 0 | 0.087 | 1 | 0.768 | 1 | NBPF3 | 0 | | |
| rs4430298 | 1 | 21785540 | a | g | -0.096 | 0.017 | 6.88E-09 | -- | 0 | 0.090 | 1 | 0.764 | 1 | NBPF3 | 0 | | |
| rs4654938 | 1 | 21785981 | t | g | -0.092 | 0.016 | 1.84E-08 | -- | 0 | 0.160 | 1 | 0.689 | 2 | NBPF3 | 0 | PFN1P10 | 0 |
| rs4654748 | 1 | 21786068 | t | c | -0.104 | 0.017 | 8.89E-10 | -- | 0 | 0.170 | 1 | 0.680 | 2 | NBPF3 | 0 | PFN1P10 | 0 |
| rs4654750 | 1 | 21786251 | t | c | 0.096 | 0.017 | 6.88E-09 | ++ | 0 | 0.090 | 1 | 0.764 | 2 | NBPF3 | 0 | PFN1P10 | 0 |
| rs12738545 | 1 | 21786750 | a | g | 0.091 | 0.016 | 2.63E-08 | ++ | 0 | 0.213 | 1 | 0.645 | 1 | NBPF3 | 0 | | |
| rs2316509 | 1 | 21787018 | a | g | 0.092 | 0.016 | 1.84E-08 | ++ | 0 | 0.160 | 1 | 0.689 | 1 | NBPF3 | 0 | | |
| rs4654939 | 1 | 21787731 | a | c | 0.092 | 0.016 | 1.84E-08 | ++ | 0 | 0.160 | 1 | 0.689 | 1 | NBPF3 | 0 | | |
| rs6660003 | 1 | 21789442 | a | g | 0.096 | 0.017 | 3.14E-08 | ++ | 0 | 0.187 | 1 | 0.665 | 1 | NBPF3 | 0 | | |
| rs2316505 | 1 | 21793958 | t | c | -0.094 | 0.016 | 8.97E-09 | -- | 0 | 0.160 | 1 | 0.689 | 1 | NBPF3 | 0 | | |
| rs7541779 | 1 | 21794612 | t | g | -0.093 | 0.017 | 2.24E-08 | -- | 0 | 0.262 | 1 | 0.609 | 1 | NBPF3 | 0 | | |
| rs1827293 | 1 | 21795388 | a | g | 0.090 | 0.015 | 1.67E-09 | ++ | 0 | 0.409 | 1 | 0.522 | 1 | NBPF3 | 0 | | |
| rs4654752 | 1 | 21795824 | t | c | 0.092 | 0.016 | 1.54E-08 | ++ | 0 | 0.185 | 1 | 0.667 | 1 | NBPF3 | 0 | | |
| rs7537914 | 1 | 21796616 | a | g | 0.092 | 0.016 | 1.54E-08 | ++ | 0 | 0.185 | 1 | 0.667 | 1 | NBPF3 | 0 | | |
| rs6687836 | 1 | 21796770 | a | c | -0.092 | 0.015 | 5.81E-10 | -- | 0 | 0.454 | 1 | 0.501 | 1 | NBPF3 | 0 | | |
| rs4654944 | 1 | 21797419 | t | c | 0.092 | 0.016 | 1.54E-08 | ++ | 0 | 0.185 | 1 | 0.667 | 1 | NBPF3 | 0 | | |
| rs2004380 | 1 | 21797996 | a | g | -0.090 | 0.015 | 1.35E-09 | -- | 0 | 0.454 | 1 | 0.501 | 1 | NBPF3 | 0 | | |
| rs7514120 | 1 | 21798866 | a | g | 0.092 | 0.016 | 1.84E-08 | ++ | 0 | 0.213 | 1 | 0.645 | 1 | NBPF3 | 0 | | |
| rs1971327 | 1 | 21799489 | t | c | 0.092 | 0.016 | 1.84E-08 | ++ | 0 | 0.213 | 1 | 0.645 | 1 | NBPF3 | 0 | | |
| rs7520015 | 1 | 21800185 | t | g | 0.092 | 0.016 | 1.84E-08 | ++ | 0 | 0.213 | 1 | 0.645 | 1 | NBPF3 | 0 | | |
| rs10916996 | 1 | 21800219 | t | c | -0.092 | 0.016 | 1.84E-08 | -- | 0 | 0.213 | 1 | 0.645 | 1 | NBPF3 | 0 | | |
| rs10799694 | 1 | 21800436 | c | g | 0.092 | 0.016 | 1.84E-08 | ++ | 0 | 0.213 | 1 | 0.645 | 1 | NBPF3 | 0 | | |
| rs7554122 | 1 | 21801116 | t | c | -0.092 | 0.016 | 1.84E-08 | -- | 0 | 0.213 | 1 | 0.645 | 1 | NBPF3 | 0 | | |
| rs7531217 | 1 | 21801346 | a | g | -0.092 | 0.016 | 1.84E-08 | -- | 0 | 0.213 | 1 | 0.645 | 1 | NBPF3 | 0 | | |
| rs4654753 | 1 | 21801838 | t | c | 0.092 | 0.016 | 1.84E-08 | ++ | 0 | 0.213 | 1 | 0.645 | 1 | NBPF3 | 0 | | |
| rs4654945 | 1 | 21802029 | t | c | -0.092 | 0.016 | 1.84E-08 | -- | 0 | 0.213 | 1 | 0.645 | 1 | NBPF3 | 0 | | |
| rs4654946 | 1 | 21802122 | t | c | 0.092 | 0.016 | 1.54E-08 | ++ | 0 | 0.242 | 1 | 0.623 | 1 | NBPF3 | 0 | | |
| rs6657015 | 1 | 21802450 | t | c | -0.091 | 0.016 | 2.63E-08 | -- | 0 | 0.213 | 1 | 0.645 | 1 | NBPF3 | 0 | | |
| rs6669637 | 1 | 21802453 | a | g | 0.091 | 0.016 | 2.63E-08 | ++ | 0 | 0.213 | 1 | 0.645 | 1 | NBPF3 | 0 | | |
| rs6426713 | 1 | 21803177 | a | g | 0.091 | 0.015 | 8.89E-10 | ++ | 0 | 0.454 | 1 | 0.501 | 1 | NBPF3 | 0 | | |
| rs7527656 | 1 | 21803664 | a | g | 0.092 | 0.016 | 1.84E-08 | ++ | 0 | 0.213 | 1 | 0.645 | 1 | NBPF3 | 0 | | |
| rs7527672 | 1 | 21803734 | a | g | 0.092 | 0.016 | 1.84E-08 | ++ | 0 | 0.213 | 1 | 0.645 | 1 | NBPF3 | 0 | | |
| rs4654947 | 1 | 21805048 | t | c | -0.092 | 0.016 | 1.84E-08 | -- | 0 | 0.213 | 1 | 0.645 | 1 | NBPF3 | 0 | | |
| rs2316504 | 1 | 21805730 | a | g | 0.095 | 0.017 | 1.13E-08 | ++ | 0 | 0.204 | 1 | 0.652 | 1 | NBPF3 | 0 | | |
| rs4654948 | 1 | 21805978 | a | g | -0.094 | 0.016 | 8.97E-09 | -- | 0 | 0.213 | 1 | 0.645 | 1 | NBPF3 | 0 | | |
| rs3820296 | 1 | 21806447 | t | c | 0.100 | 0.016 | 9.48E-10 | ++ | 0 | 0.417 | 1 | 0.519 | 1 | NBPF3 | 0 | | |
| rs12043777 | 1 | 21806667 | c | g | 0.127 | 0.021 | 9.94E-10 | ++ | 0 | 0.331 | 1 | 0.565 | 1 | NBPF3 | 0 | | |
| rs12034222 | 1 | 21806710 | t | g | 0.114 | 0.019 | 2.77E-09 | ++ | 0 | 0.363 | 1 | 0.547 | 1 | NBPF3 | 0 | | |
| rs56286426 | 1 | 21806792 | t | c | -0.136 | 0.023 | 1.85E-09 | -- | 0 | 0.195 | 1 | 0.659 | 1 | NBPF3 | 0 | | |
| rs1106357 | 1 | 21817085 | t | c | -0.101 | 0.016 | 2.03E-10 | -- | 0 | 0.720 | 1 | 0.396 | 1 | NBPF3 | 5692 | | |
| rs72659148 | 1 | 21817397 | a | g | 0.106 | 0.017 | 5.08E-10 | ++ | 0 | 0.001 | 1 | 0.977 | 1 | NBPF3 | 6004 | | |
| rs7555005 | 1 | 21819611 | a | g | 0.102 | 0.017 | 1.85E-09 | ++ | 0 | 0.087 | 1 | 0.768 | 1 | NBPF3 | 8218 | | |
| rs1697421 | 1 | 21823292 | t | c | -0.090 | 0.015 | 1.67E-09 | -- | 0 | 0.954 | 1 | 0.329 | 1 | NBPF3 | 11899 | | |
| rs1697420 | 1 | 21823629 | t | c | -0.104 | 0.015 | 3.17E-12 | -- | 60.1 | 2.505 | 1 | 0.114 | 1 | ALPL | -12229 | | |
| rs6667242 | 1 | 21826566 | a | g | -0.098 | 0.015 | 4.12E-11 | -- | 0 | 0.367 | 1 | 0.545 | 1 | ALPL | -9292 | | |
| rs56155961 | 1 | 21827094 | t | c | 0.098 | 0.015 | 4.12E-11 | ++ | 0 | 0.367 | 1 | 0.545 | 1 | ALPL | -8764 | | |
| rs55805990 | 1 | 21827179 | t | c | 0.098 | 0.015 | 4.12E-11 | ++ | 0 | 0.367 | 1 | 0.545 | 1 | ALPL | -8679 | | |
| rs1780328 | 1 | 21827321 | c | g | -0.092 | 0.016 | 1.84E-08 | -- | 0 | 0.273 | 1 | 0.601 | 1 | ALPL | -8537 | | |
| rs12566622 | 1 | 21828619 | a | g | -0.116 | 0.018 | 1.10E-10 | -- | 0 | 0.151 | 1 | 0.698 | 1 | ALPL | -7239 | | |
| rs1936823 | 1 | 21829452 | t | c | -0.101 | 0.016 | 8.44E-11 | -- | 0 | 0.413 | 1 | 0.520 | 1 | ALPL | -6406 | | |
| rs1936824 | 1 | 21829615 | a | g | 0.100 | 0.016 | 1.29E-10 | ++ | 0 | 0.335 | 1 | 0.563 | 1 | ALPL | -6243 | | |
| rs56702468 | 1 | 21830340 | t | c | -0.091 | 0.016 | 2.20E-08 | -- | 0 | 0.306 | 1 | 0.580 | 1 | ALPL | -5518 | | |
| rs13353032 | 1 | 21830659 | t | g | -0.098 | 0.015 | 4.12E-11 | -- | 0 | 0.367 | 1 | 0.545 | 1 | ALPL | -5199 | | |
| rs13353029 | 1 | 21830686 | a | g | -0.098 | 0.015 | 4.12E-11 | -- | 0 | 0.367 | 1 | 0.545 | 1 | ALPL | -5172 | | |
| rs7520712 | 1 | 21831163 | a | g | -0.098 | 0.015 | 4.12E-11 | -- | 0 | 0.367 | 1 | 0.545 | 1 | ALPL | -4695 | | |
| rs34828194 | 1 | 21831494 | t | c | 0.092 | 0.016 | 1.54E-08 | ++ | 0 | 0.378 | 1 | 0.539 | 1 | ALPL | -4364 | | |
| rs3856178 | 1 | 21832730 | t | c | 0.098 | 0.015 | 4.12E-11 | ++ | 0 | 0.367 | 1 | 0.545 | 1 | ALPL | -3128 | | |
| rs72659157 | 1 | 21833457 | c | g | -0.091 | 0.016 | 2.20E-08 | -- | 0 | 0.242 | 1 | 0.623 | 1 | ALPL | -2401 | | |
| rs72659158 | 1 | 21833509 | t | c | 0.098 | 0.015 | 4.12E-11 | ++ | 0 | 0.367 | 1 | 0.545 | 1 | ALPL | -2349 | | |
| rs12092961 | 1 | 21833856 | t | c | -0.091 | 0.016 | 2.20E-08 | -- | 0 | 0.242 | 1 | 0.623 | 1 | ALPL | -2002 | | |
| rs12066162 | 1 | 21833925 | t | c | 0.099 | 0.015 | 3.28E-11 | ++ | 0 | 0.409 | 1 | 0.522 | 1 | ALPL | -1933 | | |
| rs77362499 | 1 | 21836204 | c | g | -0.098 | 0.015 | 5.17E-11 | -- | 0 | 0.409 | 1 | 0.522 | 1 | ALPL | 0 | | |
| rs10917002 | 1 | 21836340 | t | c | 0.098 | 0.015 | 5.17E-11 | ++ | 0 | 0.409 | 1 | 0.522 | 1 | ALPL | 0 | | |
| rs6703976 | 1 | 21836934 | t | c | 0.098 | 0.015 | 5.17E-11 | ++ | 0 | 0.409 | 1 | 0.522 | 1 | ALPL | 0 | | |
| rs16825455 | 1 | 21837755 | a | g | -0.098 | 0.015 | 5.17E-11 | -- | 0 | 0.409 | 1 | 0.522 | 1 | ALPL | 0 | | |
| rs1472563 | 1 | 21840498 | t | c | 0.085 | 0.016 | 4.66E-08 | ++ | 0 | 0.103 | 1 | 0.748 | 1 | ALPL | 0 | | |
| rs7521331 | 1 | 21840649 | t | c | -0.086 | 0.016 | 3.23E-08 | -- | 0 | 0.103 | 1 | 0.748 | 1 | ALPL | 0 | | |
| rs7535497 | 1 | 21840709 | t | g | 0.086 | 0.016 | 3.23E-08 | ++ | 0 | 0.103 | 1 | 0.748 | 1 | ALPL | 0 | | |
| rs7523855 | 1 | 21840886 | t | c | -0.086 | 0.016 | 3.23E-08 | -- | 0 | 0.103 | 1 | 0.748 | 1 | ALPL | 0 | | |

| | | | | | | | | | | | | | | | |
|---|---|---|---|---|---|---|---|---|---|---|---|---|---|---|---|
| rs7538283 | 1 | 21841344 | t | c | 0.086 | 0.016 | 3.23E-08 | ++ | 0 | 0.103 | 1 | 0.748 | 1 | ALPL | 0 |
| rs1780317 | 1 | 21841974 | t | c | 0.087 | 0.016 | 2.69E-08 | ++ | 0 | 0.084 | 1 | 0.772 | 1 | ALPL | 0 |
| rs1780314 | 1 | 21842877 | t | c | 0.087 | 0.016 | 2.69E-08 | ++ | 0 | 0.084 | 1 | 0.772 | 1 | ALPL | 0 |
| rs4654755 | 1 | 21843608 | t | c | 0.087 | 0.016 | 2.69E-08 | ++ | 0 | 0.084 | 1 | 0.772 | 1 | ALPL | 0 |
| rs4654956 | 1 | 21844297 | a | g | 0.088 | 0.016 | 3.46E-08 | ++ | 0 | 0.080 | 1 | 0.777 | 1 | ALPL | 0 |
| rs7536545 | 1 | 21844822 | t | c | -0.088 | 0.016 | 3.46E-08 | -- | 0 | 0.080 | 1 | 0.777 | 1 | ALPL | 0 |
| rs10917003 | 1 | 21845550 | t | c | -0.089 | 0.016 | 4.44E-08 | -- | 0 | 0.060 | 1 | 0.806 | 1 | ALPL | 0 |
| rs1260326 | 2 | 27730940 | t | c | 0.098 | 0.015 | 5.17E-11 | ++ | 19 | 1.235 | 1 | 0.267 | 1 | GCKR | 0 |
| rs6547692 | 2 | 27734972 | a | g | -0.093 | 0.015 | 3.78E-10 | -- | 31.9 | 1.469 | 1 | 0.225 | 1 | GCKR | 0 |
| rs780096 | 2 | 27741072 | c | g | 0.093 | 0.015 | 3.78E-10 | ++ | 38.9 | 1.637 | 1 | 0.201 | 1 | GCKR | 0 |
| rs780095 | 2 | 27741105 | a | g | 0.093 | 0.015 | 4.69E-10 | ++ | 42 | 1.724 | 1 | 0.189 | 1 | GCKR | 0 |
| rs780094 | 2 | 27741237 | t | c | 0.093 | 0.015 | 4.69E-10 | ++ | 47.5 | 1.906 | 1 | 0.167 | 1 | GCKR | 0 |
| rs780093 | 2 | 27742603 | t | c | 0.095 | 0.015 | 1.58E-10 | ++ | 50 | 2.000 | 1 | 0.157 | 1 | GCKR | 0 |
| rs1260334 | 2 | 27748597 | a | c | -0.098 | 0.016 | 6.12E-10 | -- | 61.1 | 2.568 | 1 | 0.109 | 1 | GCKR | 2046 |
| rs1260333 | 2 | 27748624 | a | g | 0.089 | 0.015 | 5.47E-09 | ++ | 39.2 | 1.644 | 1 | 0.200 | 1 | GCKR | 2073 |
| rs1313566 | 2 | 27748904 | a | g | -0.096 | 0.015 | 1.01E-10 | -- | 54.4 | 2.195 | 1 | 0.139 | 1 | GCKR | 2353 |
| rs2950835 | 2 | 27750545 | a | g | 0.093 | 0.017 | 2.09E-08 | ++ | 42.9 | 1.752 | 1 | 0.186 | 1 | GCKR | 3994 |
| rs2911711 | 2 | 27750546 | a | t | -0.093 | 0.017 | 2.09E-08 | -- | 42.9 | 1.752 | 1 | 0.186 | 1 | GCKR | 3995 |
| rs11127048 | 2 | 27752463 | a | g | -0.102 | 0.017 | 4.62E-09 | -- | 53.8 | 2.166 | 1 | 0.141 | 1 | GCKR | 5912 |
| rs6753534 | 2 | 27752871 | t | c | -0.097 | 0.016 | 2.97E-09 | -- | 55.9 | 2.269 | 1 | 0.132 | 1 | GCKR | 6320 |
| rs7606605 | 2 | 27755716 | a | t | -0.131 | 0.023 | 7.06E-09 | -- | 68 | 3.125 | 1 | 0.077 | 1 | GCKR | 9165 |
| rs59210625 | 2 | 27769474 | a | c | -0.131 | 0.023 | 8.05E-09 | -- | 67.2 | 3.047 | 1 | 0.081 | 1 | GCKR | 22923 |
| rs3749147 | 2 | 27851918 | a | g | 0.090 | 0.016 | 8.75E-09 | ++ | 68 | 3.125 | 1 | 0.077 | 1 | GPN1 | 0 |
| rs13002853 | 2 | 27853245 | c | g | -0.089 | 0.016 | 4.44E-08 | -- | 63.7 | 2.756 | 1 | 0.097 | 1 | GPN1 | 0 |
| rs34502053 | 2 | 27854524 | a | g | 0.090 | 0.016 | 8.75E-09 | ++ | 68 | 3.125 | 1 | 0.077 | 1 | GPN1 | 0 |
| rs10173720 | 2 | 27855397 | a | t | 0.094 | 0.016 | 1.85E-09 | ++ | 21 | 1.265 | 1 | 0.261 | 1 | GPN1 | 0 |
| rs6719175 | 2 | 27859351 | a | t | 0.089 | 0.016 | 1.28E-08 | ++ | 65.5 | 2.902 | 1 | 0.088 | 1 | GPN1 | 0 |
| rs2141371 | 2 | 27860258 | a | g | 0.102 | 0.017 | 9.66E-10 | ++ | 45.4 | 1.833 | 1 | 0.176 | 1 | GPN1 | 0 |
| rs77350364 | 2 | 27866139 | a | g | 0.093 | 0.015 | 4.69E-10 | ++ | 28 | 1.389 | 1 | 0.239 | 1 | GPN1 | 0 |
| rs12474585 | 2 | 27871593 | t | c | -0.092 | 0.015 | 1.33E-09 | -- | 35.9 | 1.561 | 1 | 0.212 | 1 | GPN1 | 0 |
| rs8731 | 2 | 27873326 | c | g | 0.091 | 0.015 | 8.89E-10 | ++ | 31.9 | 1.469 | 1 | 0.225 | 1 | GPN1 | 0 |
| rs4666010 | 2 | 27878686 | t | c | 0.090 | 0.015 | 1.35E-09 | ++ | 38.9 | 1.637 | 1 | 0.201 | 1 | SUPT7L | 0 |
| rs2272406 | 2 | 27892023 | a | t | -0.090 | 0.015 | 1.67E-09 | -- | 66.1 | 2.949 | 1 | 0.086 | 1 | SLC4A1AP | 0 |
| rs76586169 | 2 | 27897835 | a | g | -0.091 | 0.015 | 1.10E-09 | -- | 35.6 | 1.552 | 1 | 0.213 | 1 | SLC4A1AP | 0 |
| rs13021208 | 2 | 27901728 | t | c | 0.090 | 0.015 | 1.67E-09 | ++ | 66.1 | 2.949 | 1 | 0.086 | 1 | SLC4A1AP | 0 |
| rs72613829 | 2 | 27902987 | t | c | -0.091 | 0.015 | 1.10E-09 | -- | 35.6 | 1.552 | 1 | 0.213 | 1 | SLC4A1AP | 0 |
| rs13023094 | 2 | 27910706 | a | c | -0.090 | 0.015 | 1.67E-09 | -- | 66.1 | 2.949 | 1 | 0.086 | 1 | SLC4A1AP | 0 |
| rs12998770 | 2 | 27913188 | t | c | -0.090 | 0.015 | 1.67E-09 | -- | 66.1 | 2.949 | 1 | 0.086 | 1 | SLC4A1AP | 0 |
| rs13030973 | 2 | 27928797 | t | c | -0.092 | 0.015 | 5.81E-10 | -- | 67.4 | 3.066 | 1 | 0.080 | 1 | SLC4A1AP | 10950 |
| rs4665385 | 2 | 27930837 | t | c | -0.093 | 0.015 | 4.69E-10 | -- | 35.6 | 1.552 | 1 | 0.213 | 1 | SLC4A1AP | 12990 |
| rs6727388 | 2 | 27932587 | a | g | -0.092 | 0.015 | 5.81E-10 | -- | 67.4 | 3.066 | 1 | 0.080 | 1 | SLC4A1AP | 14740 |
| rs4616435 | 2 | 27933642 | t | c | 0.093 | 0.015 | 3.78E-10 | ++ | 67.4 | 3.066 | 1 | 0.080 | 1 | SLC4A1AP | 15795 |
| rs58199976 | 2 | 27934524 | t | c | 0.092 | 0.015 | 5.81E-10 | ++ | 38.9 | 1.637 | 1 | 0.201 | 1 | SLC4A1AP | 16677 |
| rs6727215 | 2 | 27934731 | a | g | 0.094 | 0.015 | 3.04E-10 | ++ | 68.6 | 3.185 | 1 | 0.074 | 1 | SLC4A1AP | 16884 |
| rs4665386 | 2 | 27935352 | a | c | -0.093 | 0.015 | 4.69E-10 | -- | 35.6 | 1.552 | 1 | 0.213 | 1 | SLC4A1AP | 17505 |
| rs115386260 | 2 | 27939071 | t | c | -0.097 | 0.017 | 4.58E-09 | -- | 81.8 | 5.506 | 1 | 0.019 | 1 | SLC4A1AP | 21224 |
| rs6742571 | 2 | 27941888 | a | g | -0.092 | 0.015 | 5.81E-10 | -- | 38.9 | 1.637 | 1 | 0.201 | 1 | SLC4A1AP | 24041 |
| rs1881394 | 2 | 27949052 | a | g | -0.092 | 0.015 | 7.19E-10 | -- | 35.6 | 1.552 | 1 | 0.213 | 1 | SLC4A1AP | 31205 |
| rs10432705 | 2 | 27950837 | a | g | 0.094 | 0.015 | 6.91E-10 | ++ | 70.5 | 3.390 | 1 | 0.066 | 1 | SLC4A1AP | 32990 |
| rs867282 | 2 | 27951658 | t | c | -0.091 | 0.015 | 8.89E-10 | -- | 38.9 | 1.637 | 1 | 0.201 | 1 | SLC4A1AP | 33811 |
| rs2384655 | 2 | 27952235 | a | t | -0.091 | 0.015 | 8.89E-10 | -- | 38.9 | 1.637 | 1 | 0.201 | 1 | SLC4A1AP | 34388 |
| rs12994085 | 2 | 27952406 | t | c | 0.091 | 0.015 | 8.89E-10 | ++ | 38.9 | 1.637 | 1 | 0.201 | 1 | SLC4A1AP | 34559 |
| rs12993377 | 2 | 27952767 | c | g | 0.091 | 0.015 | 8.89E-10 | ++ | 38.9 | 1.637 | 1 | 0.201 | 1 | SLC4A1AP | 34920 |
| rs12624148 | 2 | 27956481 | t | c | -0.093 | 0.015 | 4.69E-10 | -- | 42 | 1.724 | 1 | 0.189 | 1 | MRPL33 | -38103 |
| rs6753736 | 2 | 27958855 | c | g | -0.093 | 0.015 | 3.78E-10 | -- | 44.9 | 1.814 | 1 | 0.178 | 1 | MRPL33 | -35729 |
| rs1317580 | 2 | 27961344 | c | g | 0.108 | 0.016 | 3.12E-11 | ++ | 72.5 | 3.633 | 1 | 0.057 | 1 | MRPL33 | -33240 |
| rs75429814 | 2 | 27961614 | t | c | 0.094 | 0.015 | 2.45E-10 | ++ | 50 | 2.000 | 1 | 0.157 | 1 | MRPL33 | -32970 |
| rs111938697 | 2 | 27963133 | t | c | -0.094 | 0.015 | 2.45E-10 | -- | 50 | 2.000 | 1 | 0.157 | 1 | MRPL33 | -31451 |
| rs12714223 | 2 | 27963330 | a | g | 0.094 | 0.015 | 2.45E-10 | ++ | 50 | 2.000 | 1 | 0.157 | 1 | MRPL33 | -31254 |
| rs9678824 | 2 | 27964328 | c | g | -0.094 | 0.015 | 2.45E-10 | -- | 50 | 2.000 | 1 | 0.157 | 1 | MRPL33 | -30256 |
| rs13023194 | 2 | 27967260 | c | g | 0.104 | 0.016 | 5.94E-11 | ++ | 76 | 4.171 | 1 | 0.041 | 1 | MRPL33 | -27324 |
| rs9967838 | 2 | 27968454 | a | c | -0.094 | 0.015 | 3.04E-10 | -- | 52.3 | 2.096 | 1 | 0.148 | 1 | MRPL33 | -26130 |
| rs9677663 | 2 | 27972445 | a | g | 0.092 | 0.015 | 5.81E-10 | ++ | 50 | 2.000 | 1 | 0.157 | 1 | MRPL33 | -22139 |
| rs57238885 | 2 | 27974260 | c | g | 0.092 | 0.015 | 5.81E-10 | ++ | 50 | 2.000 | 1 | 0.157 | 1 | MRPL33 | -20324 |
| rs10201247 | 2 | 27974971 | a | g | -0.092 | 0.015 | 5.81E-10 | -- | 50 | 2.000 | 1 | 0.157 | 1 | MRPL33 | -19613 |
| rs6547796 | 2 | 27975394 | t | c | -0.092 | 0.015 | 5.81E-10 | -- | 50 | 2.000 | 1 | 0.157 | 1 | MRPL33 | -19190 |
| rs183183485 | 2 | 27980616 | a | c | -0.107 | 0.018 | 1.70E-09 | -- | 43.4 | 1.767 | 1 | 0.184 | 1 | MRPL33 | -13968 |
| rs4666011 | 2 | 27980912 | c | g | -0.091 | 0.015 | 8.89E-10 | -- | 44.9 | 1.814 | 1 | 0.178 | 1 | MRPL33 | -13672 |
| rs10209020 | 2 | 27980944 | t | c | 0.091 | 0.015 | 8.89E-10 | ++ | 44.9 | 1.814 | 1 | 0.178 | 1 | MRPL33 | -13640 |
| rs12471347 | 2 | 27981655 | c | g | -0.091 | 0.015 | 8.89E-10 | -- | 44.9 | 1.814 | 1 | 0.178 | 1 | MRPL33 | -12929 |
| rs77154381 | 2 | 27982119 | a | g | 0.091 | 0.015 | 8.89E-10 | ++ | 44.9 | 1.814 | 1 | 0.178 | 1 | MRPL33 | -12465 |
| rs12477908 | 2 | 27987046 | a | c | -0.091 | 0.015 | 8.89E-10 | -- | 44.9 | 1.814 | 1 | 0.178 | 1 | MRPL33 | -7538 |
| rs13006480 | 2 | 27987221 | c | g | -0.104 | 0.016 | 2.30E-11 | -- | 76.4 | 4.231 | 1 | 0.040 | 1 | MRPL33 | -7363 |
| rs13013484 | 2 | 27988821 | a | g | 0.091 | 0.015 | 8.89E-10 | ++ | 44.9 | 1.814 | 1 | 0.178 | 1 | MRPL33 | -5763 |
| rs4665390 | 2 | 27989909 | t | c | -0.091 | 0.015 | 8.89E-10 | -- | 44.9 | 1.814 | 1 | 0.178 | 1 | MRPL33 | -4675 |
| rs74429612 | 2 | 27993843 | t | c | -0.091 | 0.015 | 8.89E-10 | -- | 44.9 | 1.814 | 1 | 0.178 | 1 | MRPL33 | -741 |
| rs3736594 | 2 | 27995781 | a | c | -0.091 | 0.015 | 8.89E-10 | -- | 44.9 | 1.814 | 1 | 0.178 | 1 | MRPL33 | 0 |
| rs3792252 | 2 | 27995931 | a | g | 0.091 | 0.015 | 8.89E-10 | ++ | 44.9 | 1.814 | 1 | 0.178 | 1 | MRPL33 | 0 |
| rs35463245 | 2 | 28000724 | c | g | 0.091 | 0.015 | 8.89E-10 | ++ | 44.9 | 1.814 | 1 | 0.178 | 1 | MRPL33 | 0 |
| rs17006242 | 2 | 28001051 | a | c | -0.091 | 0.015 | 8.89E-10 | -- | 44.9 | 1.814 | 1 | 0.178 | 1 | MRPL33 | 0 |
| rs3792253 | 2 | 28002002 | t | c | 0.092 | 0.015 | 7.19E-10 | ++ | 47.5 | 1.906 | 1 | 0.167 | 1 | MRPL33 | 0 |
| rs13030345 | 2 | 28003174 | t | g | 0.112 | 0.017 | 5.03E-11 | ++ | 69 | 3.230 | 1 | 0.072 | 1 | MRPL33 | 566 |
| rs74327441 | 2 | 28007072 | a | g | -0.092 | 0.015 | 7.19E-10 | -- | 47.5 | 1.906 | 1 | 0.167 | 1 | RBKS | 0 |
| rs7593136 | 2 | 28017792 | a | g | 0.094 | 0.015 | 2.45E-10 | ++ | 61.7 | 2.612 | 1 | 0.106 | 1 | RBKS | 0 |
| rs6744348 | 2 | 28018025 | a | c | 0.094 | 0.015 | 2.45E-10 | ++ | 61.7 | 2.612 | 1 | 0.106 | 1 | RBKS | 0 |
| rs36035825 | 2 | 28018131 | t | g | 0.096 | 0.015 | 1.01E-10 | ++ | 67.4 | 3.066 | 1 | 0.080 | 1 | RBKS | 0 |
| rs4666012 | 2 | 28018293 | a | g | 0.094 | 0.015 | 2.45E-10 | ++ | 61.7 | 2.612 | 1 | 0.106 | 1 | RBKS | 0 |
| rs4666013 | 2 | 28019106 | t | c | -0.094 | 0.015 | 2.45E-10 | -- | 61.7 | 2.612 | 1 | 0.106 | 1 | RBKS | 0 |
| rs4666014 | 2 | 28019175 | a | g | -0.094 | 0.015 | 2.45E-10 | -- | 61.7 | 2.612 | 1 | 0.106 | 1 | RBKS | 0 |
| rs4666015 | 2 | 28019367 | a | g | -0.094 | 0.015 | 2.45E-10 | -- | 61.7 | 2.612 | 1 | 0.106 | 1 | RBKS | 0 |
| rs34181670 | 2 | 28019856 | t | c | 0.094 | 0.015 | 2.45E-10 | ++ | 61.7 | 2.612 | 1 | 0.106 | 1 | RBKS | 0 |
| rs35679500 | 2 | 28019966 | t | c | 0.096 | 0.015 | 1.01E-10 | ++ | 61.7 | 2.612 | 1 | 0.106 | 1 | RBKS | 0 |
| rs12623170 | 2 | 28020157 | t | c | -0.094 | 0.015 | 2.45E-10 | -- | 61.7 | 2.612 | 1 | 0.106 | 1 | RBKS | 0 |
| rs3935148 | 2 | 28020966 | a | t | -0.095 | 0.015 | 1.58E-10 | -- | 61.7 | 2.612 | 1 | 0.106 | 1 | RBKS | 0 |
| rs13027427 | 2 | 28021043 | t | g | -0.092 | 0.016 | 3.34E-09 | -- | 0 | 0.930 | 1 | 0.335 | 1 | RBKS | 0 |
| rs6745545 | 2 | 28021845 | c | g | 0.092 | 0.016 | 3.34E-09 | ++ | 0 | 0.930 | 1 | 0.335 | 1 | RBKS | 0 |
| rs7570730 | 2 | 28022675 | a | g | -0.092 | 0.016 | 3.34E-09 | -- | 0 | 0.930 | 1 | 0.335 | 1 | RBKS | 0 |
| rs11127125 | 2 | 28023120 | t | c | 0.092 | 0.016 | 3.34E-09 | ++ | 0 | 0.930 | 1 | 0.335 | 1 | RBKS | 0 |
| rs11127126 | 2 | 28023284 | t | c | -0.092 | 0.016 | 4.06E-09 | -- | 0 | 0.993 | 1 | 0.319 | 1 | RBKS | 0 |
| rs12473340 | 2 | 28023797 | a | g | -0.092 | 0.016 | 4.06E-09 | -- | 0 | 0.993 | 1 | 0.319 | 1 | RBKS | 0 |
| rs12473400 | 2 | 28024024 | t | g | -0.092 | 0.016 | 4.06E-09 | -- | 0 | 0.993 | 1 | 0.319 | 1 | RBKS | 0 |
| rs4630723 | 2 | 28024331 | t | c | 0.095 | 0.016 | 5.18E-09 | ++ | 0 | 0.968 | 1 | 0.325 | 1 | RBKS | 0 |
| rs13019037 | 2 | 28024642 | t | c | -0.092 | 0.016 | 4.06E-09 | -- | 0 | 0.993 | 1 | 0.319 | 1 | RBKS | 0 |
| rs4632296 | 2 | 28024984 | t | c | 0.091 | 0.016 | 5.97E-09 | ++ | 0 | 0.993 | 1 | 0.319 | 1 | RBKS | 0 |
| rs13001060 | 2 | 28025061 | t | c | 0.091 | 0.016 | 5.97E-09 | ++ | 0 | 0.993 | 1 | 0.319 | 1 | RBKS | 0 |
| rs12995865 | 2 | 28025066 | a | c | 0.091 | 0.016 | 5.97E-09 | ++ | 0 | 0.993 | 1 | 0.319 | 1 | RBKS | 0 |
| rs11891769 | 2 | 28025580 | c | g | -0.091 | 0.016 | 5.97E-09 | -- | 0 | 0.993 | 1 | 0.319 | 1 | RBKS | 0 |
| rs13153019 | 5 | 176782218 | t | c | -0.136 | 0.017 | 1.41E-15 | -- | 66.9 | 3.022 | 1 | 0.082 | 1 | RGS14 | -2626 |
| rs4976688 | 5 | 176784439 | t | c | -0.133 | 0.016 | 3.73E-16 | -- | 69.6 | 3.290 | 1 | 0.070 | 1 | RGS14 | -405 |
| rs4976689 | 5 | 176784448 | c | g | -0.101 | 0.015 | 2.62E-11 | -- | 67.1 | 3.037 | 1 | 0.081 | 1 | RGS14 | -396 |
| rs4075958 | 5 | 176784512 | a | g | 0.133 | 0.016 | 3.73E-16 | ++ | 69.6 | 3.290 | 1 | 0.070 | 1 | RGS14 | -332 |

| | | | | | | | | | | | | | | | | | |
|---|---|---|---|---|---|---|---|---|---|---|---|---|---|---|---|---|---|
| rs4075959 | 5 | 176784612 | a | g | 0.134 | 0.016 | 2.24E-16 | ++ | 69.6 | 3.290 | 1 | 0.070 | 1 | RGS14 | -232 | | |
| rs4976646 | 5 | 176788570 | t | c | -0.103 | 0.015 | 1.32E-11 | -- | 65.8 | 2.923 | 1 | 0.087 | 1 | RGS14 | 0 | | |
| rs4976647 | 5 | 176788622 | a | c | -0.103 | 0.015 | 1.06E-11 | -- | 67.1 | 3.037 | 1 | 0.081 | 1 | RGS14 | 0 | | |
| rs67111717 | 5 | 176790162 | a | g | -0.105 | 0.016 | 1.48E-11 | -- | 66.8 | 3.012 | 1 | 0.083 | 1 | RGS14 | 0 | | |
| rs6556313 | 5 | 176792491 | a | g | -0.106 | 0.015 | 1.21E-12 | -- | 66.1 | 2.949 | 1 | 0.086 | 1 | RGS14 | 0 | | |
| rs11135015 | 5 | 176792557 | t | c | -0.106 | 0.015 | 1.21E-12 | -- | 66.1 | 2.949 | 1 | 0.086 | 1 | RGS14 | 0 | | |
| rs11741640 | 5 | 176792743 | a | g | 0.136 | 0.016 | 6.15E-17 | ++ | 68.5 | 3.180 | 1 | 0.075 | 1 | RGS14 | 0 | | |
| rs12654812 | 5 | 176794191 | a | g | 0.105 | 0.015 | 1.54E-12 | ++ | 64.7 | 2.834 | 1 | 0.092 | 1 | RGS14 | 0 | | |
| rs4074995 | 5 | 176797343 | a | g | 0.136 | 0.016 | 7.98E-17 | ++ | 69.6 | 3.290 | 1 | 0.070 | 1 | RGS14 | 0 | | |
| rs56235845 | 5 | 176798040 | t | g | -0.128 | 0.017 | 1.14E-14 | -- | 68.3 | 3.150 | 1 | 0.076 | 1 | RGS14 | 0 | | |
| rs11746443 | 5 | 176798306 | a | g | 0.136 | 0.016 | 7.98E-17 | ++ | 69.6 | 3.290 | 1 | 0.070 | 1 | RGS14 | 0 | | |
| rs10051765 | 5 | 176799992 | t | c | -0.107 | 0.015 | 2.02E-12 | -- | 63 | 2.703 | 1 | 0.100 | 1 | RGS14 | 393 | | |
| rs11748165 | 5 | 176800054 | t | c | 0.136 | 0.016 | 6.15E-17 | ++ | 68.5 | 3.180 | 1 | 0.075 | 1 | RGS14 | 455 | | |
| rs11748297 | 5 | 176800361 | a | g | 0.136 | 0.016 | 6.15E-17 | ++ | 68.5 | 3.180 | 1 | 0.075 | 1 | RGS14 | 762 | | |
| rs10866705 | 5 | 176801131 | a | c | -0.133 | 0.016 | 1.63E-17 | -- | 62.8 | 2.687 | 1 | 0.101 | 1 | RGS14 | 1532 | | |
| rs79098771 | 5 | 176803733 | a | g | 0.142 | 0.021 | 5.19E-12 | ++ | 83.5 | 6.065 | 1 | 0.014 | 1 | RGS14 | 4134 | | |
| rs35716097 | 5 | 176806636 | t | c | 0.110 | 0.016 | 1.94E-12 | ++ | 59.7 | 2.480 | 1 | 0.115 | 1 | SLC34A1 | -4796 | | |
| rs12659266 | 5 | 176807197 | t | c | 0.135 | 0.017 | 3.57E-16 | ++ | 60.7 | 2.542 | 1 | 0.111 | 1 | SLC34A1 | -4235 | | |
| rs3812036 | 5 | 176813404 | t | c | 0.136 | 0.017 | 2.12E-16 | ++ | 57.5 | 2.354 | 1 | 0.125 | 1 | SLC34A1 | 0 | | |
| rs33921462 | 5 | 176814656 | a | g | 0.129 | 0.016 | 1.11E-16 | ++ | 0 | 0.930 | 1 | 0.335 | 1 | SLC34A1 | 0 | | |
| rs5030873 | 5 | 176815124 | t | c | -0.134 | 0.016 | 2.24E-16 | -- | 0 | 0.908 | 1 | 0.341 | 1 | SLC34A1 | 0 | | |
| rs35610898 | 5 | 176815766 | c | g | 0.130 | 0.016 | 8.46E-17 | ++ | 0 | 0.869 | 1 | 0.351 | 1 | SLC34A1 | 0 | | |
| rs56297224 | 5 | 176815880 | a | g | 0.130 | 0.016 | 8.46E-17 | ++ | 0 | 0.869 | 1 | 0.351 | 1 | SLC34A1 | 0 | | |
| rs3812035 | 5 | 176817143 | t | g | 0.128 | 0.016 | 1.90E-16 | ++ | 0 | 0.930 | 1 | 0.335 | 1 | SLC34A1 | 0 | | |
| rs55785724 | 5 | 176817583 | a | t | -0.128 | 0.016 | 1.90E-16 | -- | 0 | 0.930 | 1 | 0.335 | 1 | SLC34A1 | 0 | | |
| rs6420094 | 5 | 176817636 | a | g | -0.128 | 0.016 | 1.90E-16 | -- | 0 | 0.930 | 1 | 0.335 | 1 | SLC34A1 | 0 | | |
| rs6556314 | 5 | 176819252 | t | c | 0.129 | 0.016 | 1.45E-16 | ++ | 0 | 0.869 | 1 | 0.351 | 1 | SLC34A1 | 0 | | |
| rs6556316 | 5 | 176821375 | t | c | 0.129 | 0.016 | 1.45E-16 | ++ | 0 | 0.869 | 1 | 0.351 | 1 | SLC34A1 | 0 | | |
| rs6420095 | 5 | 176821449 | a | c | -0.129 | 0.016 | 1.45E-16 | -- | 0 | 0.869 | 1 | 0.351 | 1 | SLC34A1 | 0 | | |
| rs6862195 | 5 | 176822512 | t | g | 0.129 | 0.016 | 1.45E-16 | ++ | 0 | 0.869 | 1 | 0.351 | 1 | SLC34A1 | 0 | | |
| rs7447593 | 5 | 176824137 | c | g | -0.131 | 0.016 | 4.91E-17 | -- | 0 | 0.869 | 1 | 0.351 | 1 | SLC34A1 | 0 | | |
| rs4976691 | 5 | 176827815 | c | g | -0.130 | 0.016 | 2.99E-16 | -- | 0 | 0.617 | 1 | 0.432 | 1 | PFN3 | -178 | | |
| rs17876032 | 5 | 176830627 | a | g | 0.133 | 0.016 | 3.73E-16 | ++ | 0 | 0.417 | 1 | 0.519 | 1 | F12 | 0 | | |
| rs17876031 | 5 | 176831119 | a | g | -0.134 | 0.016 | 2.24E-16 | -- | 0 | 0.273 | 1 | 0.601 | 1 | F12 | 0 | | |
| rs4976649 | 5 | 176833415 | a | g | 0.115 | 0.019 | 7.82E-10 | ++ | 0 | 0.046 | 1 | 0.831 | 1 | F12 | 0 | | |
| rs228815 | 6 | 39106945 | t | g | 0.093 | 0.017 | 4.25E-08 | ++ | 0 | 0.889 | 1 | 0.346 | 1 | C6orf64 | -24080 | | |
| rs228816 | 6 | 39108006 | c | g | -0.093 | 0.017 | 4.25E-08 | -- | 0 | 0.889 | 1 | 0.346 | 1 | C6orf64 | -25141 | | |
| rs228817 | 6 | 39108973 | t | c | -0.093 | 0.017 | 4.25E-08 | -- | 0 | 0.889 | 1 | 0.346 | 1 | C6orf64 | -26108 | | |
| rs1544935 | 6 | 39124448 | t | g | -0.118 | 0.019 | 7.53E-10 | -- | 60.8 | 2.552 | 1 | 0.110 | 1 | KCNK5 | 32299 | | |
| rs2894421 | 6 | 39128098 | t | g | 0.118 | 0.019 | 7.53E-10 | ++ | 63.3 | 2.722 | 1 | 0.099 | 1 | KCNK5 | 28649 | | |
| rs56336142 | 6 | 39134099 | t | c | -0.107 | 0.019 | 2.09E-08 | -- | 50 | 2.000 | 1 | 0.157 | 1 | KCNK5 | 22648 | | |
| rs1155347 | 6 | 39146230 | t | c | -0.113 | 0.020 | 1.33E-08 | -- | 68.9 | 3.215 | 1 | 0.073 | 1 | KCNK5 | 10517 | | |
| rs74388152 | 6 | 50702485 | a | g | 0.114 | 0.018 | 6.68E-10 | ++ | 0 | 0.214 | 1 | 0.644 | 1 | TFAP2D | 0 | | |
| rs2744475 | 6 | 50784880 | c | g | -0.098 | 0.015 | 1.06E-10 | -- | 80.6 | 5.147 | 1 | 0.023 | 1 | TFAP2B | -1559 | | |
| rs2206271 | 6 | 50786008 | a | t | 0.100 | 0.015 | 5.55E-11 | ++ | 80 | 4.999 | 1 | 0.025 | 1 | TFAP2B | -431 | | |
| rs62405419 | 6 | 50787459 | t | g | 0.117 | 0.016 | 2.03E-13 | ++ | 21.8 | 1.279 | 1 | 0.258 | 1 | TFAP2B | 0 | | |
| rs3798519 | 6 | 50788778 | a | c | -0.116 | 0.016 | 2.60E-13 | -- | 17.3 | 1.209 | 1 | 0.272 | 1 | TFAP2B | 0 | | |
| rs2076308 | 6 | 50791640 | c | g | 0.111 | 0.016 | 1.22E-12 | ++ | 0 | 0.456 | 1 | 0.500 | 1 | TFAP2B | 0 | | |
| rs62405422 | 6 | 50796905 | t | c | -0.111 | 0.016 | 1.22E-12 | -- | 0 | 0.456 | 1 | 0.500 | 1 | TFAP2B | 0 | | |
| rs2206277 | 6 | 50798526 | t | c | 0.110 | 0.016 | 1.54E-12 | ++ | 0 | 0.500 | 1 | 0.480 | 1 | TFAP2B | 0 | | |
| rs987237 | 6 | 50803050 | a | g | -0.110 | 0.017 | 1.92E-10 | -- | 0 | 0.120 | 1 | 0.729 | 1 | TFAP2B | 0 | | |
| rs4715207 | 6 | 50809278 | t | c | 0.110 | 0.018 | 5.86E-10 | ++ | 0 | 0.180 | 1 | 0.671 | 1 | TFAP2B | 0 | | |
| rs3798526 | 6 | 50809787 | a | g | -0.110 | 0.018 | 5.86E-10 | -- | 0 | 0.180 | 1 | 0.671 | 1 | TFAP2B | 0 | | |
| rs62405437 | 6 | 50813729 | t | c | -0.109 | 0.018 | 7.01E-10 | -- | 0 | 0.157 | 1 | 0.692 | 1 | TFAP2B | 0 | | |
| rs62405438 | 6 | 50815314 | a | t | 0.109 | 0.018 | 7.01E-10 | ++ | 0 | 0.157 | 1 | 0.692 | 1 | TFAP2B | 0 | | |
| rs72892910 | 6 | 50816887 | t | g | 0.109 | 0.018 | 7.01E-10 | ++ | 0 | 0.157 | 1 | 0.692 | 1 | TFAP2B | 1561 | | |
| rs6904450 | 6 | 50921602 | a | t | -0.092 | 0.017 | 2.99E-08 | -- | 0 | 0.566 | 1 | 0.452 | 1 | FTH1P5 | -40427 | | |
| rs933064 | 6 | 131183678 | t | c | -0.089 | 0.015 | 3.93E-09 | -- | 0 | 0.010 | 1 | 0.921 | 1 | EPB41L2 | 0 | | |
| rs17059760 | 6 | 131205074 | t | g | -0.096 | 0.017 | 1.83E-08 | -- | 0 | 0.070 | 1 | 0.791 | 1 | EPB41L2 | 0 | | |
| rs2275064 | 6 | 131220539 | a | g | 0.092 | 0.017 | 3.30E-08 | ++ | 0 | 0.130 | 1 | 0.718 | 1 | EPB41L2 | 0 | | |
| rs17059787 | 6 | 131222893 | a | g | 0.092 | 0.017 | 3.30E-08 | ++ | 0 | 0.130 | 1 | 0.718 | 1 | EPB41L2 | 0 | | |
| rs17059799 | 6 | 131223370 | c | g | -0.092 | 0.017 | 3.30E-08 | -- | 0 | 0.130 | 1 | 0.718 | 1 | EPB41L2 | 0 | | |
| rs17059801 | 6 | 131223481 | a | g | 0.092 | 0.017 | 3.30E-08 | ++ | 0 | 0.130 | 1 | 0.718 | 1 | EPB41L2 | 0 | | |
| rs79989737 | 6 | 131223539 | a | g | -0.092 | 0.017 | 3.30E-08 | -- | 0 | 0.130 | 1 | 0.718 | 1 | EPB41L2 | 0 | | |
| rs3777448 | 6 | 131224356 | t | c | 0.092 | 0.017 | 3.30E-08 | ++ | 0 | 0.130 | 1 | 0.718 | 1 | EPB41L2 | 0 | | |
| rs17059808 | 6 | 131226488 | t | c | -0.092 | 0.017 | 3.30E-08 | -- | 0 | 0.130 | 1 | 0.718 | 1 | EPB41L2 | 0 | | |
| rs79151412 | 6 | 131226663 | a | t | 0.092 | 0.017 | 3.30E-08 | ++ | 0 | 0.130 | 1 | 0.718 | 1 | EPB41L2 | 0 | | |
| rs59036192 | 6 | 131264216 | t | c | 0.094 | 0.017 | 3.60E-08 | ++ | 0 | 0.147 | 1 | 0.702 | 1 | EPB41L2 | 0 | | |
| rs17059851 | 6 | 131269681 | t | c | 0.093 | 0.017 | 4.25E-08 | ++ | 0 | 0.087 | 1 | 0.768 | 1 | EPB41L2 | 0 | | |
| rs17059853 | 6 | 131275525 | a | g | 0.091 | 0.017 | 4.57E-08 | ++ | 0 | 0.090 | 1 | 0.764 | 1 | EPB41L2 | 0 | | |
| rs3822865 | 6 | 131276061 | a | c | -0.091 | 0.017 | 4.57E-08 | -- | 0 | 0.090 | 1 | 0.764 | 1 | EPB41L2 | 0 | | |
| rs45577234 | 6 | 131276530 | t | c | 0.091 | 0.017 | 4.57E-08 | ++ | 0 | 0.090 | 1 | 0.764 | 1 | EPB41L2 | 0 | | |
| rs59271461 | 6 | 131294183 | t | c | -0.098 | 0.018 | 2.96E-08 | -- | 0 | 0.115 | 1 | 0.734 | 1 | EPB41L2 | 0 | | |
| rs12530288 | 6 | 131298399 | a | t | 0.081 | 0.015 | 4.90E-08 | ++ | 0 | 0.113 | 1 | 0.736 | 1 | EPB41L2 | 0 | | |
| rs9483201 | 6 | 131304860 | a | g | -0.082 | 0.015 | 4.05E-08 | -- | 0 | 0.092 | 1 | 0.762 | 1 | EPB41L2 | 0 | | |
| rs6928986 | 6 | 131323992 | t | c | 0.092 | 0.015 | 5.81E-10 | ++ | 0 | 0.163 | 1 | 0.686 | 1 | EPB41L2 | 0 | | |
| rs1888959 | 6 | 131335324 | t | c | 0.082 | 0.015 | 3.35E-08 | ++ | 0 | 0.163 | 1 | 0.686 | 1 | EPB41L2 | 0 | | |
| rs1855077 | 6 | 131342218 | a | g | -0.090 | 0.015 | 1.35E-09 | -- | 0 | 0.113 | 1 | 0.736 | 1 | EPB41L2 | 0 | | |
| rs7742376 | 6 | 131349836 | a | c | -0.087 | 0.015 | 5.70E-09 | -- | 0 | 0.010 | 1 | 0.920 | 1 | EPB41L2 | 0 | | |
| rs6909134 | 6 | 131354258 | t | c | -0.089 | 0.015 | 2.05E-09 | -- | 0 | 0.005 | 1 | 0.946 | 1 | EPB41L2 | 0 | | |
| rs6908917 | 6 | 131355864 | c | g | 0.087 | 0.015 | 5.70E-09 | ++ | 0 | 0.010 | 1 | 0.920 | 1 | EPB41L2 | 0 | | |
| rs1332490 | 6 | 131356873 | t | g | -0.090 | 0.015 | 1.67E-09 | -- | 0 | 0.137 | 1 | 0.711 | 1 | EPB41L2 | 0 | | |
| rs4897488 | 6 | 131362995 | a | g | 0.087 | 0.015 | 5.70E-09 | ++ | 0 | 0.010 | 1 | 0.920 | 1 | EPB41L2 | 0 | | |
| rs4895910 | 6 | 131371198 | a | g | -0.092 | 0.015 | 1.70E-09 | -- | 0 | 0.523 | 1 | 0.470 | 1 | EPB41L2 | 0 | | |
| rs6958406 | 7 | 30809318 | t | c | 0.099 | 0.017 | 6.47E-09 | ++ | 0 | 0.008 | 1 | 0.930 | 1 | INMT-FAM188B | 0 | | |
| rs12666880 | 7 | 30811746 | t | c | 0.095 | 0.017 | 9.21E-09 | ++ | 0 | 0.023 | 1 | 0.880 | 2 | INMT-FAM188B | 0 | FAM188B | 0 |
| rs73087704 | 7 | 30815535 | t | c | 0.098 | 0.017 | 7.71E-09 | ++ | 0 | 0.014 | 1 | 0.906 | 2 | INMT-FAM188B | 0 | FAM188B | 0 |
| rs12671080 | 7 | 30816908 | a | t | -0.099 | 0.017 | 6.47E-09 | -- | 0 | 0.008 | 1 | 0.930 | 2 | INMT-FAM188B | 0 | FAM188B | 0 |
| rs12667436 | 7 | 30818300 | t | c | 0.099 | 0.017 | 6.47E-09 | ++ | 0 | 0.008 | 1 | 0.930 | 2 | INMT-FAM188B | 0 | FAM188B | 0 |
| rs12672184 | 7 | 30825575 | a | c | 0.099 | 0.017 | 6.47E-09 | ++ | 0 | 0.043 | 1 | 0.837 | 2 | INMT-FAM188B | 0 | FAM188B | 0 |
| rs12665941 | 7 | 30828246 | a | g | -0.100 | 0.017 | 4.54E-09 | -- | 0 | 0.022 | 1 | 0.883 | 2 | INMT-FAM188B | 0 | FAM188B | 0 |
| rs12672119 | 7 | 30831181 | t | c | 0.101 | 0.017 | 3.18E-09 | ++ | 0 | 0.022 | 1 | 0.883 | 2 | INMT-FAM188B | 0 | FAM188B | 0 |
| rs79714917 | 7 | 30853003 | t | c | 0.099 | 0.016 | 1.39E-09 | ++ | 0 | 0.001 | 1 | 0.976 | 2 | INMT-FAM188B | 0 | FAM188B | 0 |
| rs76655111 | 7 | 30853248 | t | g | 0.099 | 0.016 | 1.39E-09 | ++ | 0 | 0.001 | 1 | 0.976 | 2 | INMT-FAM188B | 0 | FAM188B | 0 |
| rs76503667 | 7 | 30878389 | a | g | 0.108 | 0.018 | 1.19E-09 | ++ | 0 | 0.769 | 1 | 0.381 | 2 | INMT-FAM188B | 0 | FAM188B | 0 |
| rs12667955 | 7 | 30878708 | a | g | 0.108 | 0.018 | 1.19E-09 | ++ | 0 | 0.769 | 1 | 0.381 | 2 | INMT-FAM188B | 0 | FAM188B | 0 |
| rs59578124 | 7 | 30879148 | a | c | -0.106 | 0.018 | 2.02E-09 | -- | 0 | 0.925 | 1 | 0.336 | 2 | INMT-FAM188B | 0 | FAM188B | 0 |
| rs12672329 | 7 | 30879317 | a | g | -0.108 | 0.018 | 1.19E-09 | -- | 0 | 0.769 | 1 | 0.381 | 2 | INMT-FAM188B | 0 | FAM188B | 0 |
| rs78466775 | 7 | 30879980 | c | g | 0.108 | 0.018 | 1.19E-09 | ++ | 0 | 0.769 | 1 | 0.381 | 2 | INMT-FAM188B | 0 | FAM188B | 0 |
| rs76719927 | 7 | 30880075 | a | t | -0.108 | 0.018 | 1.19E-09 | -- | 0 | 0.769 | 1 | 0.381 | 2 | INMT-FAM188B | 0 | FAM188B | 0 |
| rs62449078 | 7 | 30880152 | a | c | -0.106 | 0.018 | 2.40E-09 | -- | 0 | 0.871 | 1 | 0.351 | 2 | INMT-FAM188B | 0 | FAM188B | 0 |
| rs59423432 | 7 | 30880628 | c | g | -0.107 | 0.018 | 1.42E-09 | -- | 0 | 0.720 | 1 | 0.396 | 2 | INMT-FAM188B | 0 | FAM188B | 0 |
| rs57742424 | 7 | 30880629 | a | g | -0.107 | 0.018 | 1.42E-09 | -- | 0 | 0.720 | 1 | 0.396 | 2 | INMT-FAM188B | 0 | FAM188B | 0 |
| rs75138450 | 7 | 30881147 | t | c | -0.107 | 0.018 | 1.42E-09 | -- | 0 | 0.720 | 1 | 0.396 | 2 | INMT-FAM188B | 0 | FAM188B | 0 |
| rs77395672 | 7 | 30881539 | a | t | 0.107 | 0.018 | 1.42E-09 | ++ | 0 | 0.720 | 1 | 0.396 | 2 | INMT-FAM188B | 0 | FAM188B | 0 |
| rs76504996 | 7 | 30882127 | t | c | 0.107 | 0.018 | 1.42E-09 | ++ | 0 | 0.720 | 1 | 0.396 | 2 | INMT-FAM188B | 0 | FAM188B | 0 |
| rs74902332 | 7 | 30882180 | t | c | 0.107 | 0.018 | 1.42E-09 | ++ | 0 | 0.720 | 1 | 0.396 | 2 | INMT-FAM188B | 0 | FAM188B | 0 |
| rs79035878 | 7 | 30884472 | t | g | 0.107 | 0.018 | 1.70E-09 | ++ | 0 | 0.673 | 1 | 0.412 | 2 | INMT-FAM188B | 0 | FAM188B | 0 |
| rs185913886 | 7 | 30885331 | a | c | -0.107 | 0.018 | 1.70E-09 | -- | 0 | 0.673 | 1 | 0.412 | 2 | INMT-FAM188B | 0 | FAM188B | 0 |
| rs189803492 | 7 | 30885332 | a | t | 0.107 | 0.018 | 1.70E-09 | ++ | 0 | 0.673 | 1 | 0.412 | 2 | INMT-FAM188B | 0 | FAM188B | 0 |
| rs74917173 | 7 | 30885519 | a | g | -0.107 | 0.018 | 1.70E-09 | -- | 0 | 0.673 | 1 | 0.412 | 2 | INMT-FAM188B | 0 | FAM188B | 0 |

| | | | | | | | | | | | | | | | | | |
|---|---|---|---|---|---|---|---|---|---|---|---|---|---|---|---|---|---|
| rs79925923 | 7 | 30885524 | t | g | -0.107 | 0.018 | 1.70E-09 | -- | 0 | 0.673 | 1 | 0.412 | 2 | INMT-FAM188B | 0 | FAM188B | 0 |
| rs80253417 | 7 | 30885652 | a | g | 0.106 | 0.018 | 2.02E-09 | ++ | 0 | 0.720 | 1 | 0.396 | 2 | INMT-FAM188B | 0 | FAM188B | 0 |
| rs77124469 | 7 | 30885781 | t | c | -0.106 | 0.018 | 2.02E-09 | -- | 0 | 0.720 | 1 | 0.396 | 2 | INMT-FAM188B | 0 | FAM188B | 0 |
| rs7778852 | 7 | 30886232 | c | g | 0.105 | 0.018 | 3.39E-09 | ++ | 0 | 0.769 | 1 | 0.381 | 2 | INMT-FAM188B | 0 | FAM188B | 0 |
| rs116973897 | 7 | 30886458 | a | g | -0.106 | 0.018 | 2.40E-09 | -- | 0 | 0.673 | 1 | 0.412 | 2 | INMT-FAM188B | 0 | FAM188B | 0 |
| rs7791980 | 7 | 30886500 | a | g | 0.106 | 0.018 | 2.40E-09 | ++ | 0 | 0.673 | 1 | 0.412 | 2 | INMT-FAM188B | 0 | FAM188B | 0 |
| rs7792019 | 7 | 30886563 | c | g | 0.106 | 0.018 | 2.40E-09 | ++ | 0 | 0.673 | 1 | 0.412 | 2 | INMT-FAM188B | 0 | FAM188B | 0 |
| rs6959949 | 7 | 30887230 | a | g | 0.105 | 0.018 | 3.39E-09 | ++ | 0 | 0.769 | 1 | 0.381 | 2 | INMT-FAM188B | 0 | FAM188B | 0 |
| rs1468312 | 7 | 30887781 | t | c | 0.106 | 0.018 | 2.40E-09 | ++ | 0 | 0.500 | 1 | 0.480 | 2 | INMT-FAM188B | 0 | FAM188B | 0 |
| rs1468313 | 7 | 30887782 | t | g | -0.106 | 0.018 | 2.40E-09 | -- | 0 | 0.500 | 1 | 0.480 | 2 | INMT-FAM188B | 0 | FAM188B | 0 |
| rs75796975 | 7 | 30887895 | c | g | -0.106 | 0.018 | 2.40E-09 | -- | 0 | 0.673 | 1 | 0.412 | 2 | INMT-FAM188B | 0 | FAM188B | 0 |
| rs34490481 | 7 | 30888042 | a | t | 0.106 | 0.018 | 2.40E-09 | ++ | 0 | 0.673 | 1 | 0.412 | 2 | INMT-FAM188B | 0 | FAM188B | 0 |
| rs12665880 | 7 | 30888114 | a | g | -0.105 | 0.018 | 2.86E-09 | -- | 0 | 0.720 | 1 | 0.396 | 2 | INMT-FAM188B | 0 | FAM188B | 0 |
| rs62449084 | 7 | 30888583 | a | g | 0.104 | 0.018 | 4.03E-09 | ++ | 0 | 0.819 | 1 | 0.365 | 2 | INMT-FAM188B | 0 | FAM188B | 0 |
| rs62449085 | 7 | 30888842 | c | g | -0.104 | 0.018 | 4.03E-09 | -- | 0 | 0.819 | 1 | 0.365 | 2 | INMT-FAM188B | 0 | FAM188B | 0 |
| rs62449086 | 7 | 30889051 | a | g | -0.103 | 0.018 | 5.66E-09 | -- | 0 | 0.819 | 1 | 0.365 | 2 | INMT-FAM188B | 0 | FAM188B | 0 |
| rs62449087 | 7 | 30889069 | t | c | 0.103 | 0.018 | 5.66E-09 | ++ | 0 | 0.819 | 1 | 0.365 | 2 | INMT-FAM188B | 0 | FAM188B | 0 |
| rs12671536 | 7 | 30890329 | t | c | 0.105 | 0.018 | 2.86E-09 | ++ | 0 | 0.627 | 1 | 0.428 | 2 | INMT-FAM188B | 0 | FAM188B | 0 |
| rs12672714 | 7 | 30892186 | a | g | 0.106 | 0.018 | 2.40E-09 | ++ | 0 | 0.583 | 1 | 0.445 | 2 | INMT-FAM188B | 0 | FAM188B | 0 |
| rs79987856 | 7 | 30893847 | a | t | 0.106 | 0.018 | 2.40E-09 | ++ | 0 | 0.583 | 1 | 0.445 | 2 | INMT-FAM188B | 0 | FAM188B | 0 |
| rs79483141 | 7 | 30893911 | t | c | 0.106 | 0.018 | 2.40E-09 | ++ | 0 | 0.583 | 1 | 0.445 | 2 | INMT-FAM188B | 0 | FAM188B | 0 |
| rs60168018 | 7 | 30894257 | a | g | 0.106 | 0.018 | 2.02E-09 | ++ | 0 | 0.541 | 1 | 0.462 | 2 | INMT-FAM188B | 0 | FAM188B | 0 |
| rs58640303 | 7 | 30894294 | t | g | 0.106 | 0.018 | 2.02E-09 | ++ | 0 | 0.541 | 1 | 0.462 | 2 | INMT-FAM188B | 0 | FAM188B | 0 |
| rs12670734 | 7 | 30895840 | a | g | -0.107 | 0.018 | 1.70E-09 | -- | 0 | 0.500 | 1 | 0.480 | 2 | INMT-FAM188B | 0 | FAM188B | 0 |
| rs12674155 | 7 | 30899543 | t | c | 0.107 | 0.018 | 1.42E-09 | ++ | 0 | 0.320 | 1 | 0.572 | 2 | INMT-FAM188B | 0 | FAM188B | 0 |
| rs111946193 | 7 | 30900307 | t | c | -0.108 | 0.018 | 1.19E-09 | -- | 0 | 0.289 | 1 | 0.591 | 2 | INMT-FAM188B | 0 | FAM188B | 0 |
| rs12669187 | 7 | 30915478 | a | g | 0.109 | 0.018 | 3.60E-09 | ++ | 0 | 0.060 | 1 | 0.807 | 2 | INMT-FAM188B | 0 | FAM188B | 0 |
| rs12669292 | 7 | 30916284 | a | c | 0.134 | 0.022 | 9.77E-10 | ++ | 0 | 0.033 | 1 | 0.855 | 2 | INMT-FAM188B | 0 | FAM188B | 0 |
| rs12666466 | 7 | 30916430 | c | g | -0.134 | 0.022 | 9.77E-10 | -- | 0 | 0.033 | 1 | 0.855 | 2 | INMT-FAM188B | 0 | FAM188B | 0 |
| rs73301967 | 7 | 30917117 | a | g | 0.134 | 0.022 | 9.77E-10 | ++ | 0 | 0.033 | 1 | 0.855 | 2 | INMT-FAM188B | 0 | FAM188B | 0 |
| rs73301968 | 7 | 30917675 | a | t | -0.134 | 0.022 | 9.77E-10 | -- | 0 | 0.033 | 1 | 0.855 | 2 | INMT-FAM188B | 0 | FAM188B | 0 |
| rs6975977 | 7 | 30917831 | a | g | 0.130 | 0.021 | 8.89E-10 | ++ | 0 | 0.002 | 1 | 0.962 | 2 | INMT-FAM188B | 0 | FAM188B | 0 |
| rs73303991 | 7 | 30918764 | t | g | -0.109 | 0.018 | 3.60E-09 | -- | 0 | 0.060 | 1 | 0.807 | 2 | INMT-FAM188B | 0 | FAM188B | 0 |
| rs34373236 | 7 | 30932837 | t | g | 0.102 | 0.018 | 9.37E-09 | ++ | 0 | 0.039 | 1 | 0.843 | 2 | INMT-FAM188B | 835 | FAM188B | 835 |
| rs80200031 | 7 | 30934436 | t | c | 0.118 | 0.020 | 2.95E-09 | ++ | 0 | 0.695 | 1 | 0.405 | 2 | INMT-FAM188B | 2434 | FAM188B | 2434 |
| rs4723021 | 7 | 30934659 | t | c | 0.101 | 0.018 | 1.31E-08 | ++ | 0 | 0.231 | 1 | 0.631 | 2 | INMT-FAM188B | 2657 | FAM188B | 2657 |
| rs12666237 | 7 | 30935739 | t | c | 0.102 | 0.018 | 7.93E-09 | ++ | 0 | 0.205 | 1 | 0.651 | 2 | INMT-FAM188B | 3737 | FAM188B | 3737 |
| rs1000597 | 7 | 30937178 | t | c | -0.098 | 0.018 | 2.96E-08 | -- | 0 | 0.157 | 1 | 0.692 | 2 | INMT-FAM188B | 5176 | FAM188B | 5176 |
| rs2324894 | 13 | 42642120 | a | t | 0.120 | 0.016 | 1.22E-14 | ++ | 58 | 2.380 | 1 | 0.123 | 2 | DGKH | 0 | MAPK6PS3 | 0 |
| rs9566906 | 13 | 42642546 | a | g | -0.115 | 0.016 | 4.40E-13 | -- | 72.8 | 3.673 | 1 | 0.055 | 2 | DGKH | 0 | MAPK6PS3 | 0 |
| rs9315885 | 13 | 42642810 | t | c | -0.118 | 0.015 | 2.52E-15 | -- | 70.8 | 3.430 | 1 | 0.064 | 2 | DGKH | 0 | MAPK6PS3 | 0 |
| rs9525566 | 13 | 42643194 | a | g | 0.113 | 0.015 | 8.29E-14 | ++ | 58.1 | 2.388 | 1 | 0.122 | 2 | DGKH | 0 | MAPK6PS3 | 0 |
| rs7993632 | 13 | 42644176 | a | t | -0.118 | 0.015 | 2.52E-15 | -- | 70.8 | 3.430 | 1 | 0.064 | 2 | DGKH | 0 | MAPK6PS3 | 0 |
| rs9532985 | 13 | 42644377 | a | c | 0.114 | 0.015 | 2.11E-14 | ++ | 52.3 | 2.096 | 1 | 0.148 | 2 | DGKH | 0 | MAPK6PS3 | 0 |
| rs12427854 | 13 | 42645358 | a | g | 0.116 | 0.015 | 5.64E-15 | ++ | 71.9 | 3.556 | 1 | 0.059 | 2 | DGKH | 0 | MAPK6PS3 | 0 |
| rs9566908 | 13 | 42645981 | t | c | -0.111 | 0.016 | 3.38E-12 | -- | 71.9 | 3.554 | 1 | 0.059 | 2 | DGKH | 0 | MAPK6PS3 | 0 |
| rs7999315 | 13 | 42646593 | t | g | 0.113 | 0.015 | 3.56E-14 | ++ | 63.3 | 2.722 | 1 | 0.099 | 1 | DGKH | 0 | | |
| rs73187215 | 13 | 42646769 | a | g | -0.111 | 0.015 | 7.71E-14 | -- | 64.7 | 2.834 | 1 | 0.092 | 1 | DGKH | 0 | | |
| rs7990452 | 13 | 42649220 | a | t | 0.116 | 0.015 | 7.36E-15 | ++ | 70.8 | 3.430 | 1 | 0.064 | 1 | DGKH | 0 | | |
| rs9532987 | 13 | 42649234 | a | g | 0.116 | 0.015 | 7.36E-15 | ++ | 70.8 | 3.430 | 1 | 0.064 | 1 | DGKH | 0 | | |
| rs9562372 | 13 | 42649246 | t | c | 0.114 | 0.016 | 2.33E-13 | ++ | 74.8 | 3.971 | 1 | 0.046 | 1 | DGKH | 0 | | |
| rs9532989 | 13 | 42649810 | t | g | -0.116 | 0.015 | 7.36E-15 | -- | 70.8 | 3.430 | 1 | 0.064 | 1 | DGKH | 0 | | |
| rs4142112 | 13 | 42650094 | c | g | -0.116 | 0.015 | 7.36E-15 | -- | 70.8 | 3.430 | 1 | 0.064 | 1 | DGKH | 0 | | |
| rs1170183 | 13 | 42650733 | a | c | -0.112 | 0.015 | 4.61E-14 | -- | 61.7 | 2.612 | 1 | 0.106 | 1 | DGKH | 0 | | |
| rs9566910 | 13 | 42651002 | a | g | -0.114 | 0.016 | 2.33E-13 | -- | 74.8 | 3.971 | 1 | 0.046 | 1 | DGKH | 0 | | |
| rs942896 | 13 | 42651460 | a | t | 0.116 | 0.015 | 7.36E-15 | ++ | 70.8 | 3.430 | 1 | 0.064 | 1 | DGKH | 0 | | |
| rs2021802 | 13 | 42652165 | t | c | -0.110 | 0.016 | 4.26E-12 | -- | 72.8 | 3.673 | 1 | 0.055 | 1 | DGKH | 0 | | |
| rs1012053 | 13 | 42653437 | a | c | -0.112 | 0.015 | 5.97E-14 | -- | 56.4 | 2.296 | 1 | 0.130 | 1 | DGKH | 0 | | |
| rs9532990 | 13 | 42653898 | t | c | -0.111 | 0.016 | 9.65E-13 | -- | 69.1 | 3.240 | 1 | 0.072 | 1 | DGKH | 0 | | |
| rs9532991 | 13 | 42654189 | a | g | -0.111 | 0.016 | 9.65E-13 | -- | 69.1 | 3.240 | 1 | 0.072 | 1 | DGKH | 0 | | |
| rs2253650 | 13 | 42656841 | t | c | -0.112 | 0.015 | 4.61E-14 | -- | 54.4 | 2.195 | 1 | 0.139 | 1 | DGKH | 0 | | |
| rs35208748 | 13 | 42658751 | a | t | 0.112 | 0.015 | 4.61E-14 | ++ | 54.4 | 2.195 | 1 | 0.139 | 1 | DGKH | 0 | | |
| rs669642 | 13 | 42660845 | c | g | -0.114 | 0.015 | 2.11E-14 | -- | 52.3 | 2.096 | 1 | 0.148 | 1 | DGKH | 0 | | |
| rs1170203 | 13 | 42661379 | c | g | -0.114 | 0.015 | 2.11E-14 | -- | 52.3 | 2.096 | 1 | 0.148 | 1 | DGKH | 0 | | |
| rs7993986 | 13 | 42661742 | a | g | -0.112 | 0.016 | 1.60E-12 | -- | 63.9 | 2.773 | 1 | 0.096 | 1 | DGKH | 0 | | |
| rs681657 | 13 | 42662600 | a | g | -0.114 | 0.015 | 1.63E-14 | -- | 50 | 2.000 | 1 | 0.157 | 1 | DGKH | 0 | | |
| rs9525568 | 13 | 42663251 | a | g | 0.112 | 0.016 | 1.60E-12 | ++ | 63.9 | 2.773 | 1 | 0.096 | 1 | DGKH | 0 | | |
| rs1182959 | 13 | 42674601 | a | g | 0.127 | 0.015 | 1.61E-17 | ++ | 0 | 0.600 | 1 | 0.439 | 1 | DGKH | 0 | | |
| rs1170191 | 13 | 42675493 | a | g | 0.127 | 0.015 | 1.20E-17 | ++ | 0 | 0.653 | 1 | 0.419 | 1 | DGKH | 0 | | |
| rs1170190 | 13 | 42675667 | a | g | 0.127 | 0.015 | 1.20E-17 | ++ | 0 | 0.653 | 1 | 0.419 | 1 | DGKH | 0 | | |
| rs672898 | 13 | 42676537 | t | c | 0.127 | 0.015 | 1.20E-17 | ++ | 0 | 0.653 | 1 | 0.419 | 1 | DGKH | 0 | | |
| rs912875 | 13 | 42677238 | a | c | 0.126 | 0.016 | 7.18E-16 | ++ | 29.3 | 1.414 | 1 | 0.234 | 1 | DGKH | 0 | | |
| rs7984523 | 13 | 42678357 | t | c | -0.132 | 0.015 | 8.32E-19 | -- | 19 | 1.235 | 1 | 0.267 | 1 | DGKH | 0 | | |
| rs1170188 | 13 | 42678971 | a | g | 0.126 | 0.015 | 2.15E-17 | ++ | 0 | 0.653 | 1 | 0.419 | 1 | DGKH | 0 | | |
| rs611760 | 13 | 42679595 | a | c | -0.127 | 0.015 | 1.20E-17 | -- | 0 | 0.653 | 1 | 0.419 | 1 | DGKH | 0 | | |
| rs1170187 | 13 | 42679835 | a | g | -0.127 | 0.015 | 1.20E-17 | -- | 0 | 0.653 | 1 | 0.419 | 1 | DGKH | 0 | | |
| rs7333528 | 13 | 42680253 | t | c | -0.126 | 0.016 | 7.18E-16 | -- | 29.3 | 1.414 | 1 | 0.234 | 1 | DGKH | 0 | | |
| rs2767389 | 13 | 42681678 | c | g | -0.115 | 0.015 | 3.87E-14 | -- | 0 | 0.676 | 1 | 0.411 | 1 | DGKH | 0 | | |
| rs6561034 | 13 | 42685221 | t | c | -0.126 | 0.016 | 7.18E-16 | -- | 29.3 | 1.414 | 1 | 0.234 | 1 | DGKH | 0 | | |
| rs9566919 | 13 | 42685577 | a | g | -0.111 | 0.020 | 2.39E-08 | -- | 0 | 0.536 | 1 | 0.464 | 1 | DGKH | 0 | | |
| rs9525576 | 13 | 42685989 | t | c | 0.111 | 0.020 | 2.39E-08 | ++ | 0 | 0.337 | 1 | 0.561 | 1 | DGKH | 0 | | |
| rs686810 | 13 | 42686172 | t | c | 0.115 | 0.015 | 3.87E-14 | ++ | 0 | 0.676 | 1 | 0.411 | 1 | DGKH | 0 | | |
| rs657351 | 13 | 42686263 | t | c | -0.115 | 0.015 | 3.87E-14 | -- | 0 | 0.676 | 1 | 0.411 | 1 | DGKH | 0 | | |
| rs685951 | 13 | 42686359 | a | g | 0.128 | 0.015 | 8.98E-18 | ++ | 0 | 0.709 | 1 | 0.400 | 1 | DGKH | 0 | | |
| rs1170174 | 13 | 42688211 | a | g | 0.127 | 0.015 | 1.20E-17 | ++ | 0 | 0.766 | 1 | 0.381 | 1 | DGKH | 0 | | |
| rs1170173 | 13 | 42688615 | a | g | 0.128 | 0.015 | 8.98E-18 | ++ | 0 | 0.709 | 1 | 0.400 | 1 | DGKH | 0 | | |
| rs1170172 | 13 | 42688645 | a | g | -0.128 | 0.015 | 8.98E-18 | -- | 0 | 0.600 | 1 | 0.439 | 1 | DGKH | 0 | | |
| rs9533002 | 13 | 42689009 | t | c | -0.110 | 0.020 | 2.76E-08 | -- | 0 | 0.367 | 1 | 0.545 | 1 | DGKH | 0 | | |
| rs1170171 | 13 | 42689717 | c | g | -0.115 | 0.015 | 2.97E-14 | -- | 0 | 0.623 | 1 | 0.430 | 1 | DGKH | 0 | | |
| rs585206 | 13 | 42689853 | a | g | 0.115 | 0.015 | 2.97E-14 | ++ | 0 | 0.623 | 1 | 0.430 | 1 | DGKH | 0 | | |
| rs7981733 | 13 | 42690060 | t | c | -0.126 | 0.016 | 7.18E-16 | -- | 29.3 | 1.414 | 1 | 0.234 | 1 | DGKH | 0 | | |
| rs1177637 | 13 | 42692823 | a | g | -0.130 | 0.015 | 2.76E-18 | -- | 0 | 0.709 | 1 | 0.400 | 1 | DGKH | 0 | | |
| rs9566923 | 13 | 42694928 | a | g | -0.114 | 0.019 | 4.65E-09 | -- | 0 | 0.291 | 1 | 0.589 | 1 | DGKH | 0 | | |
| rs1170169 | 13 | 42697807 | c | g | -0.118 | 0.016 | 3.32E-14 | -- | 0 | 0.595 | 1 | 0.441 | 1 | DGKH | 0 | | |
| rs2148004 | 13 | 42697839 | a | g | 0.114 | 0.019 | 2.36E-09 | ++ | 0 | 0.274 | 1 | 0.600 | 1 | DGKH | 0 | | |
| rs635852 | 13 | 42698310 | a | c | 0.118 | 0.016 | 3.32E-14 | ++ | 0 | 0.595 | 1 | 0.441 | 1 | DGKH | 0 | | |
| rs636264 | 13 | 42698380 | a | g | -0.130 | 0.015 | 2.76E-18 | -- | 0 | 0.709 | 1 | 0.400 | 1 | DGKH | 0 | | |
| rs9315891 | 13 | 42699079 | t | c | -0.132 | 0.016 | 2.84E-17 | -- | 42.4 | 1.737 | 1 | 0.188 | 1 | DGKH | 0 | | |
| rs9533003 | 13 | 42699604 | c | g | 0.125 | 0.016 | 9.34E-16 | ++ | 25.3 | 1.339 | 1 | 0.247 | 1 | DGKH | 0 | | |
| rs670676 | 13 | 42701739 | a | t | 0.134 | 0.015 | 1.81E-19 | ++ | 13.9 | 1.161 | 1 | 0.281 | 1 | DGKH | 0 | | |
| rs1170158 | 13 | 42701941 | t | g | -0.134 | 0.015 | 1.81E-19 | -- | 13.9 | 1.161 | 1 | 0.281 | 1 | DGKH | 0 | | |
| rs1170155 | 13 | 42702711 | t | c | -0.125 | 0.016 | 9.34E-16 | -- | 0 | 0.500 | 1 | 0.480 | 1 | DGKH | 0 | | |
| rs1184924 | 13 | 42704400 | a | g | 0.131 | 0.015 | 1.52E-18 | ++ | 60.1 | 2.505 | 1 | 0.114 | 1 | DGKH | 0 | | |
| rs1750017 | 13 | 42704451 | a | g | -0.131 | 0.015 | 1.52E-18 | -- | 60.1 | 2.505 | 1 | 0.114 | 1 | DGKH | 0 | | |
| rs9562380 | 13 | 42704847 | a | g | -0.114 | 0.019 | 2.77E-09 | -- | 0 | 0.198 | 1 | 0.656 | 1 | DGKH | 0 | | |
| rs1170178 | 13 | 42705808 | t | c | 0.130 | 0.015 | 2.05E-18 | ++ | 61.7 | 2.612 | 1 | 0.106 | 1 | DGKH | 0 | | |
| rs619659 | 13 | 42706673 | a | g | -0.125 | 0.016 | 9.34E-16 | -- | 0 | 0.595 | 1 | 0.441 | 1 | DGKH | 0 | | |
| rs9533006 | 13 | 42707301 | t | c | 0.115 | 0.020 | 7.33E-09 | ++ | 0 | 0.143 | 1 | 0.705 | 1 | DGKH | 0 | | |
| rs7983995 | 13 | 42708231 | a | c | 0.114 | 0.019 | 2.77E-09 | ++ | 0 | 0.198 | 1 | 0.656 | 1 | DGKH | 0 | | |
| rs682573 | 13 | 42708655 | t | g | -0.129 | 0.015 | 3.71E-18 | -- | 77.1 | 4.358 | 1 | 0.037 | 1 | DGKH | 0 | | |
| rs7327259 | 13 | 42709345 | a | g | -0.114 | 0.019 | 2.77E-09 | -- | 0 | 0.198 | 1 | 0.656 | 1 | DGKH | 0 | | |

| | | | | | | | | | | | | | | | |
|---|---|---|---|---|---|---|---|---|---|---|---|---|---|---|---|
| rs9533007 | 13 | 42710235 | a | g | -0.125 | 0.016 | 9.34E-16 | -- | 0 | 0.595 | 1 | 0.441 | 1 | DGKH | 0 |
| rs12585075 | 13 | 42711224 | t | c | 0.130 | 0.015 | 2.05E-18 | ++ | 61.7 | 2.612 | 1 | 0.106 | 1 | DGKH | 0 |
| rs12876965 | 13 | 42712741 | a | t | -0.130 | 0.015 | 2.05E-18 | -- | 61.7 | 2.612 | 1 | 0.106 | 1 | DGKH | 0 |
| rs17696391 | 13 | 42713049 | a | g | -0.132 | 0.015 | 8.32E-19 | -- | 60.1 | 2.505 | 1 | 0.114 | 1 | DGKH | 0 |
| rs57995007 | 13 | 42715118 | t | c | 0.131 | 0.015 | 1.52E-18 | ++ | 60.1 | 2.505 | 1 | 0.114 | 1 | DGKH | 0 |
| rs35792791 | 13 | 42715617 | t | c | 0.131 | 0.015 | 1.52E-18 | ++ | 60.1 | 2.505 | 1 | 0.114 | 1 | DGKH | 0 |
| rs80067912 | 13 | 42717067 | a | g | 0.131 | 0.016 | 4.91E-17 | ++ | 47.6 | 1.910 | 1 | 0.167 | 1 | DGKH | 0 |
| rs9533010 | 13 | 42717279 | c | g | 0.114 | 0.019 | 2.77E-09 | ++ | 0 | 0.198 | 1 | 0.656 | 1 | DGKH | 0 |
| rs9525579 | 13 | 42717556 | a | t | 0.114 | 0.019 | 2.77E-09 | ++ | 0 | 0.198 | 1 | 0.656 | 1 | DGKH | 0 |
| rs12867964 | 13 | 42717898 | t | g | -0.133 | 0.015 | 3.35E-19 | -- | 44.9 | 1.814 | 1 | 0.178 | 1 | DGKH | 0 |
| rs9533011 | 13 | 42720204 | a | g | 0.114 | 0.019 | 2.77E-09 | ++ | 0 | 0.198 | 1 | 0.656 | 1 | DGKH | 0 |
| rs34695805 | 13 | 42720666 | t | c | 0.131 | 0.015 | 1.12E-18 | ++ | 61.7 | 2.612 | 1 | 0.106 | 1 | DGKH | 0 |
| rs7329683 | 13 | 42721707 | c | g | 0.114 | 0.019 | 2.77E-09 | ++ | 0 | 0.198 | 1 | 0.656 | 1 | DGKH | 0 |
| rs9533012 | 13 | 42722882 | t | c | -0.114 | 0.019 | 2.77E-09 | -- | 0 | 0.198 | 1 | 0.656 | 1 | DGKH | 0 |
| rs9566931 | 13 | 42724498 | a | g | 0.114 | 0.019 | 2.77E-09 | ++ | 0 | 0.198 | 1 | 0.656 | 1 | DGKH | 0 |
| rs71429406 | 13 | 42725370 | a | g | 0.131 | 0.015 | 1.12E-18 | ++ | 61.7 | 2.612 | 1 | 0.106 | 1 | DGKH | 0 |
| rs9525583 | 13 | 42725929 | t | g | 0.113 | 0.019 | 3.24E-09 | ++ | 0 | 0.222 | 1 | 0.637 | 1 | DGKH | 0 |
| rs9533015 | 13 | 42726118 | a | g | -0.113 | 0.019 | 3.24E-09 | -- | 0 | 0.222 | 1 | 0.637 | 1 | DGKH | 0 |
| rs12872074 | 13 | 42728934 | a | t | 0.131 | 0.015 | 1.52E-18 | ++ | 47.5 | 1.906 | 1 | 0.167 | 1 | DGKH | 0 |
| rs56134371 | 13 | 42728992 | t | c | 0.113 | 0.019 | 3.80E-09 | ++ | 0 | 0.198 | 1 | 0.656 | 1 | DGKH | 0 |
| rs9533019 | 13 | 42730949 | a | g | 0.113 | 0.019 | 3.80E-09 | ++ | 0 | 0.198 | 1 | 0.656 | 1 | DGKH | 0 |
| rs10459372 | 13 | 42732565 | c | g | -0.113 | 0.019 | 3.80E-09 | -- | 0 | 0.198 | 1 | 0.656 | 1 | DGKH | 0 |
| rs34000170 | 13 | 42732805 | t | c | -0.132 | 0.015 | 6.15E-19 | -- | 61.7 | 2.612 | 1 | 0.106 | 1 | DGKH | 0 |
| rs9533020 | 13 | 42733029 | a | g | -0.113 | 0.019 | 3.80E-09 | -- | 0 | 0.198 | 1 | 0.656 | 1 | DGKH | 0 |
| rs2274287 | 13 | 42733328 | a | g | -0.132 | 0.015 | 6.15E-19 | -- | 61.7 | 2.612 | 1 | 0.106 | 1 | DGKH | 0 |
| rs9315894 | 13 | 42733977 | t | g | 0.113 | 0.019 | 3.80E-09 | ++ | 0 | 0.198 | 1 | 0.656 | 1 | DGKH | 0 |
| rs17645393 | 13 | 42734840 | a | g | 0.133 | 0.015 | 4.54E-19 | ++ | 63.3 | 2.722 | 1 | 0.099 | 1 | DGKH | 0 |
| rs9525584 | 13 | 42734894 | t | c | 0.110 | 0.019 | 9.73E-09 | ++ | 0 | 0.363 | 1 | 0.547 | 1 | DGKH | 0 |
| rs12865430 | 13 | 42736013 | t | g | 0.133 | 0.015 | 4.54E-19 | ++ | 63.3 | 2.722 | 1 | 0.099 | 1 | DGKH | 0 |
| rs9525585 | 13 | 42736710 | t | c | -0.113 | 0.019 | 3.24E-09 | -- | 0 | 0.222 | 1 | 0.637 | 1 | DGKH | 0 |
| rs12428755 | 13 | 42736747 | t | c | 0.135 | 0.015 | 9.78E-20 | ++ | 58.3 | 2.399 | 1 | 0.121 | 1 | DGKH | 0 |
| rs12430054 | 13 | 42737012 | t | c | -0.135 | 0.015 | 9.78E-20 | -- | 58.3 | 2.399 | 1 | 0.121 | 1 | DGKH | 0 |
| rs12430974 | 13 | 42737100 | a | g | -0.135 | 0.015 | 9.78E-20 | -- | 58.3 | 2.399 | 1 | 0.121 | 1 | DGKH | 0 |
| rs9315896 | 13 | 42738420 | a | g | 0.106 | 0.016 | 8.76E-11 | ++ | 0 | 0.114 | 1 | 0.735 | 1 | DGKH | 0 |
| rs12429318 | 13 | 42738831 | c | g | -0.133 | 0.015 | 3.35E-19 | -- | 64.7 | 2.834 | 1 | 0.092 | 1 | DGKH | 0 |
| rs9525588 | 13 | 42739233 | a | g | 0.106 | 0.016 | 8.76E-11 | ++ | 0 | 0.114 | 1 | 0.735 | 1 | DGKH | 0 |
| rs17598311 | 13 | 42739826 | a | g | 0.114 | 0.019 | 2.36E-09 | ++ | 0 | 0.222 | 1 | 0.637 | 1 | DGKH | 0 |
| rs7329496 | 13 | 42740908 | a | t | -0.133 | 0.015 | 3.35E-19 | -- | 64.7 | 2.834 | 1 | 0.092 | 1 | DGKH | 0 |
| rs1555819 | 13 | 42741873 | t | c | -0.114 | 0.019 | 2.36E-09 | -- | 0 | 0.222 | 1 | 0.637 | 1 | DGKH | 0 |
| rs12429782 | 13 | 42745294 | t | c | 0.134 | 0.015 | 2.46E-19 | ++ | 66.1 | 2.949 | 1 | 0.086 | 1 | DGKH | 0 |
| rs7328064 | 13 | 42746118 | a | c | -0.139 | 0.015 | 5.87E-20 | -- | 79.4 | 4.853 | 1 | 0.028 | 1 | DGKH | 0 |
| rs73187293 | 13 | 42746500 | t | c | 0.133 | 0.015 | 3.35E-19 | ++ | 67.4 | 3.066 | 1 | 0.080 | 1 | DGKH | 0 |
| rs9590675 | 13 | 42749192 | t | g | 0.131 | 0.015 | 1.52E-18 | ++ | 42 | 1.724 | 1 | 0.189 | 1 | DGKH | 0 |
| rs9533022 | 13 | 42749606 | t | c | -0.102 | 0.016 | 5.50E-11 | -- | 0 | 0.017 | 1 | 0.898 | 1 | DGKH | 0 |
| rs67332916 | 13 | 42749711 | t | c | 0.136 | 0.016 | 2.28E-18 | ++ | 66.8 | 3.012 | 1 | 0.083 | 1 | DGKH | 0 |
| rs12585865 | 13 | 42751707 | t | c | -0.130 | 0.015 | 2.76E-18 | -- | 35.6 | 1.552 | 1 | 0.213 | 1 | DGKH | 0 |
| rs1925762 | 13 | 42753787 | t | g | 0.130 | 0.018 | 2.38E-13 | ++ | 25.6 | 1.345 | 1 | 0.246 | 1 | DGKH | 0 |
| rs4142110 | 13 | 42754522 | t | c | -0.130 | 0.015 | 2.76E-18 | -- | 0 | 0.827 | 1 | 0.363 | 1 | DGKH | 0 |
| rs9525589 | 13 | 42755838 | t | c | 0.102 | 0.016 | 3.57E-10 | ++ | 0 | 0.095 | 1 | 0.759 | 1 | DGKH | 0 |
| rs34445998 | 13 | 42757213 | t | c | 0.130 | 0.015 | 2.05E-18 | ++ | 71.9 | 3.556 | 1 | 0.059 | 1 | DGKH | 0 |
| rs34028145 | 13 | 42757247 | c | g | 0.131 | 0.015 | 1.52E-18 | ++ | 70.8 | 3.430 | 1 | 0.064 | 1 | DGKH | 0 |
| rs35859771 | 13 | 42758189 | a | g | -0.131 | 0.015 | 1.52E-18 | -- | 70.8 | 3.430 | 1 | 0.064 | 1 | DGKH | 0 |
| rs36122479 | 13 | 42758435 | a | t | 0.131 | 0.015 | 1.52E-18 | ++ | 70.8 | 3.430 | 1 | 0.064 | 1 | DGKH | 0 |
| rs9590676 | 13 | 42758805 | t | c | -0.125 | 0.015 | 3.83E-17 | -- | 2 | 1.020 | 1 | 0.312 | 1 | DGKH | 0 |
| rs12585267 | 13 | 42759971 | a | c | -0.127 | 0.015 | 1.20E-17 | -- | 73.8 | 3.814 | 1 | 0.051 | 1 | DGKH | 0 |
| rs12876247 | 13 | 42761457 | t | c | 0.126 | 0.015 | 2.15E-17 | ++ | 73.8 | 3.814 | 1 | 0.051 | 1 | DGKH | 0 |
| rs12877348 | 13 | 42761522 | t | c | -0.126 | 0.015 | 2.15E-17 | -- | 73.8 | 3.814 | 1 | 0.051 | 1 | DGKH | 0 |
| rs12877052 | 13 | 42761891 | a | g | 0.126 | 0.015 | 2.15E-17 | ++ | 73.8 | 3.814 | 1 | 0.051 | 1 | DGKH | 0 |
| rs10492441 | 13 | 42762085 | c | g | -0.126 | 0.015 | 2.15E-17 | -- | 73.8 | 3.814 | 1 | 0.051 | 1 | DGKH | 0 |
| rs10492439 | 13 | 42762316 | c | g | 0.126 | 0.015 | 2.15E-17 | ++ | 73.8 | 3.814 | 1 | 0.051 | 1 | DGKH | 0 |
| rs17645560 | 13 | 42762421 | t | g | -0.126 | 0.015 | 2.15E-17 | -- | 73.8 | 3.814 | 1 | 0.051 | 1 | DGKH | 0 |
| rs34556067 | 13 | 42762524 | t | c | -0.126 | 0.015 | 2.15E-17 | -- | 73.8 | 3.814 | 1 | 0.051 | 1 | DGKH | 0 |
| rs4640076 | 13 | 42762620 | t | c | -0.127 | 0.015 | 1.61E-17 | -- | 74.7 | 3.947 | 1 | 0.047 | 1 | DGKH | 0 |
| rs4272927 | 13 | 42762829 | t | g | 0.127 | 0.015 | 1.61E-17 | ++ | 74.7 | 3.947 | 1 | 0.047 | 1 | DGKH | 0 |
| rs4598803 | 13 | 42762871 | a | g | -0.127 | 0.015 | 1.61E-17 | -- | 74.7 | 3.947 | 1 | 0.047 | 1 | DGKH | 0 |
| rs2167059 | 13 | 42762971 | a | g | 0.126 | 0.015 | 2.87E-17 | ++ | 0 | 0.954 | 1 | 0.329 | 1 | DGKH | 0 |
| rs2122246 | 13 | 42763054 | a | g | 0.127 | 0.015 | 1.61E-17 | ++ | 74.7 | 3.947 | 1 | 0.047 | 1 | DGKH | 0 |
| rs2122247 | 13 | 42763140 | a | g | 0.127 | 0.015 | 1.61E-17 | ++ | 74.7 | 3.947 | 1 | 0.047 | 1 | DGKH | 0 |
| rs9525594 | 13 | 42763154 | a | g | -0.120 | 0.022 | 2.43E-08 | -- | 0 | 0.138 | 1 | 0.711 | 1 | DGKH | 0 |
| rs9566933 | 13 | 42763505 | t | c | -0.120 | 0.022 | 2.43E-08 | -- | 0 | 0.138 | 1 | 0.711 | 1 | DGKH | 0 |
| rs12868961 | 13 | 42763813 | a | t | -0.127 | 0.016 | 1.19E-15 | -- | 66.5 | 2.986 | 1 | 0.084 | 1 | DGKH | 0 |
| rs75066565 | 13 | 42763814 | a | t | -0.117 | 0.016 | 5.43E-14 | -- | 61.3 | 2.583 | 1 | 0.108 | 1 | DGKH | 0 |
| rs67462156 | 13 | 42763871 | t | c | 0.131 | 0.015 | 1.12E-18 | ++ | 75.5 | 4.082 | 1 | 0.043 | 1 | DGKH | 0 |
| rs35535159 | 13 | 42763934 | t | c | -0.128 | 0.015 | 8.98E-18 | -- | 74.7 | 3.947 | 1 | 0.047 | 1 | DGKH | 0 |
| rs35993291 | 13 | 42764048 | t | c | -0.130 | 0.015 | 2.05E-18 | -- | 77.1 | 4.358 | 1 | 0.037 | 1 | DGKH | 0 |
| rs67192248 | 13 | 42764314 | t | g | -0.126 | 0.015 | 2.15E-17 | -- | 73.8 | 3.814 | 1 | 0.051 | 1 | DGKH | 0 |
| rs36095934 | 13 | 42764440 | t | g | 0.124 | 0.016 | 1.57E-15 | ++ | 64.2 | 2.793 | 1 | 0.095 | 1 | DGKH | 0 |
| rs17645590 | 13 | 42765049 | t | c | -0.127 | 0.015 | 1.20E-17 | -- | 73.8 | 3.814 | 1 | 0.051 | 1 | DGKH | 0 |
| rs17645596 | 13 | 42765098 | a | g | 0.127 | 0.015 | 1.20E-17 | ++ | 73.8 | 3.814 | 1 | 0.051 | 1 | DGKH | 0 |
| rs12854774 | 13 | 42765334 | t | g | 0.128 | 0.015 | 8.98E-18 | ++ | 74.7 | 3.947 | 1 | 0.047 | 1 | DGKH | 0 |
| rs12854131 | 13 | 42765400 | a | c | -0.128 | 0.015 | 8.98E-18 | -- | 74.7 | 3.947 | 1 | 0.047 | 1 | DGKH | 0 |
| rs61959254 | 13 | 42765404 | t | c | -0.120 | 0.022 | 2.43E-08 | -- | 0 | 0.138 | 1 | 0.711 | 1 | DGKH | 0 |
| rs12854149 | 13 | 42765419 | a | g | -0.128 | 0.015 | 8.98E-18 | -- | 74.7 | 3.947 | 1 | 0.047 | 1 | DGKH | 0 |
| rs12857339 | 13 | 42765450 | t | c | -0.124 | 0.015 | 9.03E-17 | -- | 60.1 | 2.505 | 1 | 0.114 | 1 | DGKH | 0 |
| rs34059466 | 13 | 42765546 | t | g | -0.127 | 0.015 | 1.61E-17 | -- | 0 | 0.954 | 1 | 0.329 | 1 | DGKH | 0 |
| rs34749620 | 13 | 42765608 | c | g | -0.128 | 0.015 | 8.98E-18 | -- | 74.7 | 3.947 | 1 | 0.047 | 1 | DGKH | 0 |
| rs17645608 | 13 | 42765819 | a | g | -0.128 | 0.015 | 6.70E-18 | -- | 73.8 | 3.814 | 1 | 0.051 | 1 | DGKH | 0 |
| rs4941418 | 13 | 42766652 | a | c | -0.126 | 0.015 | 2.87E-17 | -- | 0 | 0.954 | 1 | 0.329 | 1 | DGKH | 0 |
| rs12867273 | 13 | 42766748 | t | c | -0.145 | 0.017 | 2.77E-18 | -- | 64.8 | 2.838 | 1 | 0.092 | 1 | DGKH | 0 |
| rs347389 | 13 | 42766823 | a | t | 0.127 | 0.015 | 1.61E-17 | ++ | 19 | 1.235 | 1 | 0.267 | 1 | DGKH | 0 |
| rs12870816 | 13 | 42767252 | a | g | 0.127 | 0.015 | 1.20E-17 | ++ | 73.8 | 3.814 | 1 | 0.051 | 1 | DGKH | 0 |
| rs12870823 | 13 | 42767261 | a | g | 0.127 | 0.015 | 1.20E-17 | ++ | 73.8 | 3.814 | 1 | 0.051 | 1 | DGKH | 0 |
| rs12870877 | 13 | 42767479 | a | c | 0.127 | 0.015 | 1.20E-17 | ++ | 73.8 | 3.814 | 1 | 0.051 | 1 | DGKH | 0 |
| rs10492438 | 13 | 42767706 | a | c | 0.127 | 0.015 | 1.20E-17 | ++ | 73.8 | 3.814 | 1 | 0.051 | 1 | DGKH | 0 |
| rs10492437 | 13 | 42767900 | a | t | 0.129 | 0.015 | 4.99E-18 | ++ | 76.3 | 4.219 | 1 | 0.040 | 1 | DGKH | 0 |
| rs12875392 | 13 | 42768108 | c | g | -0.128 | 0.015 | 6.70E-18 | -- | 73.8 | 3.814 | 1 | 0.051 | 1 | DGKH | 0 |
| rs10492436 | 13 | 42768238 | t | c | 0.126 | 0.015 | 2.87E-17 | ++ | 0 | 0.954 | 1 | 0.329 | 1 | DGKH | 0 |
| rs12584515 | 13 | 42768456 | a | g | 0.127 | 0.015 | 1.20E-17 | ++ | 73.8 | 3.814 | 1 | 0.051 | 1 | DGKH | 0 |
| rs12584544 | 13 | 42768720 | a | g | 0.127 | 0.015 | 1.20E-17 | ++ | 73.8 | 3.814 | 1 | 0.051 | 1 | DGKH | 0 |
| rs71429407 | 13 | 42769253 | a | g | -0.127 | 0.015 | 1.20E-17 | -- | 73.8 | 3.814 | 1 | 0.051 | 1 | DGKH | 0 |
| rs71429408 | 13 | 42769267 | a | t | 0.127 | 0.015 | 1.20E-17 | ++ | 73.8 | 3.814 | 1 | 0.051 | 1 | DGKH | 0 |
| rs34697334 | 13 | 42769461 | c | g | 0.128 | 0.015 | 3.15E-17 | ++ | 69.4 | 3.270 | 1 | 0.071 | 1 | DGKH | 0 |
| rs7983070 | 13 | 42769494 | t | c | -0.127 | 0.015 | 1.20E-17 | -- | 73.8 | 3.814 | 1 | 0.051 | 1 | DGKH | 0 |
| rs7986154 | 13 | 42769597 | t | c | 0.127 | 0.015 | 1.20E-17 | ++ | 73.8 | 3.814 | 1 | 0.051 | 1 | DGKH | 0 |
| rs60095226 | 13 | 42770097 | t | c | -0.127 | 0.015 | 1.20E-17 | -- | 73.8 | 3.814 | 1 | 0.051 | 1 | DGKH | 0 |
| rs59505610 | 13 | 42770133 | a | g | -0.127 | 0.015 | 1.20E-17 | -- | 73.8 | 3.814 | 1 | 0.051 | 1 | DGKH | 0 |
| rs57753179 | 13 | 42770333 | t | c | -0.127 | 0.015 | 1.20E-17 | -- | 73.8 | 3.814 | 1 | 0.051 | 1 | DGKH | 0 |
| rs56903934 | 13 | 42770507 | t | c | 0.127 | 0.015 | 1.20E-17 | ++ | 73.8 | 3.814 | 1 | 0.051 | 1 | DGKH | 0 |
| rs11617266 | 13 | 42770696 | a | g | -0.120 | 0.022 | 2.43E-08 | -- | 0 | 0.138 | 1 | 0.711 | 1 | DGKH | 0 |
| rs60039389 | 13 | 42770761 | t | c | -0.127 | 0.015 | 1.20E-17 | -- | 73.8 | 3.814 | 1 | 0.051 | 1 | DGKH | 0 |
| rs34816851 | 13 | 42770817 | t | c | 0.127 | 0.015 | 1.61E-17 | ++ | 74.7 | 3.947 | 1 | 0.047 | 1 | DGKH | 0 |

| | | | | | | | | | | | | | | | |
|---|---|---|---|---|---|---|---|---|---|---|---|---|---|---|---|
| rs12863254 | 13 | 42771231 | a | c | 0.127 | 0.015 | 1.20E-17 | ++ | 73.8 | 3.814 | 1 | 0.051 | 1 | DGKH | 0 |
| rs12864038 | 13 | 42771294 | a | g | 0.127 | 0.015 | 1.20E-17 | ++ | 73.8 | 3.814 | 1 | 0.051 | 1 | DGKH | 0 |
| rs12863619 | 13 | 42771310 | t | c | 0.127 | 0.015 | 1.20E-17 | ++ | 73.8 | 3.814 | 1 | 0.051 | 1 | DGKH | 0 |
| rs7999553 | 13 | 42771871 | t | g | -0.084 | 0.015 | 1.88E-08 | -- | 0 | 0.092 | 1 | 0.762 | 1 | DGKH | 0 |
| rs35699135 | 13 | 42772129 | a | g | 0.128 | 0.015 | 8.98E-18 | ++ | 76.3 | 4.219 | 1 | 0.040 | 1 | DGKH | 0 |
| rs35164787 | 13 | 42772133 | a | t | -0.128 | 0.015 | 8.98E-18 | -- | 76.3 | 4.219 | 1 | 0.040 | 1 | DGKH | 0 |
| rs17598799 | 13 | 42772222 | c | g | -0.127 | 0.015 | 1.61E-17 | -- | 19 | 1.235 | 1 | 0.267 | 1 | DGKH | 0 |
| rs9533031 | 13 | 42772717 | t | g | -0.095 | 0.016 | 1.02E-09 | -- | 0 | 0.037 | 1 | 0.847 | 1 | DGKH | 0 |
| rs9566937 | 13 | 42773061 | a | g | -0.120 | 0.022 | 2.78E-08 | -- | 0 | 0.121 | 1 | 0.728 | 1 | DGKH | 0 |
| rs12873956 | 13 | 42773103 | t | c | 0.127 | 0.015 | 1.20E-17 | ++ | 73.8 | 3.814 | 1 | 0.051 | 1 | DGKH | 0 |
| rs9566938 | 13 | 42773322 | a | c | 0.120 | 0.022 | 2.78E-08 | ++ | 0 | 0.121 | 1 | 0.728 | 1 | DGKH | 0 |
| rs12583267 | 13 | 42775439 | a | g | -0.117 | 0.015 | 3.30E-15 | -- | 69.8 | 3.306 | 1 | 0.069 | 1 | DGKH | 0 |
| rs1170100 | 13 | 42775491 | t | c | 0.109 | 0.019 | 2.26E-08 | ++ | 0 | 0.054 | 1 | 0.817 | 1 | DGKH | 0 |
| rs1170102 | 13 | 42776117 | a | g | 0.126 | 0.015 | 2.15E-17 | ++ | 75.5 | 4.082 | 1 | 0.043 | 1 | DGKH | 0 |
| rs1170103 | 13 | 42777027 | c | g | 0.126 | 0.015 | 2.15E-17 | ++ | 75.5 | 4.082 | 1 | 0.043 | 1 | DGKH | 0 |
| rs1170106 | 13 | 42778280 | a | c | -0.126 | 0.015 | 2.87E-17 | -- | 19 | 1.235 | 1 | 0.267 | 1 | DGKH | 0 |
| rs1170107 | 13 | 42778448 | c | g | -0.126 | 0.015 | 2.87E-17 | -- | 19 | 1.235 | 1 | 0.267 | 1 | DGKH | 0 |
| rs1037270 | 13 | 42779257 | c | g | 0.125 | 0.015 | 3.83E-17 | ++ | 23.7 | 1.311 | 1 | 0.252 | 1 | DGKH | 0 |
| rs1037271 | 13 | 42779410 | t | c | -0.133 | 0.016 | 1.24E-17 | -- | 61.3 | 2.583 | 1 | 0.108 | 1 | DGKH | 0 |
| rs1170109 | 13 | 42779694 | t | g | 0.125 | 0.015 | 3.83E-17 | ++ | 75.5 | 4.082 | 1 | 0.043 | 1 | DGKH | 0 |
| rs9562385 | 13 | 42780628 | a | c | -0.121 | 0.016 | 7.36E-15 | -- | 16.3 | 1.194 | 1 | 0.275 | 1 | DGKH | 0 |
| rs8001980 | 13 | 42781012 | t | c | 0.121 | 0.015 | 3.68E-16 | ++ | 13.9 | 1.161 | 1 | 0.281 | 1 | DGKH | 0 |
| rs8001077 | 13 | 42781262 | t | c | -0.111 | 0.017 | 2.03E-11 | -- | 0 | 0.927 | 1 | 0.336 | 1 | DGKH | 0 |
| rs347409 | 13 | 42781631 | t | g | -0.125 | 0.015 | 3.83E-17 | -- | 75.5 | 4.082 | 1 | 0.043 | 1 | DGKH | 0 |
| rs347410 | 13 | 42781703 | t | c | 0.125 | 0.015 | 3.83E-17 | ++ | 75.5 | 4.082 | 1 | 0.043 | 1 | DGKH | 0 |
| rs67354544 | 13 | 42781707 | t | c | 0.124 | 0.015 | 6.79E-17 | ++ | 73.8 | 3.814 | 1 | 0.051 | 1 | DGKH | 0 |
| rs9590677 | 13 | 42781855 | a | g | -0.111 | 0.016 | 9.65E-13 | -- | 0 | 0.595 | 1 | 0.441 | 1 | DGKH | 0 |
| rs9566939 | 13 | 42782887 | a | c | -0.121 | 0.015 | 3.68E-16 | -- | 13.9 | 1.161 | 1 | 0.281 | 1 | DGKH | 0 |
| rs76416563 | 13 | 42785874 | c | g | -0.124 | 0.015 | 6.79E-17 | -- | 73.8 | 3.814 | 1 | 0.051 | 1 | DGKH | 0 |
| rs9533035 | 13 | 42786040 | a | t | 0.121 | 0.022 | 3.39E-08 | ++ | 0 | 0.169 | 1 | 0.681 | 1 | DGKH | 0 |
| rs9533037 | 13 | 42786873 | a | g | 0.121 | 0.022 | 3.39E-08 | ++ | 0 | 0.169 | 1 | 0.681 | 1 | DGKH | 0 |
| rs347413 | 13 | 42787040 | a | g | 0.125 | 0.015 | 3.83E-17 | ++ | 75.5 | 4.082 | 1 | 0.043 | 1 | DGKH | 0 |
| rs347403 | 13 | 42789366 | a | g | 0.127 | 0.019 | 3.45E-11 | ++ | 58.1 | 2.388 | 1 | 0.122 | 1 | DGKH | 0 |
| rs35321674 | 13 | 42796125 | t | g | -0.092 | 0.016 | 4.06E-09 | -- | 72.2 | 3.596 | 1 | 0.058 | 1 | DGKH | 0 |
| rs73543320 | 16 | 20372696 | a | g | 0.121 | 0.021 | 3.62E-09 | ++ | 77.3 | 4.397 | 1 | 0.036 | 1 | PDILT | 0 |
| rs62034970 | 16 | 20383049 | c | g | 0.109 | 0.018 | 7.01E-10 | ++ | 88.4 | 8.653 | 1 | 0.003 | 1 | PDILT | 0 |
| rs4408552 | 16 | 20383531 | a | g | -0.110 | 0.017 | 2.04E-10 | -- | 88.5 | 8.663 | 1 | 0.003 | 1 | PDILT | 0 |
| rs34495073 | 16 | 20385144 | t | c | -0.110 | 0.017 | 2.04E-10 | -- | 88.5 | 8.663 | 1 | 0.003 | 1 | PDILT | 0 |
| rs34532024 | 16 | 20385148 | c | g | -0.110 | 0.017 | 2.04E-10 | -- | 88.5 | 8.663 | 1 | 0.003 | 1 | PDILT | 0 |
| rs35449439 | 16 | 20385182 | c | g | 0.110 | 0.017 | 2.04E-10 | ++ | 88.5 | 8.663 | 1 | 0.003 | 1 | PDILT | 0 |
| rs78751740 | 16 | 20386162 | a | g | 0.122 | 0.019 | 3.62E-10 | ++ | 85.5 | 6.876 | 1 | 0.009 | 1 | PDILT | 0 |
| rs35208507 | 16 | 20388929 | a | g | -0.110 | 0.017 | 2.04E-10 | -- | 88.5 | 8.663 | 1 | 0.003 | 1 | PDILT | 0 |
| rs77924615 | 16 | 20392332 | a | g | 0.111 | 0.018 | 4.08E-10 | ++ | 88.2 | 8.487 | 1 | 0.004 | 1 | PDILT | 0 |
| rs62034975 | 16 | 20392415 | c | g | 0.112 | 0.017 | 1.15E-10 | ++ | 88.7 | 8.833 | 1 | 0.003 | 1 | PDILT | 0 |
| rs79746097 | 16 | 20392810 | t | c | -0.112 | 0.017 | 1.15E-10 | -- | 88.7 | 8.833 | 1 | 0.003 | 1 | PDILT | 0 |
| rs35747824 | 16 | 20393308 | a | t | -0.112 | 0.017 | 1.15E-10 | -- | 88.7 | 8.833 | 1 | 0.003 | 1 | PDILT | 0 |
| rs12921916 | 16 | 20407196 | t | c | -0.114 | 0.018 | 2.99E-10 | -- | 87.7 | 8.154 | 1 | 0.004 | 1 | PDILT | 0 |
| rs7206790 | 16 | 53797908 | c | g | -0.127 | 0.021 | 1.08E-09 | -- | 0 | 0.037 | 1 | 0.848 | 1 | FTO | 0 |
| rs28429148 | 16 | 53798319 | a | g | 0.104 | 0.018 | 7.90E-09 | ++ | 0 | 0.480 | 1 | 0.488 | 1 | FTO | 0 |
| rs8047587 | 16 | 53798622 | t | g | 0.102 | 0.018 | 1.79E-08 | ++ | 0 | 0.443 | 1 | 0.506 | 1 | FTO | 0 |
| rs9940128 | 16 | 53800754 | a | g | 0.093 | 0.017 | 4.25E-08 | ++ | 0 | 0.170 | 1 | 0.680 | 1 | FTO | 0 |
| rs1421085 | 16 | 53800954 | t | c | -0.105 | 0.018 | 6.14E-09 | -- | 0 | 0.130 | 1 | 0.719 | 1 | FTO | 0 |
| rs1421086 | 16 | 53801343 | a | c | 0.093 | 0.017 | 4.25E-08 | ++ | 0 | 0.170 | 1 | 0.680 | 1 | FTO | 0 |
| rs9923147 | 16 | 53801549 | t | c | 0.094 | 0.017 | 3.60E-08 | ++ | 0 | 0.147 | 1 | 0.702 | 1 | FTO | 0 |
| rs9923544 | 16 | 53801985 | t | c | 0.093 | 0.017 | 4.25E-08 | ++ | 0 | 0.170 | 1 | 0.680 | 1 | FTO | 0 |
| rs11642015 | 16 | 53802494 | t | c | 0.104 | 0.018 | 7.29E-09 | ++ | 0 | 0.111 | 1 | 0.739 | 1 | FTO | 0 |
| rs1558901 | 16 | 53803187 | a | t | -0.094 | 0.017 | 3.60E-08 | -- | 0 | 0.147 | 1 | 0.702 | 1 | FTO | 0 |
| rs62048402 | 16 | 53803223 | a | g | 0.105 | 0.018 | 6.14E-09 | ++ | 0 | 0.130 | 1 | 0.719 | 1 | FTO | 0 |
| rs1558902 | 16 | 53803574 | a | t | 0.105 | 0.018 | 6.14E-09 | ++ | 0 | 0.130 | 1 | 0.719 | 1 | FTO | 0 |
| rs11075985 | 16 | 53805207 | a | c | 0.094 | 0.017 | 3.60E-08 | ++ | 0 | 0.147 | 1 | 0.702 | 1 | FTO | 0 |
| rs56094641 | 16 | 53806453 | a | g | -0.105 | 0.018 | 6.14E-09 | -- | 0 | 0.130 | 1 | 0.719 | 1 | FTO | 0 |
| rs55872725 | 16 | 53809123 | t | c | 0.105 | 0.018 | 5.16E-09 | ++ | 0 | 0.151 | 1 | 0.698 | 1 | FTO | 0 |
| rs1121980 | 16 | 53809247 | a | g | 0.094 | 0.017 | 3.60E-08 | ++ | 0 | 0.147 | 1 | 0.702 | 1 | FTO | 0 |
| rs7187250 | 16 | 53810546 | a | c | 0.102 | 0.018 | 1.38E-08 | ++ | 0 | 0.151 | 1 | 0.698 | 1 | FTO | 0 |
| rs7193144 | 16 | 53810686 | t | c | -0.101 | 0.018 | 2.23E-08 | -- | 0 | 0.173 | 1 | 0.678 | 1 | FTO | 0 |
| rs62033399 | 16 | 53810943 | t | c | 0.101 | 0.018 | 2.23E-08 | ++ | 0 | 0.173 | 1 | 0.678 | 1 | FTO | 0 |
| rs62033400 | 16 | 53811788 | a | g | -0.100 | 0.018 | 2.63E-08 | -- | 0 | 0.151 | 1 | 0.698 | 1 | FTO | 0 |
| rs8063057 | 16 | 53812433 | t | c | -0.101 | 0.018 | 2.23E-08 | -- | 0 | 0.173 | 1 | 0.678 | 1 | FTO | 0 |
| rs8057044 | 16 | 53812614 | a | g | 0.095 | 0.017 | 4.24E-08 | ++ | 0 | 0.301 | 1 | 0.584 | 1 | FTO | 0 |
| rs17817449 | 16 | 53813367 | t | g | -0.100 | 0.018 | 2.63E-08 | -- | 0 | 0.151 | 1 | 0.698 | 1 | FTO | 0 |
| rs8043757 | 16 | 53813450 | a | t | -0.100 | 0.018 | 2.63E-08 | -- | 0 | 0.151 | 1 | 0.698 | 1 | FTO | 0 |
| rs9972653 | 16 | 53814363 | t | g | 0.101 | 0.018 | 2.23E-08 | ++ | 0 | 0.173 | 1 | 0.678 | 1 | FTO | 0 |
| rs17817497 | 16 | 53815435 | t | c | -0.101 | 0.018 | 2.23E-08 | -- | 0 | 0.173 | 1 | 0.678 | 1 | FTO | 0 |
| rs8050136 | 16 | 53816275 | a | c | 0.101 | 0.018 | 2.23E-08 | ++ | 0 | 0.173 | 1 | 0.678 | 1 | FTO | 0 |
| rs8051591 | 16 | 53816752 | a | g | -0.101 | 0.018 | 2.23E-08 | -- | 0 | 0.173 | 1 | 0.678 | 1 | FTO | 0 |
| rs9935401 | 16 | 53816838 | a | g | 0.100 | 0.018 | 2.63E-08 | ++ | 0 | 0.151 | 1 | 0.698 | 1 | FTO | 0 |
| rs3751812 | 16 | 53818460 | t | g | 0.099 | 0.018 | 2.52E-08 | ++ | 0 | 0.097 | 1 | 0.756 | 1 | FTO | 0 |
| rs3751814 | 16 | 53818724 | a | g | 0.099 | 0.018 | 2.52E-08 | ++ | 0 | 0.097 | 1 | 0.756 | 1 | FTO | 0 |
| rs56313538 | 16 | 53818834 | a | g | -0.098 | 0.018 | 2.96E-08 | -- | 0 | 0.080 | 1 | 0.777 | 1 | FTO | 0 |
| rs9936385 | 16 | 53819169 | t | c | -0.098 | 0.018 | 2.96E-08 | -- | 0 | 0.080 | 1 | 0.777 | 1 | FTO | 0 |
| rs9923233 | 16 | 53819198 | c | g | 0.098 | 0.018 | 2.96E-08 | ++ | 0 | 0.080 | 1 | 0.777 | 1 | FTO | 0 |
| rs9923312 | 16 | 53819367 | a | g | 0.099 | 0.018 | 2.52E-08 | ++ | 0 | 0.097 | 1 | 0.756 | 1 | FTO | 0 |
| rs11075988 | 16 | 53819771 | a | g | 0.098 | 0.018 | 2.96E-08 | ++ | 0 | 0.080 | 1 | 0.777 | 1 | FTO | 0 |
| rs11075989 | 16 | 53819877 | t | c | 0.099 | 0.018 | 2.52E-08 | ++ | 0 | 0.097 | 1 | 0.756 | 1 | FTO | 0 |
| rs11075990 | 16 | 53819893 | a | g | -0.099 | 0.018 | 2.52E-08 | -- | 0 | 0.097 | 1 | 0.756 | 1 | FTO | 0 |
| rs11075991 | 16 | 53819937 | a | t | -0.099 | 0.018 | 2.52E-08 | -- | 0 | 0.097 | 1 | 0.756 | 1 | FTO | 0 |
| rs11075992 | 16 | 53820066 | t | c | -0.099 | 0.018 | 2.52E-08 | -- | 0 | 0.097 | 1 | 0.756 | 1 | FTO | 0 |
| rs9926289 | 16 | 53820503 | a | g | 0.099 | 0.018 | 2.52E-08 | ++ | 0 | 0.097 | 1 | 0.756 | 1 | FTO | 0 |
| rs9939609 | 16 | 53820527 | a | t | 0.099 | 0.018 | 2.52E-08 | ++ | 0 | 0.097 | 1 | 0.756 | 1 | FTO | 0 |
| rs9927317 | 16 | 53820996 | c | g | -0.100 | 0.018 | 3.10E-08 | -- | 0 | 0.130 | 1 | 0.719 | 1 | FTO | 0 |
| rs17817712 | 16 | 53821125 | a | g | -0.099 | 0.018 | 2.52E-08 | -- | 0 | 0.097 | 1 | 0.756 | 1 | FTO | 0 |
| rs7206410 | 16 | 53821297 | t | c | -0.099 | 0.018 | 2.52E-08 | -- | 0 | 0.097 | 1 | 0.756 | 1 | FTO | 0 |
| rs7202116 | 16 | 53821615 | a | g | -0.099 | 0.018 | 4.31E-08 | -- | 0 | 0.093 | 1 | 0.760 | 1 | FTO | 0 |
| rs7202296 | 16 | 53821690 | a | g | -0.099 | 0.018 | 4.31E-08 | -- | 0 | 0.093 | 1 | 0.760 | 1 | FTO | 0 |
| rs66908032 | 16 | 53822142 | a | c | 0.099 | 0.018 | 4.31E-08 | ++ | 0 | 0.093 | 1 | 0.760 | 1 | FTO | 0 |
| rs72803697 | 16 | 53822183 | t | c | 0.099 | 0.018 | 4.31E-08 | ++ | 0 | 0.093 | 1 | 0.760 | 1 | FTO | 0 |
| rs62033403 | 16 | 53822237 | a | g | -0.099 | 0.018 | 4.31E-08 | -- | 0 | 0.093 | 1 | 0.760 | 1 | FTO | 0 |
| rs62033404 | 16 | 53822239 | a | g | -0.099 | 0.018 | 4.31E-08 | -- | 0 | 0.093 | 1 | 0.760 | 1 | FTO | 0 |
| rs62033405 | 16 | 53822387 | t | c | 0.099 | 0.018 | 4.31E-08 | ++ | 0 | 0.093 | 1 | 0.760 | 1 | FTO | 0 |
| rs7206122 | 16 | 53822440 | a | g | 0.099 | 0.018 | 4.31E-08 | ++ | 0 | 0.093 | 1 | 0.760 | 1 | FTO | 0 |
| rs10468280 | 16 | 53827479 | a | g | -0.098 | 0.018 | 4.94E-08 | -- | 0 | 0.151 | 1 | 0.698 | 1 | FTO | 0 |
| rs62033408 | 16 | 53827962 | a | g | -0.098 | 0.018 | 4.94E-08 | -- | 0 | 0.151 | 1 | 0.698 | 1 | FTO | 0 |
| rs17817964 | 16 | 53828066 | t | c | 0.099 | 0.018 | 3.95E-08 | ++ | 0 | 0.480 | 1 | 0.488 | 1 | FTO | 0 |
| rs62033413 | 16 | 53830055 | c | g | -0.097 | 0.018 | 4.09E-08 | -- | 0 | 0.259 | 1 | 0.611 | 1 | FTO | 0 |
| rs72805611 | 16 | 53831354 | t | c | 0.097 | 0.018 | 4.09E-08 | ++ | 0 | 0.259 | 1 | 0.611 | 1 | FTO | 0 |
| rs72805612 | 16 | 53834608 | a | g | 0.098 | 0.018 | 4.76E-08 | ++ | 0 | 0.307 | 1 | 0.579 | 1 | FTO | 0 |
| rs9907597 | 17 | 59435497 | a | c | -0.094 | 0.016 | 8.97E-09 | -- | 62.3 | 2.655 | 1 | 0.103 | 1 | BCAS3 | 0 |
| rs9905610 | 17 | 59440351 | t | g | 0.090 | 0.016 | 7.23E-09 | ++ | 66.8 | 3.012 | 1 | 0.083 | 1 | BCAS3 | 0 |
| rs1990292 | 17 | 59444758 | a | g | 0.087 | 0.015 | 4.66E-09 | ++ | 67.4 | 3.066 | 1 | 0.080 | 1 | BCAS3 | 0 |
| rs3785842 | 17 | 59446530 | c | g | -0.102 | 0.016 | 5.50E-11 | -- | 71.2 | 3.475 | 1 | 0.062 | 1 | BCAS3 | 0 |
| rs3785841 | 17 | 59446541 | c | g | -0.105 | 0.016 | 1.85E-11 | -- | 62.8 | 2.687 | 1 | 0.101 | 1 | BCAS3 | 0 |

| | | | | | | | | | | | | | | | | | |
|---|---|---|---|---|---|---|---|---|---|---|---|---|---|---|---|---|---|
| rs11653176 | 17 | 59447369 | t | c | -0.090 | 0.016 | 1.57E-08 | -- | 81 | 5.261 | 1 | 0.022 | 1 | BCAS3 | 0 | | |
| rs7217891 | 17 | 59447984 | a | g | 0.116 | 0.018 | 5.31E-11 | ++ | 73 | 3.699 | 1 | 0.054 | 1 | BCAS3 | 0 | | |
| rs1010269 | 17 | 59448945 | a | g | -0.099 | 0.016 | 5.45E-10 | -- | 85.3 | 6.801 | 1 | 0.009 | 1 | BCAS3 | 0 | | |
| rs4968556 | 17 | 59449082 | t | c | 0.098 | 0.016 | 8.26E-10 | ++ | 86 | 7.132 | 1 | 0.008 | 1 | BCAS3 | 0 | | |
| rs11657044 | 17 | 59450105 | t | c | -0.095 | 0.015 | 3.77E-10 | -- | 85.5 | 6.919 | 1 | 0.009 | 1 | BCAS3 | 0 | | |
| rs9895661 | 17 | 59456589 | t | c | 0.126 | 0.017 | 1.41E-13 | ++ | 75.8 | 4.133 | 1 | 0.042 | 1 | BCAS3 | 0 | | |
| rs2079742 | 17 | 59465697 | t | c | 0.134 | 0.018 | 4.29E-14 | ++ | 61.5 | 2.599 | 1 | 0.107 | 1 | BCAS3 | 0 | | |
| rs11079428 | 17 | 59466701 | a | t | 0.113 | 0.018 | 4.11E-10 | ++ | 73.4 | 3.766 | 1 | 0.052 | 1 | BCAS3 | 0 | | |
| rs3785837 | 17 | 59468942 | a | g | 0.114 | 0.018 | 2.84E-10 | ++ | 71.9 | 3.554 | 1 | 0.059 | 1 | BCAS3 | 0 | | |
| rs2286527 | 17 | 59471686 | a | g | 0.104 | 0.017 | 8.89E-10 | ++ | 76.5 | 4.253 | 1 | 0.039 | 1 | LOC645722 | 0 | | |
| rs2286526 | 17 | 59472057 | t | c | 0.112 | 0.016 | 6.04E-13 | ++ | 25.3 | 1.339 | 1 | 0.247 | 1 | LOC645722 | 0 | | |
| rs2286525 | 17 | 59472123 | a | g | 0.106 | 0.016 | 8.76E-11 | ++ | 65 | 2.859 | 1 | 0.091 | 1 | LOC645722 | 0 | | |
| rs11079429 | 17 | 59472403 | a | g | 0.103 | 0.016 | 4.43E-11 | ++ | 56.2 | 2.282 | 1 | 0.131 | 1 | LOC645722 | 0 | | |
| rs1000423 | 17 | 59475642 | t | c | 0.102 | 0.015 | 2.13E-11 | ++ | 15.1 | 1.177 | 1 | 0.278 | 1 | LOC645722 | 0 | | |
| rs2109019 | 17 | 59475888 | a | c | -0.087 | 0.016 | 2.69E-08 | -- | 65.5 | 2.902 | 1 | 0.088 | 1 | LOC645722 | 0 | | |
| rs62071306 | 17 | 59476066 | a | c | -0.107 | 0.016 | 5.82E-11 | -- | 0 | 0.795 | 1 | 0.373 | 1 | LOC645722 | 0 | | |
| rs1476781 | 17 | 59476415 | t | c | -0.087 | 0.016 | 2.69E-08 | -- | 65.5 | 2.902 | 1 | 0.088 | 1 | LOC645722 | 0 | | |
| rs34446110 | 17 | 59478354 | c | g | 0.098 | 0.015 | 4.12E-11 | ++ | 50 | 2.000 | 1 | 0.157 | 2 | LOC645722 | 0 | TBX2 | 0 |
| rs12952625 | 17 | 59478644 | a | g | 0.099 | 0.015 | 8.73E-11 | ++ | 45 | 1.817 | 1 | 0.178 | 2 | LOC645722 | 0 | TBX2 | 0 |
| rs2270114 | 17 | 59478776 | c | g | 0.099 | 0.015 | 8.73E-11 | ++ | 45 | 1.817 | 1 | 0.178 | 2 | LOC645722 | 0 | TBX2 | 0 |
| rs9891115 | 17 | 59482173 | a | g | 0.106 | 0.017 | 1.97E-10 | ++ | 27.4 | 1.376 | 1 | 0.241 | 2 | LOC645722 | 0 | TBX2 | 0 |
| rs8068318 | 17 | 59483766 | t | c | 0.098 | 0.015 | 4.12E-11 | ++ | 44.9 | 1.814 | 1 | 0.178 | 2 | LOC645722 | 0 | TBX2 | 0 |
| rs7215775 | 17 | 59484316 | a | g | -0.107 | 0.015 | 7.39E-13 | -- | 8.2 | 1.090 | 1 | 0.297 | 2 | LOC645722 | 0 | TBX2 | 0 |
| rs8073698 | 17 | 59484833 | t | c | 0.098 | 0.015 | 4.12E-11 | ++ | 44.9 | 1.814 | 1 | 0.178 | 2 | LOC645722 | 0 | TBX2 | 0 |
| rs8074151 | 17 | 59485017 | a | g | 0.099 | 0.015 | 3.28E-11 | ++ | 47.5 | 1.906 | 1 | 0.167 | 2 | LOC645722 | 0 | TBX2 | 0 |
| rs8078036 | 17 | 59485120 | a | g | 0.099 | 0.015 | 3.28E-11 | ++ | 47.5 | 1.906 | 1 | 0.167 | 2 | LOC645722 | 0 | TBX2 | 0 |
| rs2240736 | 17 | 59485393 | t | c | 0.108 | 0.015 | 4.51E-13 | ++ | 8.2 | 1.090 | 1 | 0.297 | 2 | LOC645722 | 0 | TBX2 | 0 |
| rs1057987 | 17 | 59485555 | t | c | 0.099 | 0.015 | 3.28E-11 | ++ | 47.5 | 1.906 | 1 | 0.167 | 2 | LOC645722 | 0 | TBX2 | 0 |
| rs3198587 | 17 | 59486624 | c | g | 0.099 | 0.018 | 2.52E-08 | ++ | 58.7 | 2.420 | 1 | 0.120 | 2 | LOC645722 | 0 | TBX2 | 0 |
| rs1058004 | 17 | 59486732 | t | c | 0.099 | 0.015 | 3.28E-11 | ++ | 47.5 | 1.906 | 1 | 0.167 | 2 | LOC645722 | 0 | TBX2 | 0 |
| rs729781 | 17 | 59486799 | a | g | -0.099 | 0.015 | 3.28E-11 | -- | 47.5 | 1.906 | 1 | 0.167 | 2 | LOC645722 | 0 | TBX2 | 0 |
| rs11871637 | 17 | 59490235 | t | c | 0.098 | 0.015 | 4.12E-11 | ++ | 2 | 1.020 | 1 | 0.312 | 1 | C17orf82 | 0 | | |
| rs11868532 | 17 | 59496083 | t | c | -0.096 | 0.015 | 1.27E-10 | -- | 35.6 | 1.552 | 1 | 0.213 | 1 | C17orf82 | 5442 | | |
| rs9905140 | 17 | 59496238 | t | c | -0.095 | 0.015 | 1.58E-10 | -- | 31.9 | 1.469 | 1 | 0.225 | 1 | C17orf82 | 5597 | | |
| rs10853031 | 17 | 59497487 | c | g | -0.089 | 0.016 | 1.28E-08 | -- | 42.4 | 1.737 | 1 | 0.188 | 1 | C17orf82 | 6846 | | |
| rs60984983 | 19 | 14553265 | t | c | -0.124 | 0.019 | 3.31E-11 | -- | 20.4 | 1.256 | 1 | 0.263 | 1 | PKN1 | 0 | | |
| rs3786654 | 19 | 14557821 | a | g | -0.115 | 0.017 | 1.23E-11 | -- | 0.3 | 1.003 | 1 | 0.317 | 1 | PKN1 | 0 | | |
| rs3786652 | 19 | 14560439 | t | c | -0.117 | 0.017 | 6.66E-12 | -- | 0 | 0.945 | 1 | 0.331 | 1 | PKN1 | 0 | | |
| rs12973696 | 19 | 14562977 | t | c | -0.117 | 0.017 | 6.66E-12 | -- | 0 | 0.945 | 1 | 0.331 | 1 | PKN1 | 0 | | |
| rs12975525 | 19 | 14563856 | a | c | -0.117 | 0.017 | 6.66E-12 | -- | 0 | 0.834 | 1 | 0.361 | 1 | PKN1 | 0 | | |
| rs11673475 | 19 | 14566227 | a | c | -0.118 | 0.017 | 4.40E-12 | -- | 0 | 0.834 | 1 | 0.361 | 1 | PKN1 | 0 | | |
| rs11085894 | 19 | 14569623 | c | g | -0.092 | 0.016 | 1.84E-08 | -- | 59.3 | 2.458 | 1 | 0.117 | 1 | PKN1 | 0 | | |
| rs74956940 | 19 | 14571966 | c | g | -0.122 | 0.017 | 6.53E-13 | -- | 0 | 0.347 | 1 | 0.556 | 1 | PKN1 | 0 | | |
| rs3826758 | 19 | 14572497 | a | c | 0.086 | 0.016 | 3.88E-08 | ++ | 0 | 0.232 | 1 | 0.630 | 1 | PKN1 | 0 | | |
| rs2420538 | 19 | 14573654 | a | g | -0.114 | 0.016 | 2.39E-12 | -- | 0 | 0.306 | 1 | 0.580 | 1 | PKN1 | 0 | | |
| rs1363403 | 19 | 14573982 | a | t | -0.085 | 0.016 | 4.66E-08 | -- | 0 | 0.202 | 1 | 0.653 | 1 | PKN1 | 0 | | |
| rs34422078 | 19 | 14578354 | c | g | -0.117 | 0.017 | 2.18E-12 | -- | 0 | 0.293 | 1 | 0.588 | 1 | PKN1 | 0 | | |
| rs2241364 | 19 | 14583064 | c | g | -0.089 | 0.016 | 1.06E-08 | -- | 66.8 | 3.012 | 1 | 0.083 | 1 | PTGER1 | 214 | | |
| rs28364040 | 19 | 14585746 | t | c | -0.119 | 0.017 | 8.79E-13 | -- | 0 | 0.177 | 1 | 0.674 | 1 | PTGER1 | 0 | | |
| rs2241360 | 19 | 14586015 | t | c | -0.096 | 0.016 | 1.65E-09 | -- | 0 | 0.830 | 1 | 0.362 | 1 | PTGER1 | 0 | | |
| rs2241359 | 19 | 14586245 | a | g | -0.090 | 0.016 | 7.23E-09 | -- | 58 | 2.380 | 1 | 0.123 | 1 | PTGER1 | -71 | | |
| rs3810253 | 19 | 14587069 | a | c | 0.096 | 0.016 | 1.62E-09 | ++ | 0 | 0.720 | 1 | 0.396 | 1 | PTGER1 | -895 | | |
| rs10416814 | 19 | 14587202 | a | t | -0.096 | 0.016 | 1.62E-09 | -- | 0 | 0.720 | 1 | 0.396 | 1 | PTGER1 | -1028 | | |
| rs3760702 | 19 | 14588237 | a | g | 0.120 | 0.017 | 1.54E-12 | ++ | 0 | 0.087 | 1 | 0.768 | 1 | GIPC1 | 334 | | |
| rs2241358 | 19 | 14588476 | c | g | -0.120 | 0.017 | 5.45E-13 | -- | 0 | 0.090 | 1 | 0.764 | 1 | GIPC1 | 95 | | |
| rs76301301 | 19 | 14589068 | t | g | -0.120 | 0.017 | 1.54E-12 | -- | 0 | 0.281 | 1 | 0.596 | 1 | GIPC1 | 0 | | |
| rs2241357 | 19 | 14590919 | a | g | 0.120 | 0.017 | 1.54E-12 | ++ | 0 | 0.222 | 1 | 0.637 | 1 | GIPC1 | 0 | | |
| rs60438220 | 19 | 14593176 | a | t | -0.125 | 0.018 | 1.54E-12 | -- | 0 | 0.115 | 1 | 0.734 | 1 | GIPC1 | 0 | | |
| rs1543703 | 20 | 52699335 | a | t | -0.086 | 0.016 | 3.23E-08 | -- | 39.5 | 1.653 | 1 | 0.199 | 1 | BCAS1 | -12031 | | |
| rs6064045 | 20 | 52701293 | t | c | -0.086 | 0.016 | 3.23E-08 | -- | 39.5 | 1.653 | 1 | 0.199 | 1 | BCAS1 | -13989 | | |
| rs6068776 | 20 | 52702494 | a | g | -0.087 | 0.016 | 2.69E-08 | -- | 42.4 | 1.737 | 1 | 0.188 | 1 | BCAS1 | -15190 | | |
| rs13041834 | 20 | 52703284 | t | c | 0.087 | 0.016 | 2.69E-08 | ++ | 42.4 | 1.737 | 1 | 0.188 | 1 | BCAS1 | -15980 | | |
| rs6123359 | 20 | 52714706 | a | g | -0.083 | 0.015 | 2.28E-08 | -- | 0 | 0.889 | 1 | 0.346 | 1 | BCAS1 | -27402 | | |
| rs4817797 | 21 | 37801396 | a | g | 0.081 | 0.015 | 4.90E-08 | ++ | 0 | 0.073 | 1 | 0.788 | 1 | CHAF1B | 12271 | | |
| rs2835345 | 21 | 37801434 | a | c | 0.082 | 0.015 | 3.35E-08 | ++ | 0 | 0.113 | 1 | 0.736 | 1 | CHAF1B | 12309 | | |
| rs218641 | 21 | 37802363 | a | g | 0.091 | 0.015 | 1.10E-09 | ++ | 0 | 0.192 | 1 | 0.662 | 1 | CHAF1B | 13238 | | |
| rs218642 | 21 | 37802629 | t | c | -0.090 | 0.015 | 2.73E-09 | -- | 0 | 0.053 | 1 | 0.818 | 1 | CHAF1B | 13504 | | |
| rs55977638 | 21 | 37804666 | a | c | 0.091 | 0.015 | 1.10E-09 | ++ | 0 | 0.137 | 1 | 0.711 | 1 | CHAF1B | 15541 | | |
| rs218644 | 21 | 37805228 | a | t | -0.089 | 0.015 | 2.52E-09 | -- | 0 | 0.028 | 1 | 0.866 | 1 | CHAF1B | 16103 | | |
| rs2835348 | 21 | 37812899 | a | g | -0.094 | 0.015 | 3.04E-10 | -- | 0 | 0.192 | 1 | 0.662 | 1 | CLDN14 | 20020 | | |
| rs2835349 | 21 | 37814114 | t | c | 0.094 | 0.015 | 2.45E-10 | ++ | 0 | 0.222 | 1 | 0.637 | 1 | CLDN14 | 18805 | | |
| rs9975445 | 21 | 37814666 | c | g | 0.094 | 0.015 | 2.45E-10 | ++ | 0 | 0.222 | 1 | 0.637 | 1 | CLDN14 | 18253 | | |
| rs2835352 | 21 | 37815013 | a | g | -0.094 | 0.015 | 2.45E-10 | -- | 0 | 0.222 | 1 | 0.637 | 1 | CLDN14 | 17906 | | |
| rs2633330 | 21 | 37815550 | t | g | -0.100 | 0.015 | 4.31E-11 | -- | 0 | 0.243 | 1 | 0.622 | 1 | CLDN14 | 17369 | | |
| rs2776287 | 21 | 37816185 | a | g | -0.083 | 0.015 | 2.76E-08 | -- | 0 | 0.409 | 1 | 0.522 | 1 | CLDN14 | 16734 | | |
| rs8128976 | 21 | 37816257 | a | g | -0.087 | 0.016 | 4.75E-08 | -- | 0 | 0.949 | 1 | 0.330 | 1 | CLDN14 | 16662 | | |
| rs2244033 | 21 | 37816274 | t | c | -0.083 | 0.015 | 2.76E-08 | -- | 0 | 0.328 | 1 | 0.567 | 1 | CLDN14 | 16645 | | |
| rs2244038 | 21 | 37816353 | t | c | -0.083 | 0.015 | 2.28E-08 | -- | 0 | 0.367 | 1 | 0.545 | 1 | CLDN14 | 16566 | | |
| rs8129181 | 21 | 37816547 | c | g | -0.083 | 0.015 | 2.28E-08 | -- | 0 | 0.367 | 1 | 0.545 | 1 | CLDN14 | 16372 | | |
| rs2835354 | 21 | 37816824 | a | g | -0.087 | 0.016 | 4.75E-08 | -- | 0 | 0.949 | 1 | 0.330 | 1 | CLDN14 | 16095 | | |
| rs11088349 | 21 | 37816927 | a | g | 0.083 | 0.015 | 2.76E-08 | ++ | 0 | 0.409 | 1 | 0.522 | 1 | CLDN14 | 15992 | | |
| rs9982272 | 21 | 37816947 | a | g | -0.087 | 0.016 | 4.75E-08 | -- | 0 | 0.949 | 1 | 0.330 | 1 | CLDN14 | 15972 | | |
| rs28556960 | 21 | 37817066 | a | t | -0.087 | 0.016 | 4.75E-08 | -- | 0 | 0.949 | 1 | 0.330 | 1 | CLDN14 | 15853 | | |
| rs2835355 | 21 | 37817159 | t | g | 0.087 | 0.016 | 4.75E-08 | ++ | 0 | 0.949 | 1 | 0.330 | 1 | CLDN14 | 15760 | | |
| rs2835356 | 21 | 37817198 | t | g | 0.082 | 0.015 | 3.35E-08 | ++ | 0 | 0.367 | 1 | 0.545 | 1 | CLDN14 | 15721 | | |
| rs2835357 | 21 | 37817268 | t | c | 0.087 | 0.016 | 4.75E-08 | ++ | 0 | 0.949 | 1 | 0.330 | 1 | CLDN14 | 15651 | | |
| rs7282694 | 21 | 37817366 | t | c | 0.082 | 0.015 | 4.05E-08 | ++ | 0 | 0.328 | 1 | 0.567 | 1 | CLDN14 | 15553 | | |
| rs57583267 | 21 | 37817528 | a | t | 0.082 | 0.015 | 4.05E-08 | ++ | 0 | 0.328 | 1 | 0.567 | 1 | CLDN14 | 15391 | | |
| rs4816534 | 21 | 37817565 | a | c | -0.087 | 0.016 | 4.75E-08 | -- | 0 | 0.949 | 1 | 0.330 | 1 | CLDN14 | 15354 | | |
| rs8134233 | 21 | 37819071 | a | g | 0.086 | 0.016 | 3.88E-08 | ++ | 0 | 0.753 | 1 | 0.386 | 1 | CLDN14 | 13848 | | |
| rs73204235 | 21 | 37819218 | a | c | 0.090 | 0.016 | 8.75E-09 | ++ | 0 | 0.299 | 1 | 0.585 | 1 | CLDN14 | 13701 | | |
| rs73204236 | 21 | 37819219 | t | c | 0.090 | 0.016 | 8.75E-09 | ++ | 0 | 0.299 | 1 | 0.585 | 1 | CLDN14 | 13700 | | |
| rs62229441 | 21 | 37819857 | a | c | 0.083 | 0.015 | 2.76E-08 | ++ | 0 | 0.328 | 1 | 0.567 | 1 | CLDN14 | 13062 | | |
| rs113080988 | 21 | 37820781 | a | g | 0.084 | 0.015 | 1.54E-08 | ++ | 0 | 0.222 | 1 | 0.637 | 1 | CLDN14 | 12138 | | |
| rs12482435 | 21 | 37820956 | t | c | 0.085 | 0.015 | 1.04E-08 | ++ | 0 | 0.222 | 1 | 0.637 | 1 | CLDN14 | 11963 | | |
| rs12482423 | 21 | 37820960 | a | g | -0.085 | 0.015 | 1.27E-08 | -- | 0 | 0.255 | 1 | 0.614 | 1 | CLDN14 | 11959 | | |
| rs12482688 | 21 | 37821589 | t | c | -0.085 | 0.015 | 1.27E-08 | -- | 0 | 0.255 | 1 | 0.614 | 1 | CLDN14 | 11330 | | |
| rs12482525 | 21 | 37821597 | a | g | -0.085 | 0.015 | 1.27E-08 | -- | 0 | 0.255 | 1 | 0.614 | 1 | CLDN14 | 11322 | | |
| rs55681268 | 21 | 37821717 | t | g | -0.085 | 0.015 | 1.27E-08 | -- | 0 | 0.255 | 1 | 0.614 | 1 | CLDN14 | 11202 | | |
| rs12627161 | 21 | 37821947 | a | g | 0.085 | 0.015 | 1.27E-08 | ++ | 0 | 0.255 | 1 | 0.614 | 1 | CLDN14 | 10972 | | |
| rs12627241 | 21 | 37821987 | t | c | -0.085 | 0.015 | 1.27E-08 | -- | 0 | 0.255 | 1 | 0.614 | 1 | CLDN14 | 10932 | | |
| rs12627173 | 21 | 37822013 | a | g | 0.085 | 0.015 | 1.27E-08 | ++ | 0 | 0.255 | 1 | 0.614 | 1 | CLDN14 | 10906 | | |
| rs12627147 | 21 | 37822094 | a | g | -0.085 | 0.015 | 1.27E-08 | -- | 0 | 0.255 | 1 | 0.614 | 1 | CLDN14 | 10825 | | |
| rs12627167 | 21 | 37822153 | a | c | 0.085 | 0.015 | 1.27E-08 | ++ | 0 | 0.255 | 1 | 0.614 | 1 | CLDN14 | 10766 | | |
| rs12233341 | 21 | 37822659 | a | t | -0.086 | 0.015 | 6.97E-09 | -- | 0 | 0.367 | 1 | 0.545 | 1 | CLDN14 | 10260 | | |
| rs12233342 | 21 | 37822686 | a | g | 0.095 | 0.017 | 2.57E-08 | ++ | 0 | 0.543 | 1 | 0.461 | 1 | CLDN14 | 10233 | | |
| rs12233346 | 21 | 37822689 | t | c | -0.096 | 0.017 | 1.54E-08 | -- | 0 | 0.500 | 1 | 0.480 | 1 | CLDN14 | 10230 | | |
| rs2835360 | 21 | 37822972 | a | g | 0.084 | 0.015 | 1.54E-08 | ++ | 0 | 0.290 | 1 | 0.590 | 1 | CLDN14 | 9947 | | |
| rs12483650 | 21 | 37823076 | a | g | 0.084 | 0.015 | 1.54E-08 | ++ | 0 | 0.290 | 1 | 0.590 | 1 | CLDN14 | 9843 | | |
| rs12483621 | 21 | 37823096 | t | c | 0.084 | 0.015 | 1.54E-08 | ++ | 0 | 0.290 | 1 | 0.590 | 1 | CLDN14 | 9823 | | |
| rs12481735 | 21 | 37823273 | t | c | -0.093 | 0.017 | 4.25E-08 | -- | 0 | 0.681 | 1 | 0.409 | 1 | CLDN14 | 9646 | | |

| | | | | | | | | | | | | | | | |
|---|---|---|---|---|---|---|---|---|---|---|---|---|---|---|---|
| rs12481748 | 21 | 37823326 | t | c | -0.086 | 0.015 | 6.97E-09 | -- | 0 | 0.290 | 1 | 0.590 | 1 | CLDN14 | 9593 |
| rs58176733 | 21 | 37823432 | t | c | 0.091 | 0.017 | 4.30E-08 | ++ | 0 | 0.566 | 1 | 0.452 | 1 | CLDN14 | 9487 |
| rs12481770 | 21 | 37823437 | t | c | -0.092 | 0.016 | 1.54E-08 | -- | 0 | 0.378 | 1 | 0.539 | 1 | CLDN14 | 9482 |
| rs12483690 | 21 | 37823563 | a | g | -0.085 | 0.015 | 1.27E-08 | -- | 0 | 0.328 | 1 | 0.567 | 1 | CLDN14 | 9356 |
| rs59368922 | 21 | 37823764 | t | c | -0.092 | 0.016 | 1.84E-08 | -- | 0 | 0.591 | 1 | 0.442 | 1 | CLDN14 | 9155 |
| rs12482079 | 21 | 37823769 | a | g | -0.093 | 0.016 | 1.29E-08 | -- | 0 | 0.417 | 1 | 0.519 | 1 | CLDN14 | 9150 |
| rs12482092 | 21 | 37823851 | a | g | -0.085 | 0.015 | 1.04E-08 | -- | 0 | 0.290 | 1 | 0.590 | 1 | CLDN14 | 9068 |
| rs12482097 | 21 | 37823890 | a | g | -0.085 | 0.015 | 1.27E-08 | -- | 0 | 0.328 | 1 | 0.567 | 1 | CLDN14 | 9029 |
| rs12482138 | 21 | 37823906 | t | c | 0.085 | 0.015 | 1.27E-08 | ++ | 0 | 0.328 | 1 | 0.567 | 1 | CLDN14 | 9013 |
| rs12482257 | 21 | 37823958 | t | c | -0.085 | 0.015 | 1.27E-08 | -- | 0 | 0.328 | 1 | 0.567 | 1 | CLDN14 | 8961 |
| rs12482140 | 21 | 37824033 | a | c | -0.085 | 0.015 | 1.27E-08 | -- | 0 | 0.328 | 1 | 0.567 | 1 | CLDN14 | 8886 |
| rs73204243 | 21 | 37824074 | a | g | -0.085 | 0.015 | 1.27E-08 | -- | 0 | 0.328 | 1 | 0.567 | 1 | CLDN14 | 8845 |
| rs78113480 | 21 | 37824320 | t | c | -0.085 | 0.015 | 1.27E-08 | -- | 0 | 0.328 | 1 | 0.567 | 1 | CLDN14 | 8599 |
| rs74439245 | 21 | 37824620 | a | g | 0.085 | 0.015 | 1.27E-08 | ++ | 0 | 0.328 | 1 | 0.567 | 1 | CLDN14 | 8299 |
| rs76422022 | 21 | 37824626 | t | g | -0.085 | 0.015 | 1.27E-08 | -- | 0 | 0.328 | 1 | 0.567 | 1 | CLDN14 | 8293 |
| rs73204244 | 21 | 37824671 | a | c | -0.085 | 0.015 | 1.27E-08 | -- | 0 | 0.328 | 1 | 0.567 | 1 | CLDN14 | 8248 |
| rs73204246 | 21 | 37824700 | c | g | 0.085 | 0.015 | 1.04E-08 | ++ | 0 | 0.367 | 1 | 0.545 | 1 | CLDN14 | 8219 |
| rs73204248 | 21 | 37824709 | a | g | -0.085 | 0.015 | 1.04E-08 | -- | 0 | 0.367 | 1 | 0.545 | 1 | CLDN14 | 8210 |
| rs4378885 | 21 | 37824911 | t | g | 0.085 | 0.015 | 1.27E-08 | ++ | 0 | 0.328 | 1 | 0.567 | 1 | CLDN14 | 8008 |
| rs4481073 | 21 | 37824955 | t | c | -0.085 | 0.015 | 1.04E-08 | -- | 0 | 0.367 | 1 | 0.545 | 1 | CLDN14 | 7964 |
| rs2835361 | 21 | 37824997 | t | c | -0.085 | 0.015 | 1.04E-08 | -- | 0 | 0.367 | 1 | 0.545 | 1 | CLDN14 | 7922 |
| rs7509937 | 21 | 37825151 | t | c | 0.085 | 0.015 | 1.04E-08 | ++ | 0 | 0.367 | 1 | 0.545 | 1 | CLDN14 | 7768 |
| rs5026478 | 21 | 37825829 | a | g | -0.089 | 0.016 | 1.06E-08 | -- | 0 | 0.264 | 1 | 0.607 | 1 | CLDN14 | 7090 |
| rs2409815 | 21 | 37825856 | t | c | 0.089 | 0.016 | 1.06E-08 | ++ | 0 | 0.264 | 1 | 0.607 | 1 | CLDN14 | 7063 |
| rs2409816 | 21 | 37825961 | t | c | -0.087 | 0.015 | 4.66E-09 | -- | 0 | 0.367 | 1 | 0.545 | 1 | CLDN14 | 6958 |
| rs60511692 | 21 | 37826389 | t | g | 0.087 | 0.015 | 5.70E-09 | ++ | 0 | 0.409 | 1 | 0.522 | 1 | CLDN14 | 6530 |
| rs2835362 | 21 | 37826547 | c | g | -0.087 | 0.015 | 5.70E-09 | -- | 0 | 0.409 | 1 | 0.522 | 1 | CLDN14 | 6372 |
| rs58602880 | 21 | 37826908 | c | g | -0.089 | 0.015 | 5.50E-09 | -- | 0 | 0.350 | 1 | 0.554 | 1 | CLDN14 | 6011 |
| rs61535886 | 21 | 37826963 | t | c | 0.089 | 0.016 | 1.06E-08 | ++ | 0 | 0.264 | 1 | 0.607 | 1 | CLDN14 | 5956 |
| rs60727080 | 21 | 37827174 | a | g | -0.089 | 0.015 | 2.52E-09 | -- | 0 | 0.409 | 1 | 0.522 | 1 | CLDN14 | 5745 |
| rs73204252 | 21 | 37827555 | a | g | -0.096 | 0.016 | 3.57E-09 | -- | 0 | 0.457 | 1 | 0.499 | 1 | CLDN14 | 5364 |
| rs4274662 | 21 | 37827694 | a | c | 0.097 | 0.016 | 1.27E-09 | ++ | 0 | 0.478 | 1 | 0.489 | 1 | CLDN14 | 5225 |
| rs2409817 | 21 | 37827708 | a | g | 0.098 | 0.016 | 1.68E-09 | ++ | 0 | 0.544 | 1 | 0.461 | 1 | CLDN14 | 5211 |
| rs2409818 | 21 | 37827846 | t | c | 0.115 | 0.020 | 6.31E-09 | ++ | 0 | 0.574 | 1 | 0.449 | 1 | CLDN14 | 5073 |
| rs128494 | 21 | 37834258 | t | c | -0.098 | 0.015 | 4.12E-11 | -- | 13.9 | 1.161 | 1 | 0.281 | 1 | CLDN14 | 0 |
| rs12626330 | 21 | 37835982 | c | g | -0.095 | 0.015 | 1.97E-10 | -- | 8.2 | 1.090 | 1 | 0.297 | 1 | CLDN14 | 0 |
| rs7277076 | 21 | 37836973 | t | c | -0.103 | 0.015 | 1.14E-11 | -- | 0 | 0.312 | 1 | 0.576 | 1 | CLDN14 | 0 |
| rs170183 | 21 | 37848334 | a | g | 0.091 | 0.015 | 8.89E-10 | ++ | 13.9 | 1.161 | 1 | 0.281 | 1 | CLDN14 | 0 |
| rs9636632 | 21 | 37855760 | a | g | 0.086 | 0.015 | 1.42E-08 | ++ | 0 | 0.039 | 1 | 0.844 | 1 | CLDN14 | 0 |
| rs13051241 | 21 | 37855854 | a | g | 0.098 | 0.016 | 3.67E-10 | ++ | 0 | 0.993 | 1 | 0.319 | 1 | CLDN14 | 0 |
| rs13051987 | 21 | 37855974 | t | c | -0.098 | 0.016 | 3.67E-10 | -- | 0 | 0.993 | 1 | 0.319 | 1 | CLDN14 | 0 |
| rs35734452 | 21 | 37856001 | t | c | 0.098 | 0.016 | 3.67E-10 | ++ | 0 | 0.993 | 1 | 0.319 | 1 | CLDN14 | 0 |
| rs219735 | 21 | 37861287 | a | g | 0.131 | 0.021 | 3.57E-10 | ++ | 5.8 | 1.062 | 1 | 0.303 | 1 | CLDN14 | 0 |
| rs219734 | 21 | 37861488 | a | g | 0.115 | 0.018 | 1.55E-10 | ++ | 0 | 0.889 | 1 | 0.346 | 1 | CLDN14 | 0 |
| rs219732 | 21 | 37864147 | t | c | 0.113 | 0.018 | 1.97E-10 | ++ | 0 | 0.673 | 1 | 0.412 | 1 | CLDN14 | 0 |
| rs9982279 | 21 | 37866276 | c | g | 0.136 | 0.022 | 5.49E-10 | ++ | 0 | 0.601 | 1 | 0.438 | 1 | CLDN14 | 0 |
| rs2071049 | 21 | 37866480 | t | c | 0.136 | 0.022 | 5.49E-10 | ++ | 0 | 0.601 | 1 | 0.438 | 1 | CLDN14 | 0 |
| rs2071052 | 21 | 37867353 | t | c | 0.140 | 0.023 | 9.91E-10 | ++ | 0 | 0.757 | 1 | 0.384 | 1 | CLDN14 | 0 |

[a]Study-specific direction of effect. [b]Heterogeneity I^2 parameter. [c]Heterogeneity Chi^2 test statistic. [d]Heterogeneity test statistics degrees of freedom. [e]Heterogeneity P value.

**Supplemental Table 2. The result of association analysis of kidney stone in the replication stage**

| SNP | Chr | Position | Effect Allele | Non Effect Allele | gene | relative loc | case | | | | control | | | | OR[b] | P |
|---|---|---|---|---|---|---|---|---|---|---|---|---|---|---|---|---|
| | | | | | | | allele1[a] | Both | allele2[a] | FREQ1[a] | allele1[a] | Both | allele2[a] | FREQ1[a] | | |
| rs6667242 | 1 | 21826566 | A | G | ALPL | -9292 | 857 | 1066 | 365 | 0.608 | 1443 | 1777 | 591 | 0.612 | 0.987 | $7.47 \times 10^{-1}$ |
| rs1260326 | 2 | 27730940 | T | C | GCKR | 0 | 770 | 1117 | 401 | 0.581 | 1223 | 1874 | 713 | 0.567 | 1.050 | $2.27 \times 10^{-1}$ |
| rs13006480 | 2 | 27987221 | C | G | MRPL33 | -7363 | 789 | 1126 | 371 | 0.591 | 1460 | 1771 | 581 | 0.615 | 0.900 | $8.98 \times 10^{-3}$ |
| rs11746443 | 5 | 176798306 | G | A | RGS14 | 0 | 1206 | 889 | 191 | 0.722 | 2134 | 1456 | 224 | 0.750 | 0.857 | $5.46 \times 10^{-4}$ |
| rs1544935 | 6 | 39124448 | T | G | KCNK5 | 32299 | 1522 | 692 | 69 | 0.818 | 2600 | 1101 | 108 | 0.827 | 0.941 | $2.39 \times 10^{-1}$ |
| rs3798519 | 6 | 50788778 | A | C | TFAP2B | 0 | 1089 | 985 | 212 | 0.692 | 1962 | 1568 | 285 | 0.720 | 0.869 | $1.28 \times 10^{-3}$ |
| rs6928986 | 6 | 131323992 | T | C | EPB41L2 | 0 | 606 | 822 | 284 | 0.594 | 1245 | 1713 | 657 | 0.581 | 1.056 | $2.23 \times 10^{-1}$ |
| rs6975977 | 7 | 30917831 | G | A | INMT-FAM188B | 0 | 1698 | 549 | 42 | 0.862 | 2895 | 846 | 70 | 0.871 | 0.927 | $1.91 \times 10^{-1}$ |
| rs7328064 | 13 | 42746118 | A | C | DGKH | 0 | 672 | 1086 | 530 | 0.531 | 1215 | 1839 | 761 | 0.560 | 0.886 | $1.95 \times 10^{-3}$ |
| rs35747824 | 16 | 20393308 | A | T | PDILT | 0 | 1370 | 797 | 119 | 0.774 | 2348 | 1284 | 182 | 0.784 | 0.943 | $2.23 \times 10^{-1}$ |
| rs7206790 | 16 | 53797908 | C | G | FTO | 0 | 1341 | 819 | 129 | 0.765 | 2312 | 1302 | 202 | 0.776 | 0.942 | $2.00 \times 10^{-1}$ |
| rs2079742 | 17 | 59465697 | T | C | BCAS3 | 0 | 596 | 1110 | 578 | 0.504 | 860 | 1870 | 1082 | 0.471 | 1.137 | $1.08 \times 10^{-3}$ |
| rs2286526 | 17 | 59472057 | C | T | LOC645722 | 0 | 960 | 1045 | 281 | 0.649 | 1741 | 1671 | 402 | 0.676 | 0.887 | $4.30 \times 10^{-3}$ |
| rs74956940 | 19 | 14571966 | C | G | PKN1 | 0 | 1244 | 866 | 176 | 0.734 | 2192 | 1395 | 227 | 0.758 | 0.869 | $1.81 \times 10^{-3}$ |
| rs13041834 | 20 | 52703284 | T | C | BCAS1 | -15980 | 1001 | 1003 | 279 | 0.658 | 1635 | 1686 | 488 | 0.651 | 1.038 | $3.68 \times 10^{-1}$ |
| rs6123359 | 20 | 52714706 | G | A | BCAS1 | -27402 | 627 | 1140 | 514 | 0.525 | 1051 | 1861 | 902 | 0.520 | 1.018 | $6.52 \times 10^{-1}$ |
| rs7277076 | 21 | 37836973 | T | C | CLDN14 | 0 | 615 | 1116 | 556 | 0.513 | 1098 | 1888 | 829 | 0.535 | 0.912 | $1.84 \times 10^{-2}$ |

[a]Allele1 and allele 2 were effect allele and non-effect allele, respectively. [b]Non-effect alleles were considered as references.

**Supplemental Table 3. Association of reported SNPs in our GWAS data set**

| SNP | Chr | Position | Effect Allele | Non Effect Allele | RSQR | gene | relative loc | screening 1 | | screening 2 | | meta_screening | | | | Risk allele in previous report |
|---|---|---|---|---|---|---|---|---|---|---|---|---|---|---|---|---|
| | | | | | | | | OR[a] | P | OR[a] | P | OR[a] | P | Q[b] | I[c] | |
| rs1256328 | 1 | 21896767 | C | T | 1.000 | ALPL | 0 | 0.960 | 1.32E-01 | 0.959 | 1.19E-01 | 0.959 | 2.97E-02 | 0.979 | 0.0 | T[1] |
| rs7627468 | 3 | 121946099 | A | G | 0.925 | CASR | 0 | 1.046 | 3.91E-02 | 1.045 | 4.16E-02 | 1.046 | 4.23E-03 | 0.974 | 0.0 | A[1] |
| rs12654812 | 5 | 176794191 | G | A | 0.999 | RGS14 | 0 | 0.878 | 1.25E-09 | 0.923 | 1.53E-04 | 0.900 | 1.54E-12 | 0.092 | 64.7 | A[1] |
| rs11746443 | 5 | 176798306 | G | A | 1.000 | RGS14 | 0 | 0.848 | 1.59E-12 | 0.899 | 4.50E-06 | 0.873 | 7.98E-17 | 0.070 | 69.6 | A[2] |
| rs1000597 | 7 | 30937178 | T | C | 0.996 | INMT-FAM188B | 5176 | 0.901 | 2.40E-05 | 0.913 | 2.58E-04 | 0.907 | 2.96E-08 | 0.692 | 0.0 | C[2] |
| rs4142110 | 13 | 42754522 | C | T | 1.000 | DGKH | 0 | 1.154 | 8.07E-12 | 1.123 | 2.21E-08 | 1.138 | 2.76E-18 | 0.363 | 0.0 | C[2] |

[a]Non-effect alleles were considered as references. [b]p-value for Cochrane's Q statistic. [c]$I^2$ heterogeneity index.

[1]Oddsson A et al Nat Commun 2015, [2]Urabe Y et al PLoS Genet 2012.

#### Supplemental Table 4. List of quantitative traits

| Category | Trait | Abbreviation |
|---|---|---|
| Metabolic | Body mass index | BMI |
| | Total cholesterol | TC |
| | High-density-lipoprotein cholesterol | HDL-C |
| | Low-density-lipoprotein cholesterol | LDL-C |
| | Triglyceride | TG |
| | Blood sugar | BS |
| | Hemoglobin A1c | HbA1c |
| Kidney-related | Blood urea nitrogen | BUN |
| | Serum creatinine | sCr |
| | Estimated glomerular filtration rate | eGFR |
| | Uric acid | UA |
| Electrolyte | Sodium | Na |
| | Potassium | K |
| | Chloride | Cl |
| | Calcium | Ca |
| | Phosphorus | P |

**Supplemental Table 5. Overview of the identified loci and their pleiotropy in metabolic traits**

| SNP | Chr | Position | Effect Allele | Non Effect Allele | meta_beta | meta_P | BMI | | | TC | | | HDL-C | | | LDL-C | | | TG | | | BS | | | HbA1c | | |
|---|---|---|---|---|---|---|---|---|---|---|---|---|---|---|---|---|---|---|---|---|---|---|---|---|---|---|---|
| | | | | | | | BETA | SE | P | BETA | SE | P | BETA | SE | P | BETA | SE | P | BETA | SE | P | BETA | SE | P | BETA | SE | P |
| rs6667242 | 1 | 21826566 | A | G | -0.088 | $2.68 \times 10^{-10}$ | 0.002 | 0.004 | 6.37E-01 | -0.001 | 0.004 | 8.33E-01 | 0.001 | 0.005 | 9.15E-01 | -0.006 | 0.005 | 2.82E-01 | -0.001 | 0.004 | 7.90E-01 | 0.004 | 0.005 | 4.21E-01 | 0.011 | 0.007 | 1.27E-01 |
| rs1260326 | 2 | 27730940 | T | C | 0.092 | $4.90 \times 10^{-11}$ | -0.005 | 0.004 | 1.37E-01 | 0.035 | 0.004 | 2.80E-19 | 0.004 | 0.005 | 4.44E-01 | 0.014 | 0.005 | 6.04E-03 | 0.089 | 0.004 | 1.69E-94 | -0.039 | 0.005 | 1.94E-16 | -0.019 | 0.007 | 6.68E-03 |
| rs13006480 | 2 | 27987221 | C | G | -0.104 | $8.25 \times 10^{-13}$ | 0.008 | 0.004 | 3.56E-02 | -0.016 | 0.004 | 1.17E-04 | -0.008 | 0.006 | 1.79E-01 | -0.007 | 0.006 | 2.08E-01 | -0.038 | 0.005 | 2.13E-16 | 0.006 | 0.005 | 2.46E-01 | 0.014 | 0.007 | 6.12E-02 |
| rs11746443 | 5 | 176798306 | G | A | -0.138 | $1.89 \times 10^{-19}$ | -0.003 | 0.004 | 5.21E-01 | -0.005 | 0.004 | 2.35E-01 | -0.004 | 0.006 | 5.54E-01 | -0.002 | 0.006 | 7.78E-01 | 0.000 | 0.005 | 9.20E-01 | -0.003 | 0.005 | 6.14E-01 | -0.014 | 0.008 | 7.16E-02 |
| rs1544935 | 6 | 39124448 | T | G | -0.111 | $5.21 \times 10^{-10}$ | -0.004 | 0.005 | 4.45E-01 | 0.010 | 0.005 | 5.02E-02 | -0.001 | 0.007 | 8.48E-01 | 0.002 | 0.007 | 7.97E-01 | 0.014 | 0.006 | 1.49E-02 | -0.013 | 0.006 | 3.17E-02 | -0.006 | 0.009 | 4.95E-01 |
| rs3798519 | 6 | 50788778 | A | C | -0.120 | $1.16 \times 10^{-15}$ | -0.031 | 0.004 | 1.67E-15 | -0.006 | 0.004 | 1.76E-01 | -0.001 | 0.006 | 9.00E-01 | 0.000 | 0.006 | 9.52E-01 | -0.004 | 0.005 | 4.43E-01 | -0.003 | 0.005 | 5.71E-01 | -0.005 | 0.008 | 5.10E-01 |
| rs6928986 | 6 | 131323992 | T | C | 0.088 | $3.63 \times 10^{-10}$ | -0.004 | 0.004 | 3.20E-01 | -0.005 | 0.004 | 2.19E-01 | 0.007 | 0.005 | 2.05E-01 | -0.003 | 0.005 | 6.35E-01 | 0.000 | 0.004 | 9.92E-01 | 0.005 | 0.005 | 2.96E-01 | 0.004 | 0.007 | 6.14E-01 |
| rs6975977 | 7 | 30917831 | G | A | -0.124 | $5.28 \times 10^{-10}$ | 0.002 | 0.005 | 7.49E-01 | -0.001 | 0.006 | 9.23E-01 | -0.003 | 0.008 | 7.43E-01 | 0.014 | 0.008 | 7.64E-02 | 0.000 | 0.006 | 9.69E-01 | -0.012 | 0.007 | 7.69E-02 | -0.018 | 0.010 | 8.26E-02 |
| rs7328064 | 13 | 42746118 | A | C | -0.137 | $3.79 \times 10^{-22}$ | 0.002 | 0.004 | 6.47E-01 | 0.000 | 0.004 | 9.39E-01 | 0.006 | 0.005 | 2.67E-01 | -0.001 | 0.005 | 8.17E-01 | 0.004 | 0.004 | 3.98E-01 | -0.007 | 0.005 | 1.48E-01 | 0.009 | 0.007 | 2.00E-01 |
| rs35747824 | 16 | 20393308 | A | T | -0.105 | $9.41 \times 10^{-11}$ | 0.004 | 0.004 | 3.68E-01 | 0.001 | 0.005 | 7.96E-01 | 0.001 | 0.006 | 8.84E-01 | 0.005 | 0.006 | 4.58E-01 | -0.001 | 0.005 | 8.72E-01 | 0.007 | 0.006 | 2.28E-01 | -0.009 | 0.008 | 2.66E-01 |
| rs7206790 | 16 | 53797908 | C | G | -0.116 | $1.03 \times 10^{-9}$ | -0.089 | 0.005 | 1.43E-65 | -0.006 | 0.006 | 2.63E-01 | 0.014 | 0.008 | 6.42E-02 | -0.005 | 0.008 | 4.80E-01 | -0.016 | 0.006 | 1.33E-02 | -0.016 | 0.007 | 1.72E-02 | -0.046 | 0.010 | 5.34E-06 |
| rs2079742 | 17 | 59465697 | T | C | 0.132 | $2.52 \times 10^{-16}$ | | | | | | | | | | | | | | | | | | | | | |
| rs2286526 | 17 | 59472057 | C | T | -0.113 | $1.13 \times 10^{-14}$ | -0.009 | 0.004 | 1.25E-02 | 0.001 | 0.004 | 7.41E-01 | 0.003 | 0.006 | 5.55E-01 | 0.002 | 0.006 | 7.51E-01 | 0.008 | 0.005 | 6.42E-02 | 0.002 | 0.005 | 6.27E-01 | -0.014 | 0.007 | 5.83E-02 |
| rs74956940 | 19 | 14571966 | C | G | -0.125 | $4.66 \times 10^{-15}$ | 0.001 | 0.004 | 8.52E-01 | -0.002 | 0.005 | 6.31E-01 | 0.007 | 0.006 | 2.34E-01 | -0.001 | 0.006 | 8.83E-01 | 0.000 | 0.005 | 9.93E-01 | 0.006 | 0.006 | 2.72E-01 | 0.002 | 0.008 | 7.77E-01 |
| rs13041834 | 20 | 52703284 | T | C | 0.080 | $3.46 \times 10^{-8}$ | -0.003 | 0.004 | 4.56E-01 | 0.009 | 0.004 | 2.49E-02 | 0.000 | 0.005 | 9.83E-01 | 0.007 | 0.005 | 1.70E-01 | 0.006 | 0.004 | 1.62E-01 | 0.009 | 0.005 | 7.35E-02 | 0.007 | 0.007 | 2.95E-01 |
| rs6123359 | 20 | 52714706 | G | A | 0.075 | $6.23 \times 10^{-8}$ | 0.000 | 0.004 | 9.30E-01 | 0.008 | 0.004 | 3.50E-02 | 0.011 | 0.005 | 3.80E-02 | 0.005 | 0.005 | 3.51E-01 | -0.001 | 0.004 | 7.77E-01 | 0.006 | 0.005 | 2.25E-01 | 0.001 | 0.007 | 8.49E-01 |
| rs7277076 | 21 | 37836973 | T | C | -0.102 | $6.37 \times 10^{-13}$ | 0.005 | 0.004 | 1.84E-01 | 0.002 | 0.004 | 6.57E-01 | 0.003 | 0.005 | 6.32E-01 | 0.000 | 0.005 | 9.71E-01 | 0.001 | 0.004 | 8.90E-01 | -0.007 | 0.005 | 1.38E-01 | 0.003 | 0.007 | 7.21E-01 |

**Supplemental Table 6. Overview of the identified loci and their pleiotropy in kidney-related traits**

| SNP | Chr | Position | Effect Allele | Non Effect Allele | meta_beta | meta_P | BUN | | | sCr | | | eGFR | | | UA | | |
|---|---|---|---|---|---|---|---|---|---|---|---|---|---|---|---|---|---|---|
| | | | | | | | BETA | SE | P | BETA | SE | P | BETA | SE | P | BETA | SE | P |
| rs6667242 | 1 | 21826566 | A | G | -0.088 | $2.68 \times 10^{-10}$ | 0.004 | 0.004 | 3.05E-01 | 0.001 | 0.004 | 7.52E-01 | -0.001 | 0.004 | 8.62E-01 | -0.003 | 0.004 | 4.63E-01 |
| rs1260326 | 2 | 27730940 | T | C | 0.092 | $4.90 \times 10^{-11}$ | 0.012 | 0.004 | 1.01E-03 | -0.034 | 0.004 | 9.25E-21 | 0.032 | 0.004 | 3.59E-17 | 0.035 | 0.004 | 2.65E-16 |
| rs13006480 | 2 | 27987221 | C | G | -0.104 | $8.25 \times 10^{-13}$ | -0.020 | 0.004 | 1.60E-07 | 0.017 | 0.004 | 1.26E-05 | -0.014 | 0.004 | 4.67E-04 | -0.003 | 0.005 | 4.96E-01 |
| rs11746443 | 5 | 176798306 | G | A | -0.138 | $1.89 \times 10^{-19}$ | -0.015 | 0.004 | 1.69E-04 | -0.034 | 0.004 | 2.53E-16 | 0.034 | 0.004 | 1.22E-15 | 0.003 | 0.005 | 5.57E-01 |
| rs1544935 | 6 | 39124448 | T | G | -0.111 | $5.21 \times 10^{-10}$ | -0.016 | 0.005 | 9.29E-04 | 0.015 | 0.005 | 1.90E-03 | -0.012 | 0.005 | 1.24E-02 | 0.019 | 0.006 | 4.97E-04 |
| rs3798519 | 6 | 50788778 | A | C | -0.120 | $1.16 \times 10^{-15}$ | -0.031 | 0.004 | 3.75E-15 | 0.007 | 0.004 | 9.24E-02 | -0.006 | 0.004 | 1.35E-01 | 0.005 | 0.005 | 2.73E-01 |
| rs6928986 | 6 | 131323992 | T | C | 0.088 | $3.63 \times 10^{-10}$ | 0.000 | 0.004 | 8.93E-01 | -0.010 | 0.004 | 5.57E-03 | 0.012 | 0.004 | 1.75E-03 | -0.007 | 0.004 | 8.71E-02 |
| rs6975977 | 7 | 30917831 | G | A | -0.124 | $5.28 \times 10^{-10}$ | 0.002 | 0.005 | 6.43E-01 | -0.013 | 0.005 | 1.76E-02 | 0.016 | 0.006 | 5.76E-03 | -0.011 | 0.006 | 9.18E-02 |
| rs7328064 | 13 | 42746118 | A | C | -0.137 | $3.79 \times 10^{-22}$ | 0.020 | 0.004 | 7.95E-08 | 0.024 | 0.004 | 2.00E-10 | -0.025 | 0.004 | 2.80E-10 | 0.010 | 0.004 | 1.80E-02 |
| rs35747824 | 16 | 20393308 | A | T | -0.105 | $9.41 \times 10^{-11}$ | 0.046 | 0.004 | 1.79E-27 | 0.068 | 0.004 | 2.82E-53 | -0.073 | 0.005 | 8.80E-59 | 0.025 | 0.005 | 6.50E-07 |
| rs7206790 | 16 | 53797908 | C | G | -0.116 | $1.03 \times 10^{-9}$ | -0.038 | 0.005 | 3.19E-13 | -0.014 | 0.005 | 8.94E-03 | 0.012 | 0.005 | 3.33E-02 | 0.003 | 0.006 | 6.60E-01 |
| rs2079742 | 17 | 59465697 | T | C | 0.132 | $2.52 \times 10^{-16}$ | | | | | | | | | | | | |
| rs2286526 | 17 | 59472057 | C | T | -0.113 | $1.13 \times 10^{-14}$ | 0.003 | 0.004 | 4.31E-01 | 0.034 | 0.004 | 1.54E-18 | -0.036 | 0.004 | 8.94E-20 | -0.030 | 0.004 | 5.07E-11 |
| rs74956940 | 19 | 14571966 | C | G | -0.125 | $4.66 \times 10^{-15}$ | 0.003 | 0.004 | 4.46E-01 | 0.017 | 0.004 | 6.28E-05 | -0.016 | 0.004 | 2.46E-04 | 0.018 | 0.005 | 3.87E-04 |
| rs13041834 | 20 | 52703284 | T | C | 0.080 | $3.46 \times 10^{-8}$ | 0.010 | 0.004 | 7.87E-03 | 0.008 | 0.004 | 4.81E-02 | -0.008 | 0.004 | 4.00E-02 | 0.010 | 0.004 | 2.76E-02 |
| rs6123359 | 20 | 52714706 | G | A | 0.075 | $6.23 \times 10^{-8}$ | 0.011 | 0.004 | 2.73E-03 | 0.014 | 0.004 | 1.81E-04 | -0.015 | 0.004 | 9.29E-05 | 0.009 | 0.004 | 2.92E-02 |
| rs7277076 | 21 | 37836973 | T | C | -0.102 | $6.37 \times 10^{-13}$ | 0.015 | 0.004 | 3.40E-05 | 0.020 | 0.004 | 1.12E-07 | -0.023 | 0.004 | 3.44E-09 | 0.006 | 0.004 | 1.78E-01 |

**Supplemental Table 7. Overview of the identified loci and their pleiotropy in electrolyte traits**

| SNP | Chr | Position | Effect Allele | Non Effect Allele | meta_beta | meta_P | Na | | | K | | | Cl | | | Ca | | | P | | |
|---|---|---|---|---|---|---|---|---|---|---|---|---|---|---|---|---|---|---|---|---|---|
| | | | | | | | BETA | SE | P | BETA | SE | P | BETA | SE | P | BETA | SE | P | BETA | SE | P |
| rs6667242 | 1 | 21826566 | A | G | -0.088 | $2.68 \times 10^{-10}$ | -0.001 | 0.004 | 7.84E-01 | -0.004 | 0.004 | 2.74E-01 | -0.012 | 0.004 | 2.69E-03 | 0.011 | 0.005 | 4.01E-02 | 0.056 | 0.007 | 1.76E-15 |
| rs1260326 | 2 | 27730940 | T | C | 0.092 | $4.90 \times 10^{-11}$ | -0.012 | 0.004 | 3.25E-03 | 0.006 | 0.004 | 1.44E-01 | -0.024 | 0.004 | 1.44E-09 | 0.022 | 0.005 | 4.55E-05 | 0.018 | 0.007 | 8.51E-03 |
| rs13006480 | 2 | 27987221 | C | G | -0.104 | $8.25 \times 10^{-13}$ | 0.007 | 0.004 | 8.53E-02 | 0.002 | 0.004 | 5.90E-01 | 0.011 | 0.004 | 8.72E-03 | -0.021 | 0.006 | 2.71E-04 | -0.014 | 0.007 | 5.82E-02 |
| rs11746443 | 5 | 176798306 | G | A | -0.138 | $1.89 \times 10^{-19}$ | 0.019 | 0.005 | 1.95E-05 | -0.002 | 0.004 | 7.05E-01 | 0.020 | 0.005 | 8.77E-06 | -0.022 | 0.006 | 3.98E-04 | 0.075 | 0.008 | 2.58E-21 |
| rs1544935 | 6 | 39124448 | T | G | -0.111 | $5.21 \times 10^{-10}$ | -0.005 | 0.005 | 3.10E-01 | 0.005 | 0.005 | 3.72E-01 | -0.002 | 0.005 | 7.31E-01 | -0.015 | 0.007 | 3.24E-02 | 0.009 | 0.009 | 3.27E-01 |
| rs3798519 | 6 | 50788778 | A | C | -0.120 | $1.16 \times 10^{-15}$ | -0.004 | 0.004 | 4.13E-01 | -0.003 | 0.004 | 5.51E-01 | -0.003 | 0.004 | 4.90E-01 | 0.008 | 0.006 | 1.56E-01 | -0.005 | 0.008 | 4.99E-01 |
| rs6928986 | 6 | 131323992 | T | C | 0.088 | $3.63 \times 10^{-10}$ | 0.008 | 0.004 | 5.02E-02 | -0.001 | 0.004 | 8.45E-01 | 0.007 | 0.004 | 9.42E-02 | 0.009 | 0.005 | 8.82E-02 | -0.005 | 0.007 | 4.92E-01 |
| rs6975977 | 7 | 30917831 | G | A | -0.124 | $5.28 \times 10^{-10}$ | 0.008 | 0.006 | 1.86E-01 | -0.005 | 0.006 | 3.92E-01 | -0.004 | 0.006 | 4.59E-01 | -0.002 | 0.008 | 8.03E-01 | 0.005 | 0.010 | 6.39E-01 |
| rs7328064 | 13 | 42746118 | A | C | -0.137 | $3.79 \times 10^{-22}$ | -0.009 | 0.004 | 2.38E-02 | 0.001 | 0.004 | 8.75E-01 | -0.005 | 0.004 | 2.61E-01 | 0.012 | 0.005 | 2.51E-02 | 0.017 | 0.007 | 1.60E-02 |
| rs35747824 | 16 | 20393308 | A | T | -0.105 | $9.41 \times 10^{-11}$ | -0.003 | 0.005 | 5.37E-01 | 0.004 | 0.005 | 3.58E-01 | 0.001 | 0.005 | 8.48E-01 | 0.010 | 0.006 | 1.31E-01 | -0.001 | 0.008 | 9.17E-01 |
| rs7206790 | 16 | 53797908 | C | G | -0.116 | $1.03 \times 10^{-9}$ | -0.010 | 0.006 | 1.02E-01 | -0.003 | 0.006 | 6.54E-01 | -0.006 | 0.006 | 2.68E-01 | -0.002 | 0.008 | 7.87E-01 | 0.012 | 0.010 | 2.42E-01 |
| rs2079742 | 17 | 59465697 | T | C | 0.132 | $2.52 \times 10^{-16}$ | | | | | | | | | | | | | | | |
| rs2286526 | 17 | 59472057 | C | T | -0.113 | $1.13 \times 10^{-14}$ | -0.004 | 0.004 | 3.11E-01 | -0.030 | 0.004 | 2.84E-13 | -0.037 | 0.004 | 2.68E-18 | 0.031 | 0.006 | 2.85E-08 | -0.005 | 0.007 | 5.23E-01 |
| rs74956940 | 19 | 14571966 | C | G | -0.125 | $4.66 \times 10^{-15}$ | 0.002 | 0.005 | 5.97E-01 | 0.009 | 0.005 | 6.32E-02 | 0.002 | 0.005 | 6.61E-01 | 0.012 | 0.006 | 5.46E-02 | 0.016 | 0.008 | 4.81E-02 |
| rs13041834 | 20 | 52703284 | T | C | 0.080 | $3.46 \times 10^{-8}$ | -0.004 | 0.004 | 3.81E-01 | 0.006 | 0.004 | 1.33E-01 | -0.022 | 0.004 | 6.57E-08 | 0.023 | 0.006 | 2.15E-05 | 0.014 | 0.007 | 5.56E-02 |
| rs6123359 | 20 | 52714706 | G | A | 0.075 | $6.23 \times 10^{-8}$ | 0.000 | 0.004 | 9.30E-01 | 0.002 | 0.004 | 5.42E-01 | -0.018 | 0.004 | 1.20E-05 | 0.033 | 0.005 | 9.52E-10 | 0.014 | 0.007 | 4.09E-02 |
| rs7277076 | 21 | 37836973 | T | C | -0.102 | $6.37 \times 10^{-13}$ | -0.002 | 0.004 | 6.72E-01 | 0.001 | 0.004 | 7.50E-01 | -0.002 | 0.004 | 6.81E-01 | 0.014 | 0.005 | 9.56E-03 | 0.020 | 0.007 | 4.47E-03 |

#### Supplemental Table 8. wGRS analysis of 17 associated SNPs.

replicatoin stage

| quantiles | wGRS | case | control | case _freq | control _freq | OR (95% CI) | P |
|---|---|---|---|---|---|---|---|
| 1 | 0.190-1.100 | 301 | 767 | 0.282 | 0.718 | | |
| 2 | 1.100-1.285 | 301 | 766 | 0.282 | 0.718 | 1.001 (0.825-1.215) | 1.00E+00 |
| 3 | 1.286-1.452 | 331 | 737 | 0.310 | 0.690 | 1.144 (0.946-1.385) | 1.69E-01 |
| 4 | 1.452-1.654 | 354 | 713 | 0.332 | 0.668 | 1.265 (1.048-1.528) | 1.29E-02 |
| 5 | 1.655-2.688 | 429 | 639 | 0.402 | 0.598 | 1.710 (1.422-2.059) | 6.46E-09 |
| | | 1716 | 3622 | | | | |

screening stage

| quantiles | wGRS | case | control | case _freq | control _freq | OR (95% CI) | P |
|---|---|---|---|---|---|---|---|
| 1 | 0.112-1.091 | 1509 | 38245 | 0.038 | 0.962 | | |
| 2 | 1.091-1.271 | 1868 | 37886 | 0.047 | 0.953 | 1.250 (1.165-1.340) | 2.95E-10 |
| 3 | 1.271-1.430 | 2170 | 37583 | 0.055 | 0.945 | 1.463 (1.368-1.566) | < 2.20E-16 |
| 4 | 1.430-1.616 | 2501 | 37253 | 0.063 | 0.937 | 1.701 (1.593-1.818) | < 2.20E-16 |
| 5 | 1.616-2.850 | 3082 | 36672 | 0.078 | 0.922 | 2.130 (1.999-2.271) | < 2.20E-16 |
| | | 11130 | 187639 | | | | |

### Supplemental Figure 1

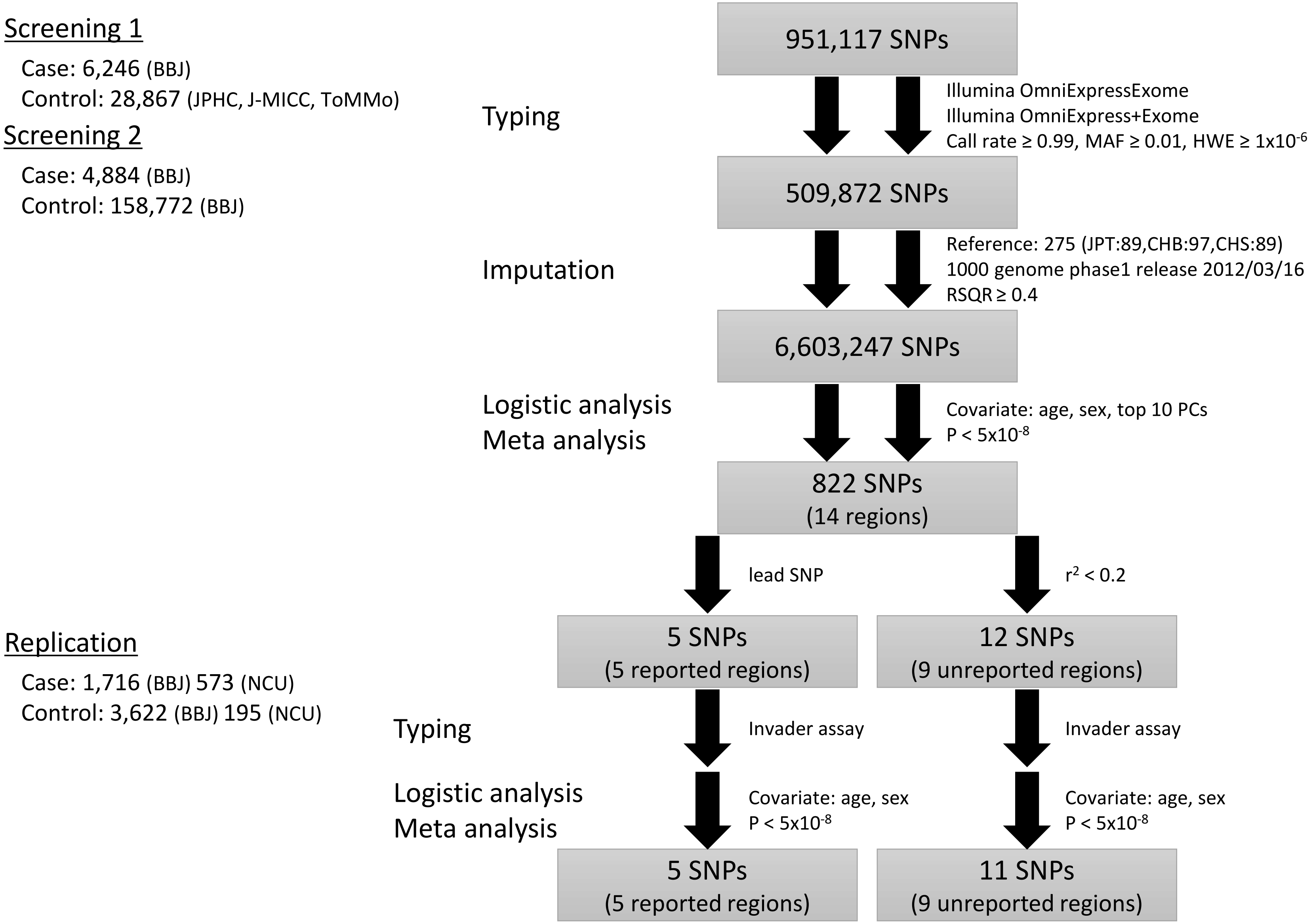


**Supplemental Figure 1** Scheme for study design and screening results.

### Supplemental Figure 2

$\lambda$=1.164
$\lambda_{1000}$=1.008

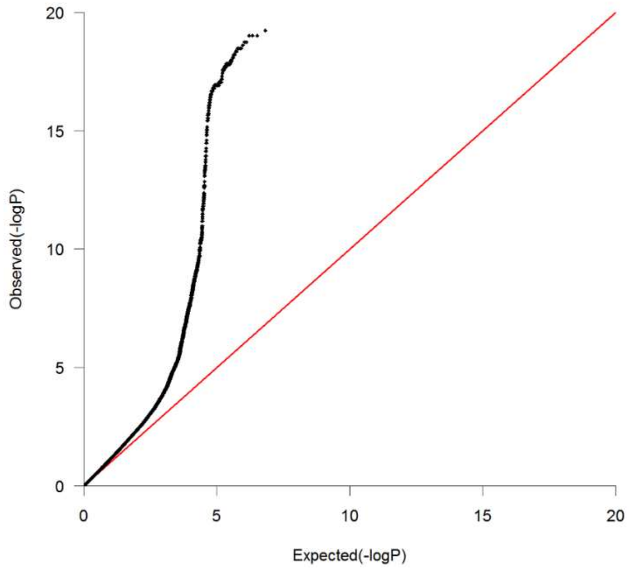


**Supplemental Figure 2** Quantile quantile plot for imputed GWAS of nephrolithiasis in the Japanese population.

### Supplemental Figure 3

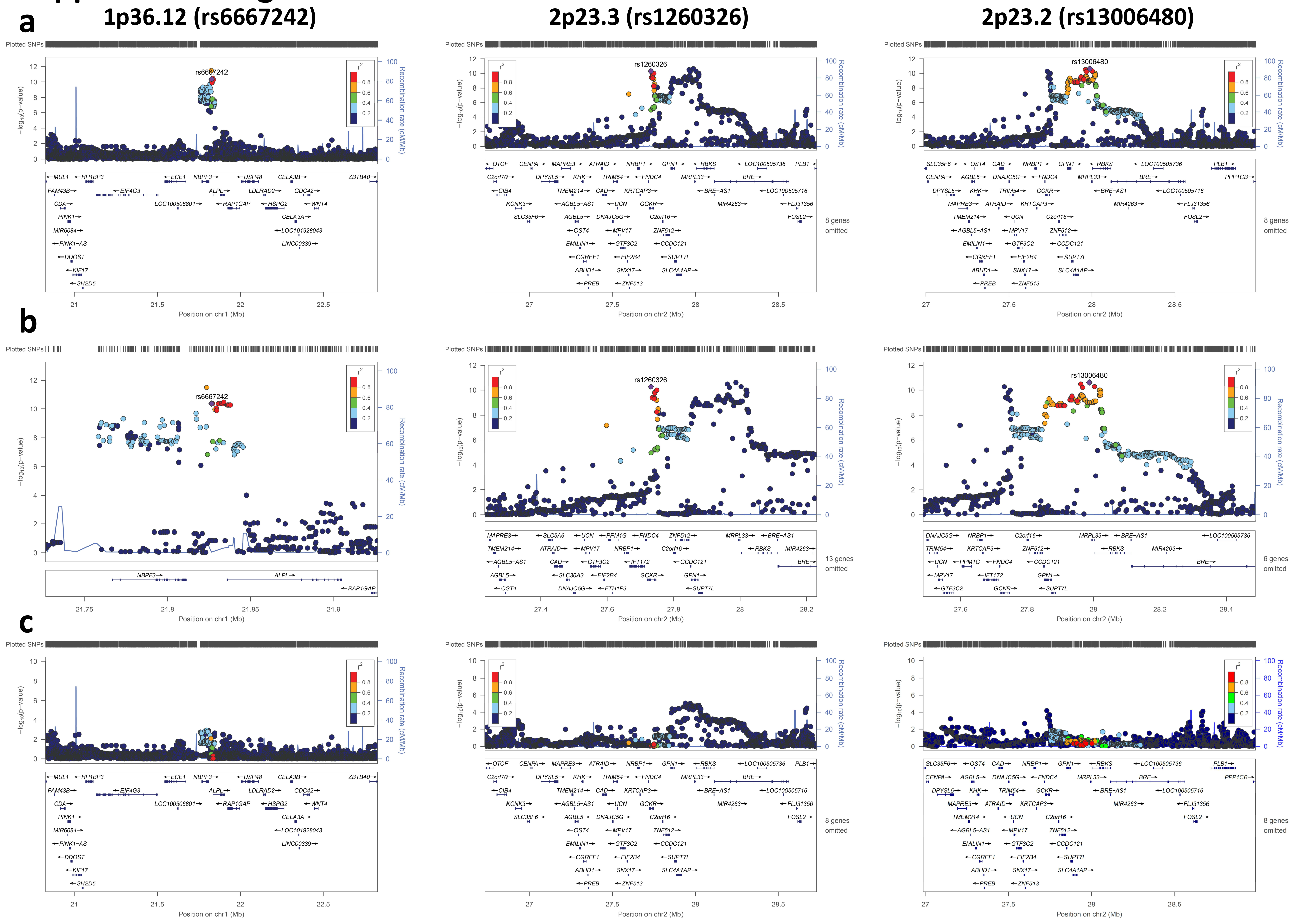

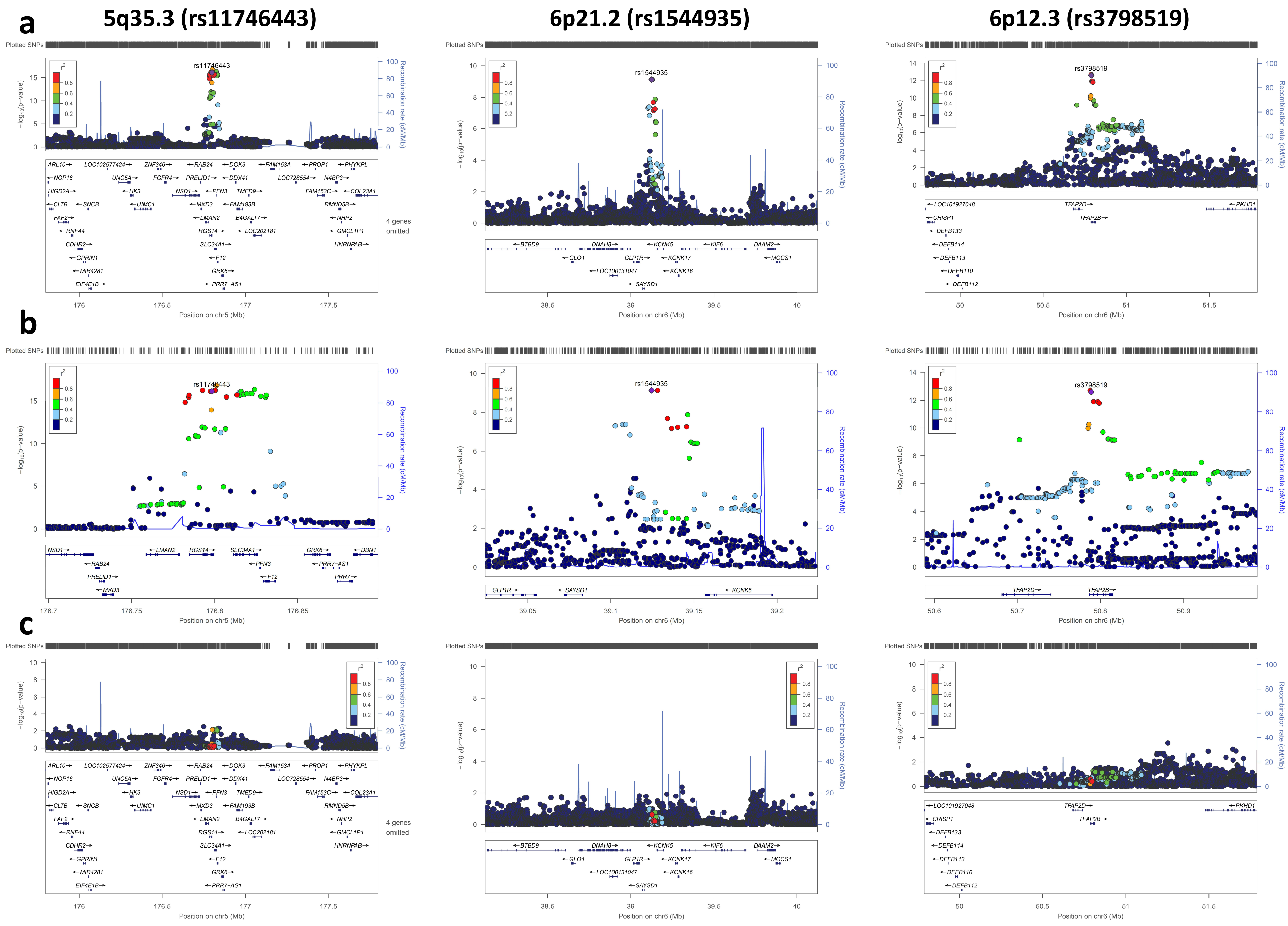

a
5q35.3 (rs11746443)
6p21.2 (rs1544935)
6p12.3 (rs3798519)
b
c
Plotted SNPs
rs11746443
rs1544935
rs3798519
−log10(p−value)
Recombination rate (cM/Mb)
Position on chr5 (Mb)
Position on chr6 (Mb)
4 genes omitted

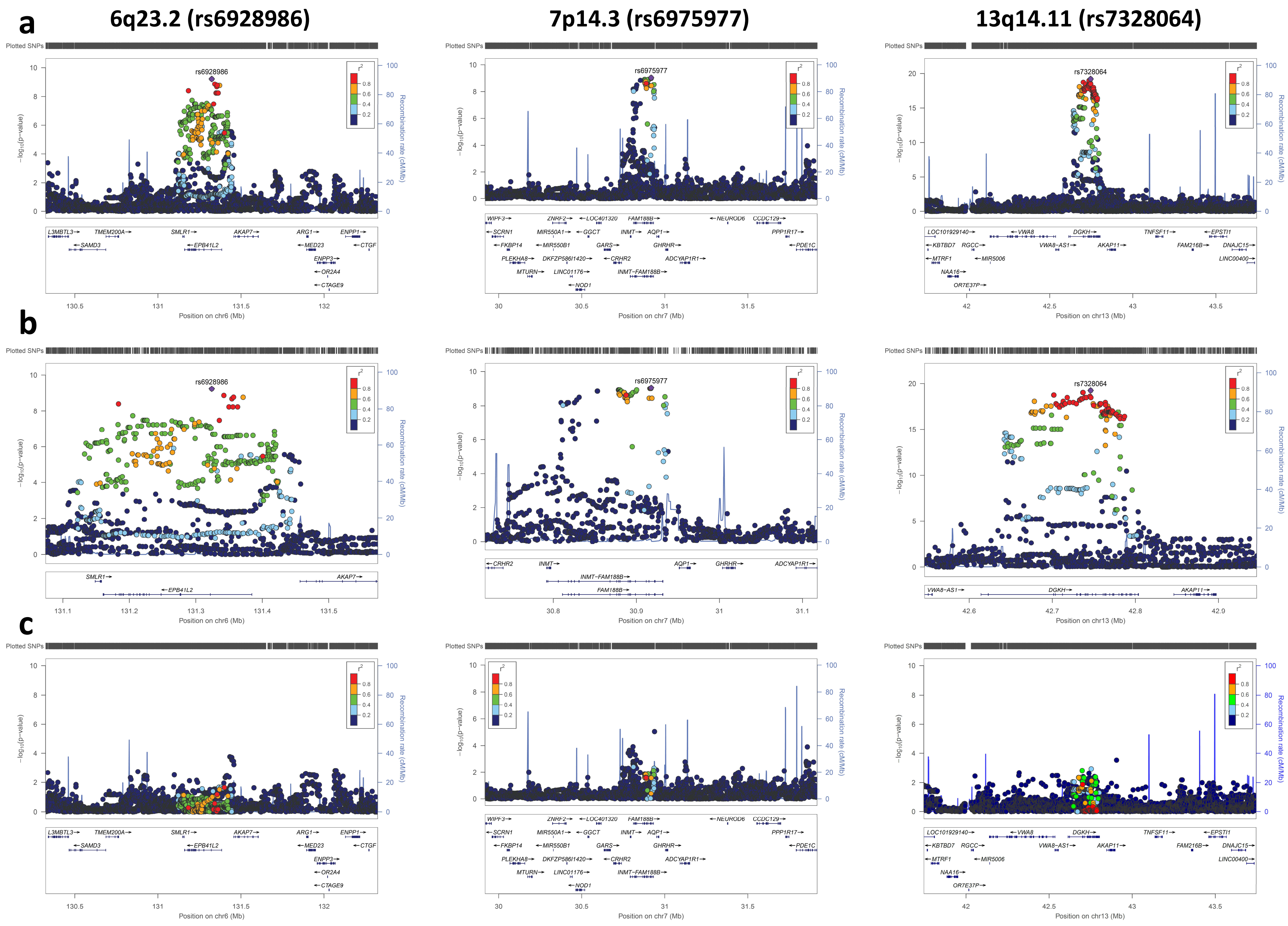
a
6q23.2 (rs6928986)
7p14.3 (rs6975977)
13q14.11 (rs7328064)
b
c
Plotted SNPs
rs6928986
rs6975977
rs7328064
−log10(p−value)
Recombination rate (cM/Mb)
r2
Position on chr6 (Mb)
Position on chr7 (Mb)
Position on chr13 (Mb)
L3MBTL3
TMEM200A
SMLR1
AKAP7
ARG1
ENPP1
SAMD3
EPB41L2
MED23
CTGF
ENPP3
OR2A4
CTAGE9
WIPF3
ZNRF2
LOC401320
FAM188B
NEUROD6
CCDC129
SCRN1
MIR550A1
GGCT
INMT
AQP1
PPP1R17
FKBP14
MIR550B1
GARS
GHRHR
PDE1C
PLEKHA8
DKFZP586I1420
CRHR2
ADCYAP1R1
MTURN
LINC01176
INMT−FAM188B
NOD1
LOC101929140
VWA8
DGKH
TNFSF11
EPSTI1
KBTBD7
RGCC
VWA8−AS1
AKAP11
FAM216B
DNAJC15
MTRF1
MIR5006
LINC00400
NAA16
OR7E37P

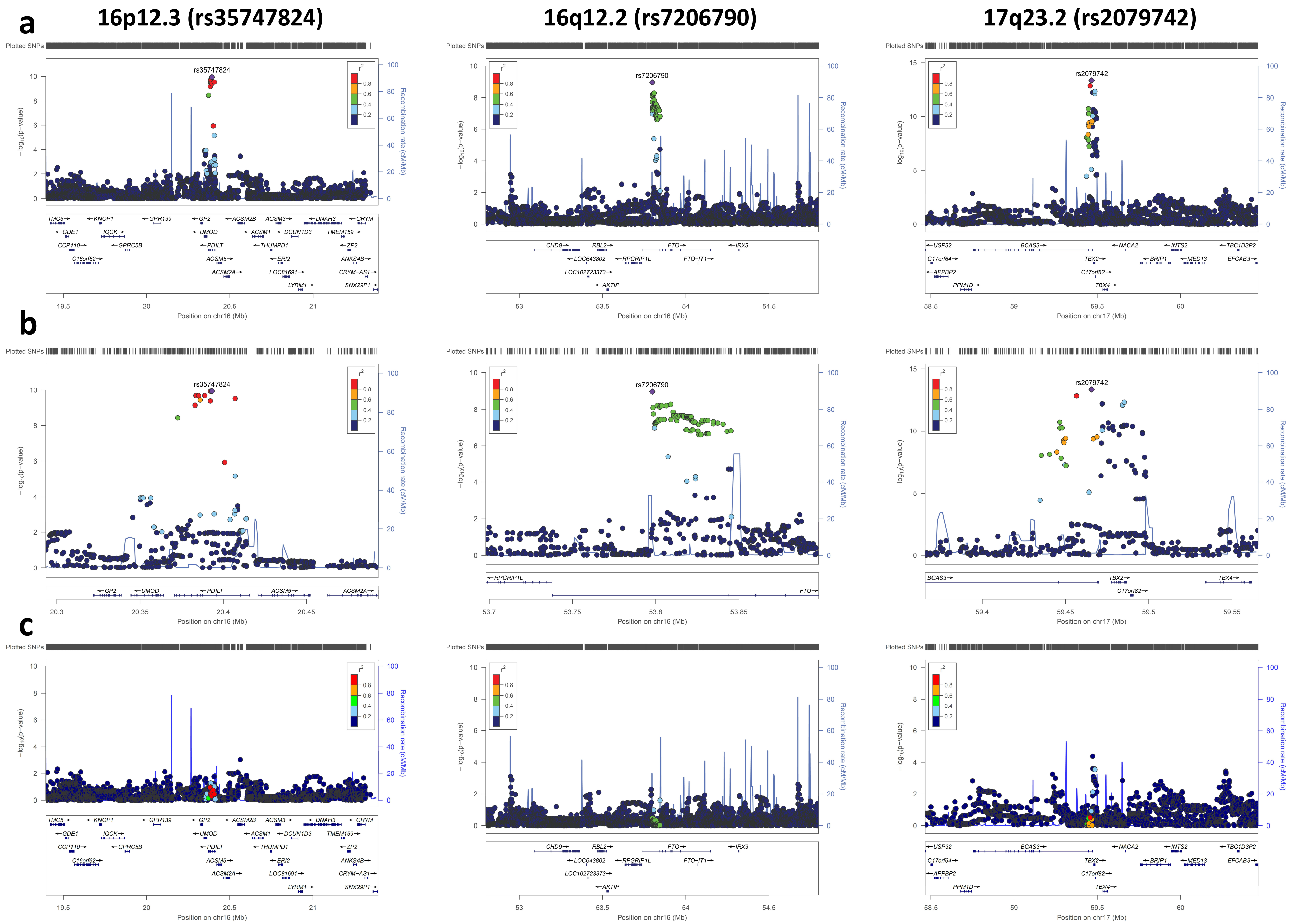
a
b
c
16p12.3 (rs35747824)
16q12.2 (rs7206790)
17q23.2 (rs2079742)
Plotted SNPs
rs35747824
rs7206790
rs2079742
r²
0.8
0.6
0.4
0.2
−log₁₀(p−value)
Recombination rate (cM/Mb)
Position on chr16 (Mb)
Position on chr17 (Mb)
TMC5 KNOP1 GPR139 GP2 ACSM2B ACSM3 DNAH3 CRYM
GDE1 IQCK UMOD ACSM1 DCUN1D3 TMEM159
CCP110 GPRC5B PDILT THUMPD1 ZP2
C16orf62 ACSM5 ERI2 ANKS4B
ACSM2A LOC81691 CRYM−AS1
LYRM1 SNX29P1
CHD9 RBL2 FTO IRX3
LOC643802 RPGRIP1L FTO−IT1
LOC102723373
AKTIP
USP32 BCAS3 NACA2 INTS2 TBC1D3P2
C17orf64 TBX2 BRIP1 MED13 EFCAB3
APPBP2 C17orf82
PPM1D TBX4
RPGRIP1L
FTO
BCAS3 TBX2 TBX4
C17orf82

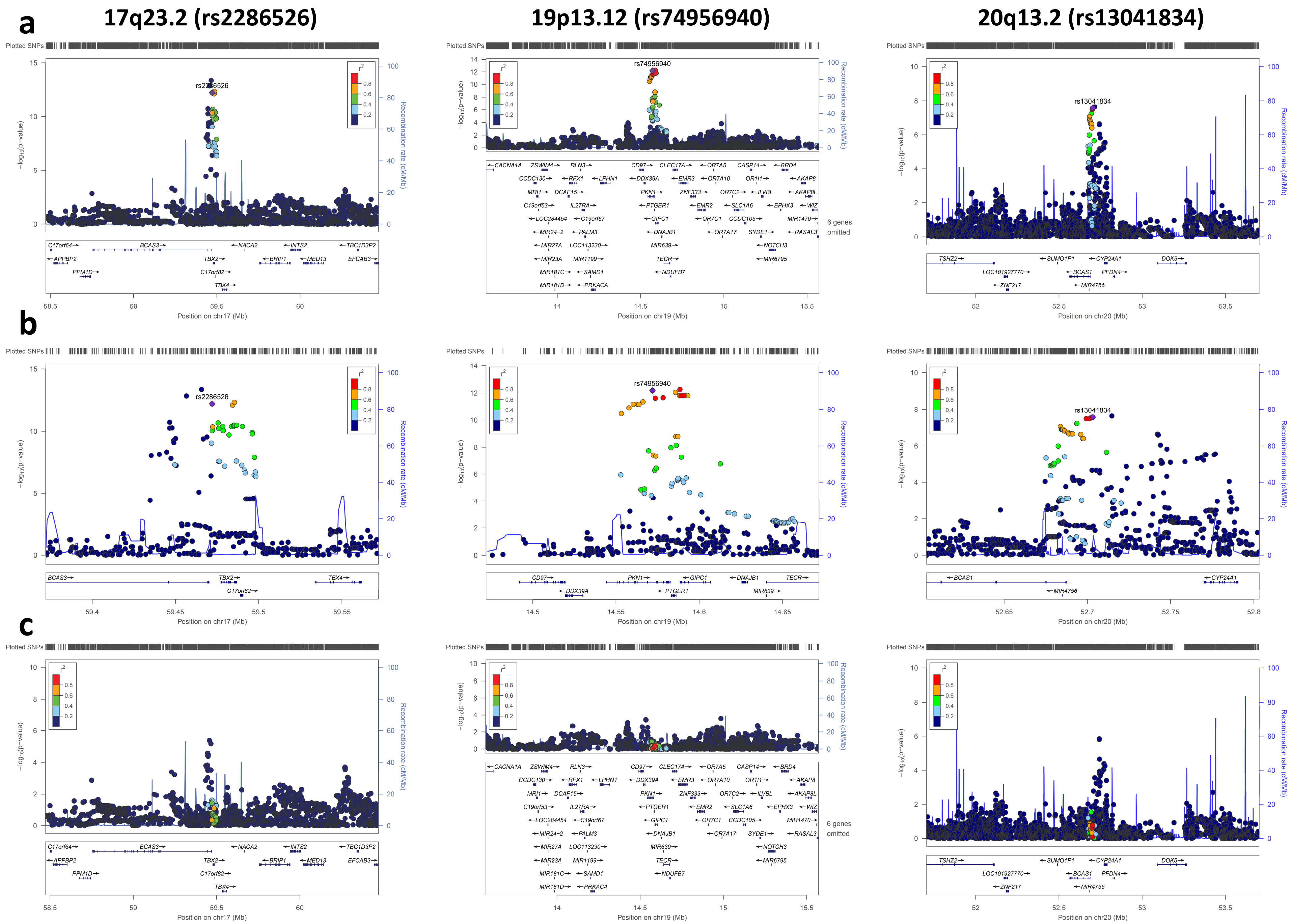
a
b
c
17q23.2 (rs2286526)
19p13.12 (rs74956940)
20q13.2 (rs13041834)
Plotted SNPs
−log10(p−value)
Recombination rate (cM/Mb)
r2
rs2286526
rs74956940
rs13041834
Position on chr17 (Mb)
Position on chr19 (Mb)
Position on chr20 (Mb)
6 genes omitted
C17orf64
BCAS3
NACA2
INTS2
TBC1D3P2
APPBP2
TBX2
BRIP1
MED13
EFCAB3
PPM1D
C17orf82
TBX4
CACNA1A
ZSWIM4
RLN3
CD97
CLEC17A
OR7A5
CASP14
BRD4
CCDC130
RFX1
LPHN1
DDX39A
EMR3
OR7A10
OR1I1
AKAP8
MRI1
DCAF15
PKN1
ZNF333
OR7C2
ILVBL
AKAP8L
C19orf53
IL27RA
PTGER1
EMR2
SLC1A6
EPHX3
WIZ
LOC284454
C19orf67
GIPC1
OR7C1
CCDC105
MIR1470
MIR24−2
PALM3
DNAJB1
OR7A17
SYDE1
RASAL3
MIR27A
LOC113230
MIR639
NOTCH3
MIR23A
MIR1199
TECR
MIR6795
MIR181C
SAMD1
NDUFB7
MIR181D
PRKACA
TSHZ2
SUMO1P1
CYP24A1
DOK5
LOC101927770
BCAS1
PFDN4
ZNF217
MIR4756

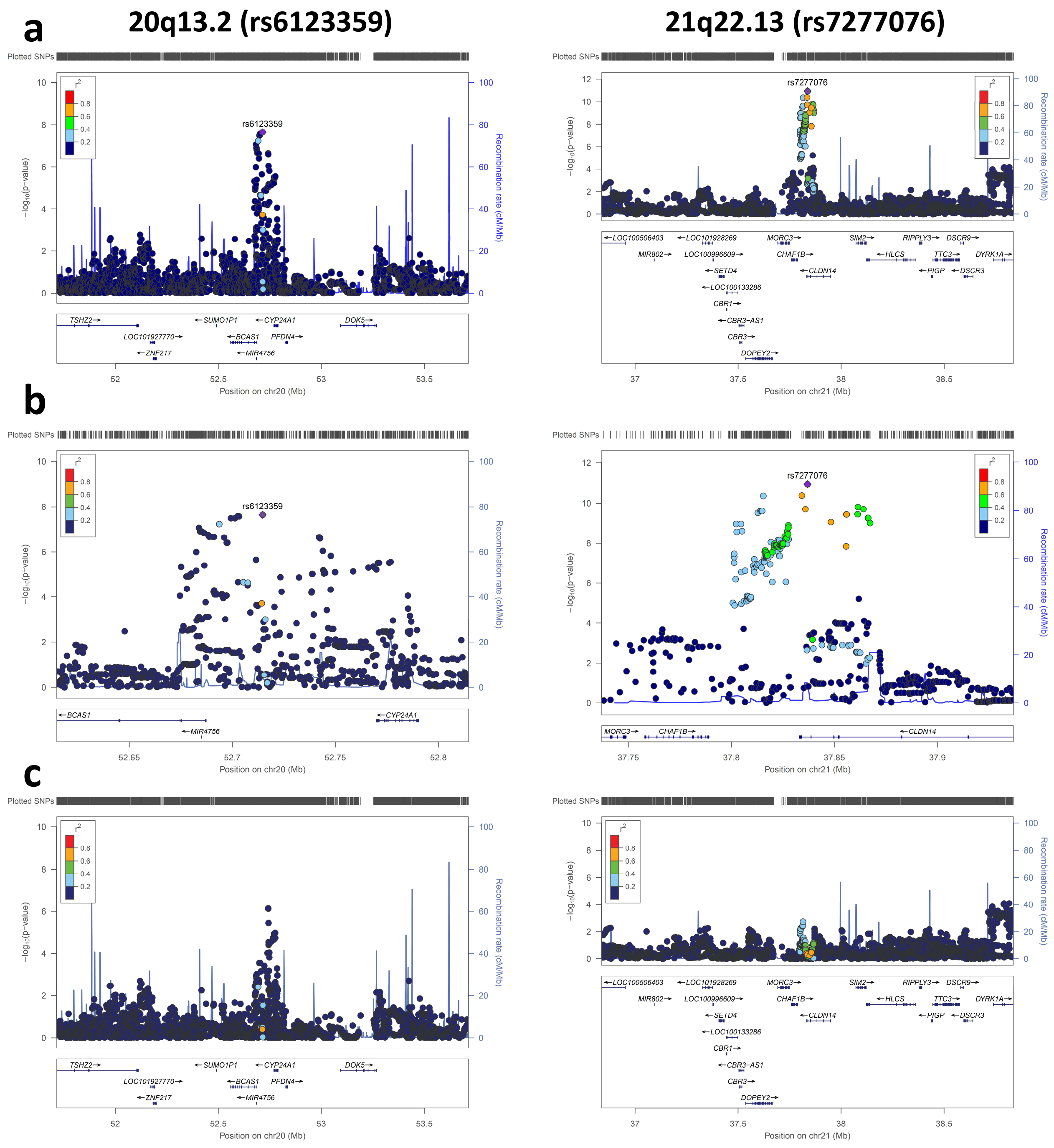


**Supplemental Figure 3 Regional plots of 17 loci for nephrolithiasis before and after condition.**
(**a**) The $-\log_{10} P$ values from the meta-analysis of screening 1 and screening 2 in 17 loci are shown. (**b**) Enlarged regional plots. (**c**) Regional plots after condition on genotyped SNP within each region. Estimated recombination rates (from 1000 Genomes) are plotted in blue. The SNPs are color-coded to reflect their correlation with genotyped SNPs. Pairwise r2 values are from 1000 Genomes East Asian data (March 2012 release). Genes, position of exons and direction of transcription from UCSC genome browser (genome.ucsc.edu) are noted. Plots were generated using LocusZoom (http://csg.sph.umich.edu/locuszoom).

### Supplemental Figure 4

* $P<5x10^{-8}$
* $P<0.05$

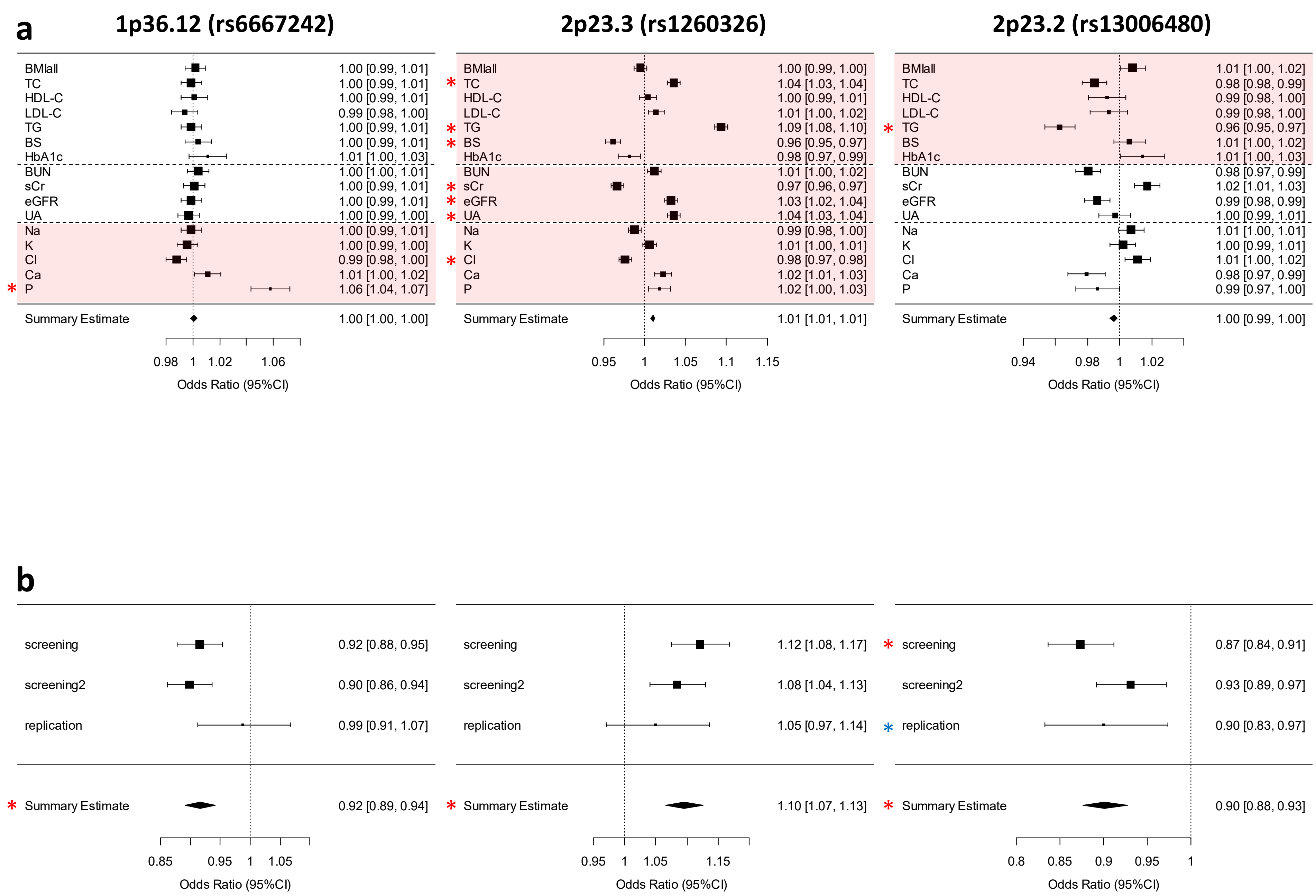

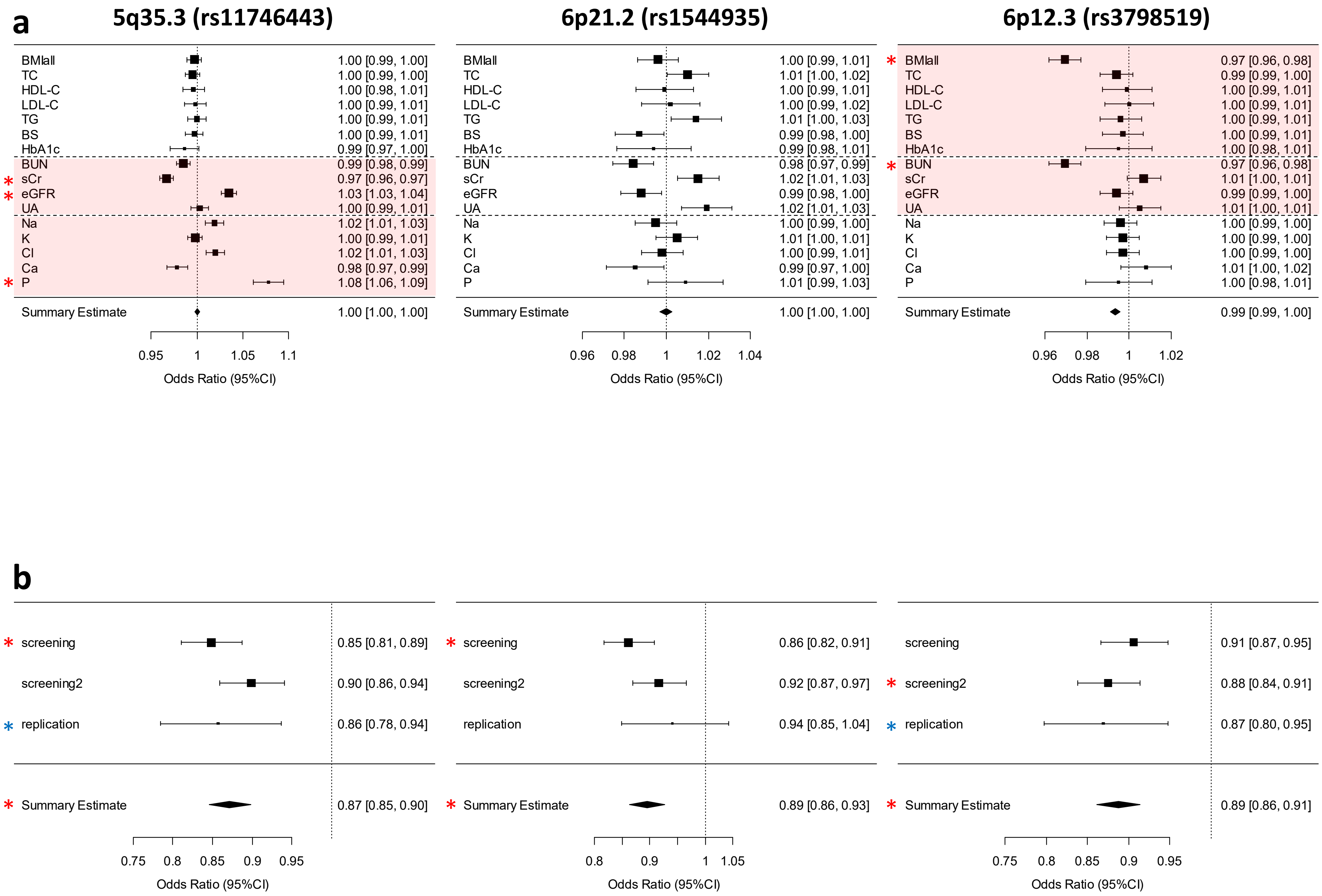
a
5q35.3 (rs11746443)
6p21.2 (rs1544935)
6p12.3 (rs3798519)
BMIall
TC
HDL-C
LDL-C
TG
BS
HbA1c
BUN
sCr
eGFR
UA
Na
K
Cl
Ca
P
Summary Estimate
1.00 [0.99, 1.00]
1.00 [0.99, 1.00]
1.00 [0.98, 1.01]
1.00 [0.99, 1.01]
1.00 [0.99, 1.01]
1.00 [0.99, 1.01]
0.99 [0.97, 1.00]
0.99 [0.98, 0.99]
0.97 [0.96, 0.97]
1.03 [1.03, 1.04]
1.00 [0.99, 1.01]
1.02 [1.01, 1.03]
1.00 [0.99, 1.01]
1.02 [1.01, 1.03]
0.98 [0.97, 0.99]
1.08 [1.06, 1.09]
1.00 [1.00, 1.00]
1.00 [0.99, 1.01]
1.01 [1.00, 1.02]
1.00 [0.99, 1.01]
1.00 [0.99, 1.02]
1.01 [1.00, 1.03]
0.99 [0.98, 1.00]
0.99 [0.98, 1.01]
0.98 [0.97, 0.99]
1.02 [1.01, 1.03]
0.99 [0.98, 1.00]
1.02 [1.01, 1.03]
1.00 [0.99, 1.00]
1.01 [1.00, 1.01]
1.00 [0.99, 1.01]
0.99 [0.97, 1.00]
1.01 [0.99, 1.03]
1.00 [1.00, 1.00]
0.97 [0.96, 0.98]
0.99 [0.99, 1.00]
1.00 [0.99, 1.01]
1.00 [0.99, 1.01]
1.00 [0.99, 1.01]
1.00 [0.99, 1.01]
1.00 [0.98, 1.01]
0.97 [0.96, 0.98]
1.01 [1.00, 1.01]
0.99 [0.99, 1.00]
1.01 [1.00, 1.01]
1.00 [0.99, 1.00]
1.00 [0.99, 1.00]
1.00 [0.99, 1.00]
1.01 [1.00, 1.02]
1.00 [0.98, 1.01]
0.99 [0.99, 1.00]
0.95 1 1.05 1.1
0.96 0.98 1 1.02 1.04
0.96 0.98 1 1.02
Odds Ratio (95%CI)
b
screening
screening2
replication
Summary Estimate
0.85 [0.81, 0.89]
0.90 [0.86, 0.94]
0.86 [0.78, 0.94]
0.87 [0.85, 0.90]
0.86 [0.82, 0.91]
0.92 [0.87, 0.97]
0.94 [0.85, 1.04]
0.89 [0.86, 0.93]
0.91 [0.87, 0.95]
0.88 [0.84, 0.91]
0.87 [0.80, 0.95]
0.89 [0.86, 0.91]
0.75 0.8 0.85 0.9 0.95
0.8 0.9 1 1.05
Odds Ratio (95%CI)

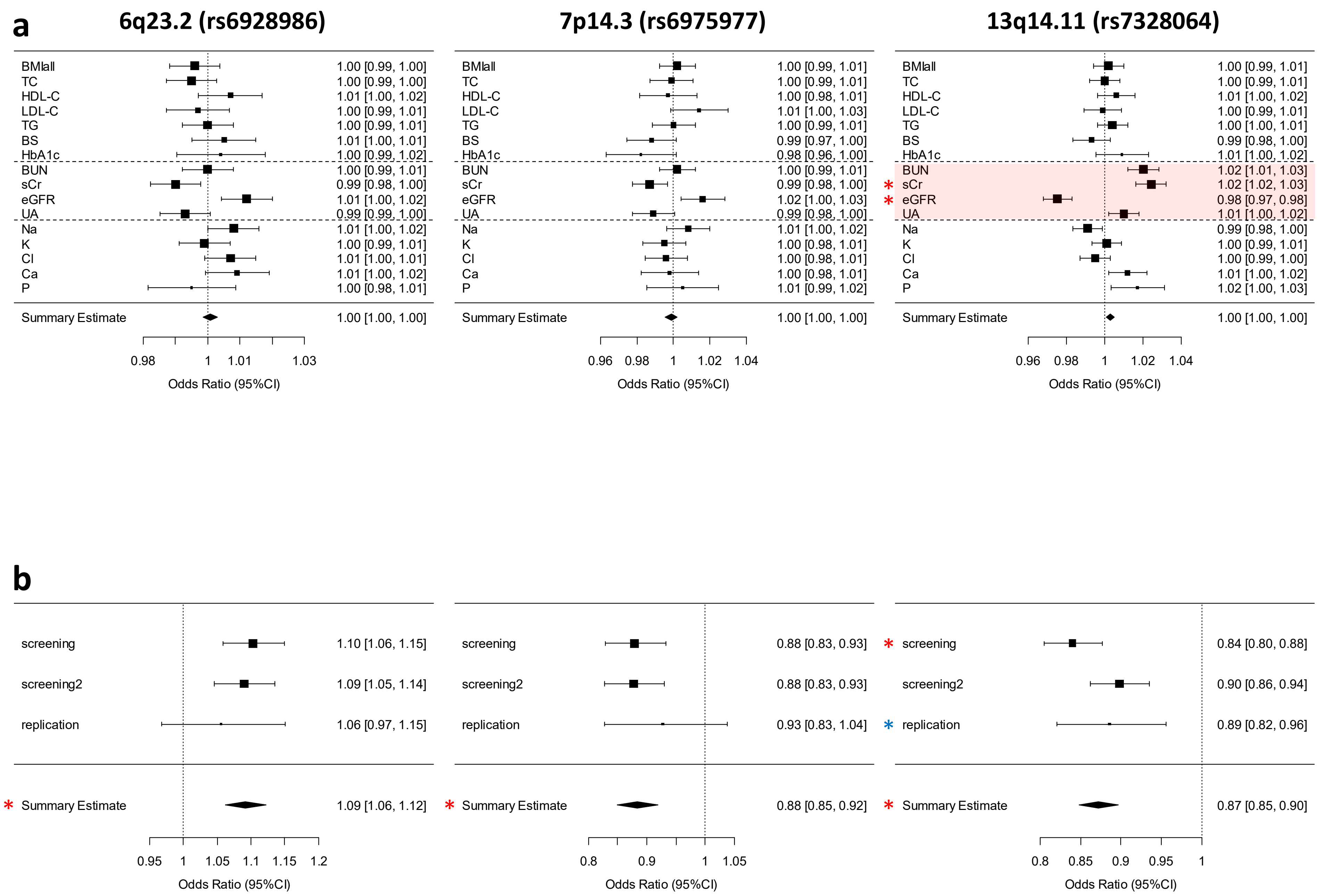
a
6q23.2 (rs6928986)
7p14.3 (rs6975977)
13q14.11 (rs7328064)
BMIall 1.00 [0.99, 1.00]
TC 1.00 [0.99, 1.00]
HDL-C 1.01 [1.00, 1.02]
LDL-C 1.00 [0.99, 1.01]
TG 1.00 [0.99, 1.01]
BS 1.01 [1.00, 1.01]
HbA1c 1.00 [0.99, 1.02]
BUN 1.00 [0.99, 1.01]
sCr 0.99 [0.98, 1.00]
eGFR 1.01 [1.00, 1.02]
UA 0.99 [0.99, 1.00]
Na 1.01 [1.00, 1.02]
K 1.00 [0.99, 1.01]
Cl 1.01 [1.00, 1.01]
Ca 1.01 [1.00, 1.02]
P 1.00 [0.98, 1.01]
Summary Estimate 1.00 [1.00, 1.00]
0.98 1 1.01 1.03
Odds Ratio (95%CI)
BMIall 1.00 [0.99, 1.01]
TC 1.00 [0.99, 1.01]
HDL-C 1.00 [0.98, 1.01]
LDL-C 1.01 [1.00, 1.03]
TG 1.00 [0.99, 1.01]
BS 0.99 [0.97, 1.00]
HbA1c 0.98 [0.96, 1.00]
BUN 1.00 [0.99, 1.01]
sCr 0.99 [0.98, 1.00]
eGFR 1.02 [1.00, 1.03]
UA 0.99 [0.98, 1.00]
Na 1.01 [1.00, 1.02]
K 1.00 [0.98, 1.01]
Cl 1.00 [0.98, 1.01]
Ca 1.00 [0.98, 1.01]
P 1.01 [0.99, 1.02]
Summary Estimate 1.00 [1.00, 1.00]
0.96 0.98 1 1.02 1.04
Odds Ratio (95%CI)
BMIall 1.00 [0.99, 1.01]
TC 1.00 [0.99, 1.01]
HDL-C 1.01 [1.00, 1.02]
LDL-C 1.00 [0.99, 1.01]
TG 1.00 [1.00, 1.01]
BS 0.99 [0.98, 1.00]
HbA1c 1.01 [1.00, 1.02]
BUN 1.02 [1.01, 1.03]
* sCr 1.02 [1.02, 1.03]
* eGFR 0.98 [0.97, 0.98]
UA 1.01 [1.00, 1.02]
Na 0.99 [0.98, 1.00]
K 1.00 [0.99, 1.01]
Cl 1.00 [0.99, 1.00]
Ca 1.01 [1.00, 1.02]
P 1.02 [1.00, 1.03]
Summary Estimate 1.00 [1.00, 1.00]
0.96 0.98 1 1.02 1.04
Odds Ratio (95%CI)
b
screening 1.10 [1.06, 1.15]
screening2 1.09 [1.05, 1.14]
replication 1.06 [0.97, 1.15]
* Summary Estimate 1.09 [1.06, 1.12]
0.95 1 1.05 1.1 1.15 1.2
Odds Ratio (95%CI)
screening 0.88 [0.83, 0.93]
screening2 0.88 [0.83, 0.93]
replication 0.93 [0.83, 1.04]
* Summary Estimate 0.88 [0.85, 0.92]
0.8 0.9 1 1.05
Odds Ratio (95%CI)
* screening 0.84 [0.80, 0.88]
screening2 0.90 [0.86, 0.94]
* replication 0.89 [0.82, 0.96]
* Summary Estimate 0.87 [0.85, 0.90]
0.8 0.85 0.9 0.95 1
Odds Ratio (95%CI)

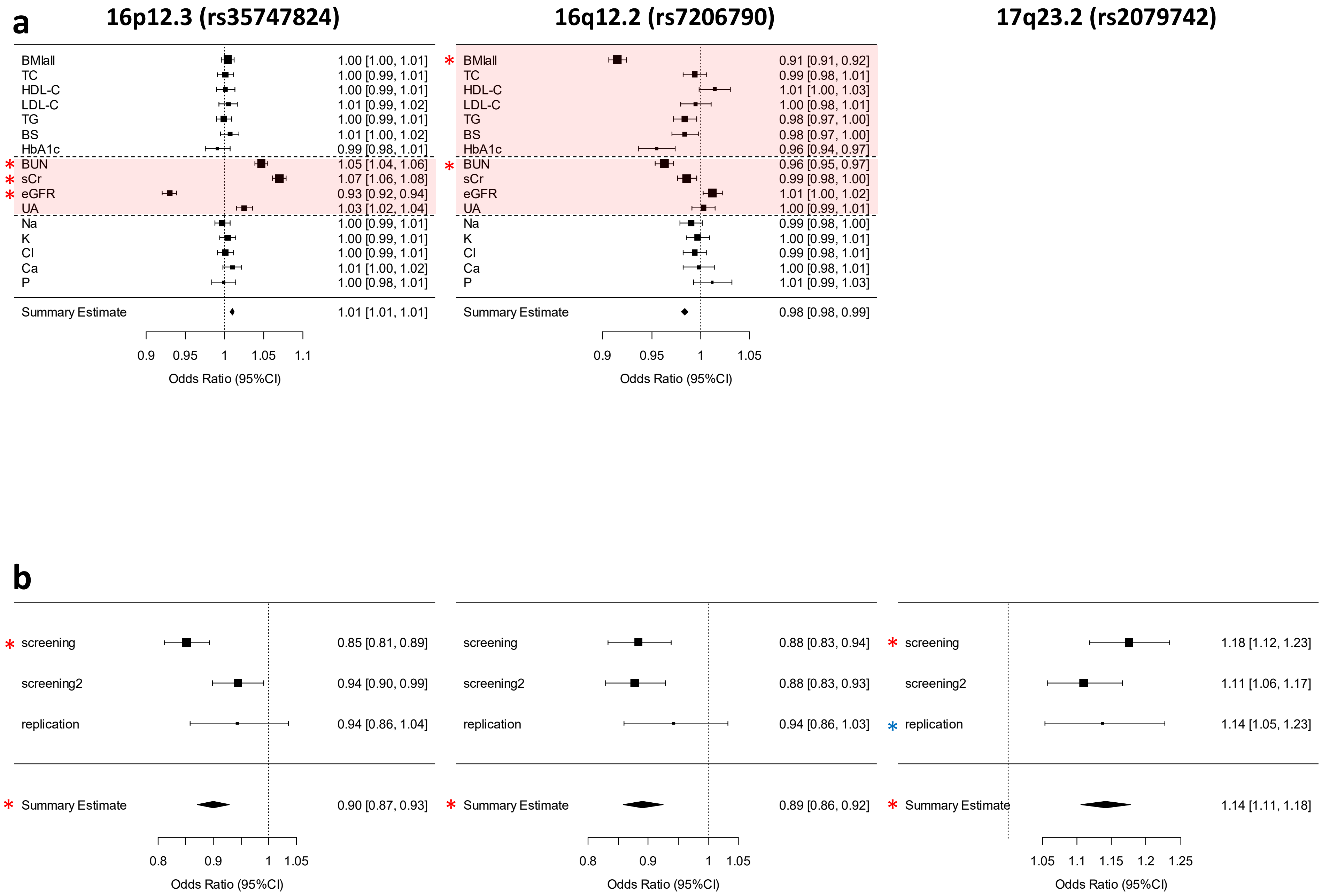
a
16p12.3 (rs35747824)
16q12.2 (rs7206790)
17q23.2 (rs2079742)
BMIall 1.00 [1.00, 1.01]
TC 1.00 [0.99, 1.01]
HDL-C 1.00 [0.99, 1.01]
LDL-C 1.01 [0.99, 1.02]
TG 1.00 [0.99, 1.01]
BS 1.01 [1.00, 1.02]
HbA1c 0.99 [0.98, 1.01]
* BUN 1.05 [1.04, 1.06]
* sCr 1.07 [1.06, 1.08]
* eGFR 0.93 [0.92, 0.94]
UA 1.03 [1.02, 1.04]
Na 1.00 [0.99, 1.01]
K 1.00 [0.99, 1.01]
Cl 1.00 [0.99, 1.01]
Ca 1.01 [1.00, 1.02]
P 1.00 [0.98, 1.01]
Summary Estimate 1.01 [1.01, 1.01]
0.9 0.95 1 1.05 1.1
Odds Ratio (95%CI)
* BMIall 0.91 [0.91, 0.92]
TC 0.99 [0.98, 1.01]
HDL-C 1.01 [1.00, 1.03]
LDL-C 1.00 [0.98, 1.01]
TG 0.98 [0.97, 1.00]
BS 0.98 [0.97, 1.00]
HbA1c 0.96 [0.94, 0.97]
* BUN 0.96 [0.95, 0.97]
sCr 0.99 [0.98, 1.00]
eGFR 1.01 [1.00, 1.02]
UA 1.00 [0.99, 1.01]
Na 0.99 [0.98, 1.00]
K 1.00 [0.99, 1.01]
Cl 0.99 [0.98, 1.01]
Ca 1.00 [0.98, 1.01]
P 1.01 [0.99, 1.03]
Summary Estimate 0.98 [0.98, 0.99]
0.9 0.95 1 1.05
Odds Ratio (95%CI)
b
* screening 0.85 [0.81, 0.89]
screening2 0.94 [0.90, 0.99]
replication 0.94 [0.86, 1.04]
* Summary Estimate 0.90 [0.87, 0.93]
0.8 0.9 1 1.05
Odds Ratio (95%CI)
screening 0.88 [0.83, 0.94]
screening2 0.88 [0.83, 0.93]
replication 0.94 [0.86, 1.03]
* Summary Estimate 0.89 [0.86, 0.92]
0.8 0.9 1 1.05
Odds Ratio (95%CI)
* screening 1.18 [1.12, 1.23]
screening2 1.11 [1.06, 1.17]
* replication 1.14 [1.05, 1.23]
* Summary Estimate 1.14 [1.11, 1.18]
1.05 1.1 1.15 1.2 1.25
Odds Ratio (95%CI)

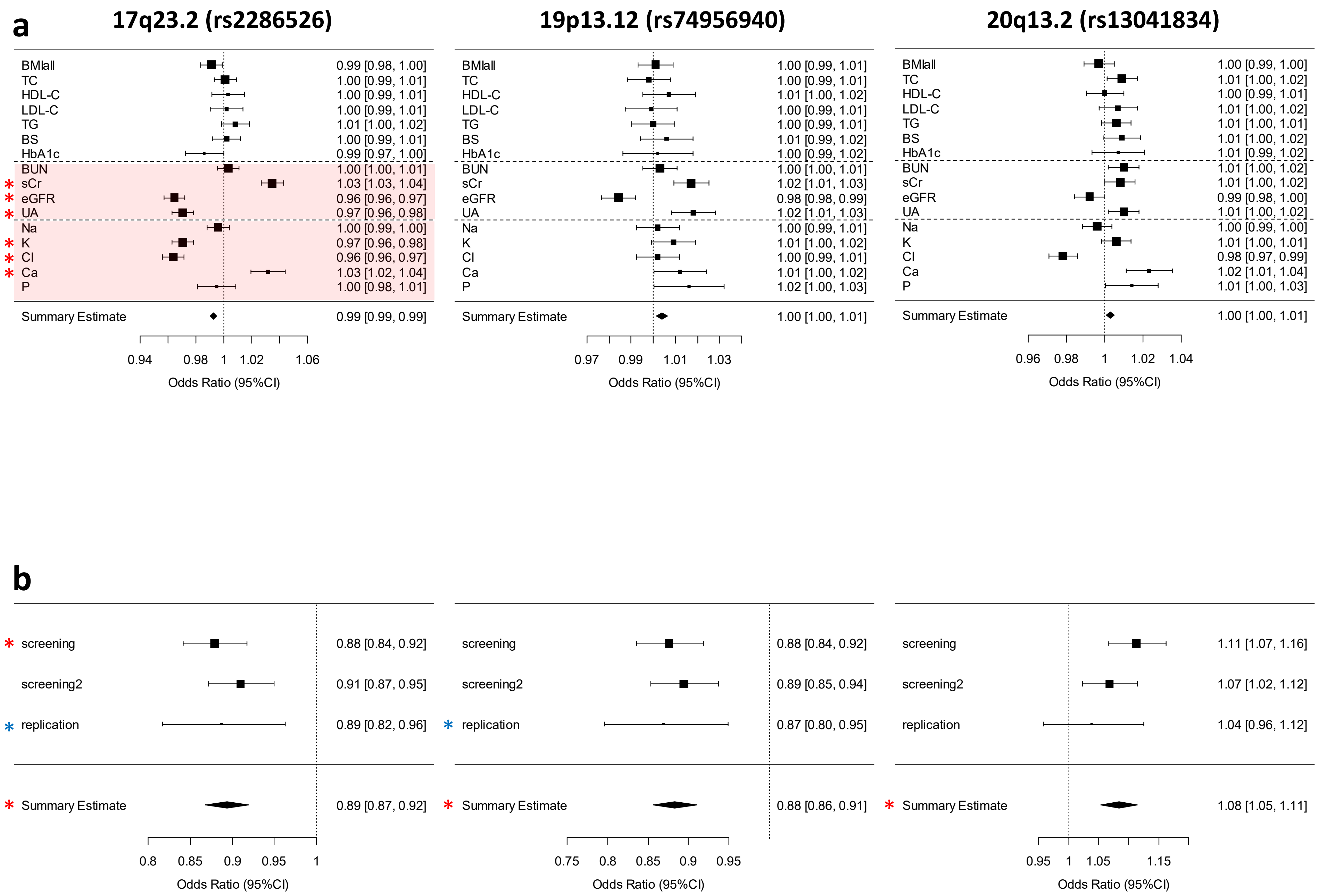

a
17q23.2 (rs2286526)
19p13.12 (rs74956940)
20q13.2 (rs13041834)
BMIall
TC
HDL-C
LDL-C
TG
BS
HbA1c
BUN
sCr
eGFR
UA
Na
K
Cl
Ca
P
Summary Estimate
0.99 [0.98, 1.00]
1.00 [0.99, 1.01]
1.00 [0.99, 1.01]
1.00 [0.99, 1.01]
1.01 [1.00, 1.02]
1.00 [0.99, 1.01]
0.99 [0.97, 1.00]
1.00 [1.00, 1.01]
1.03 [1.03, 1.04]
0.96 [0.96, 0.97]
0.97 [0.96, 0.98]
1.00 [0.99, 1.00]
0.97 [0.96, 0.98]
0.96 [0.96, 0.97]
1.03 [1.02, 1.04]
1.00 [0.98, 1.01]
0.99 [0.99, 0.99]
1.00 [0.99, 1.01]
1.00 [0.99, 1.01]
1.01 [1.00, 1.02]
1.00 [0.99, 1.01]
1.00 [0.99, 1.01]
1.01 [0.99, 1.02]
1.00 [0.99, 1.02]
1.00 [1.00, 1.01]
1.02 [1.01, 1.03]
0.98 [0.98, 0.99]
1.02 [1.01, 1.03]
1.00 [0.99, 1.01]
1.01 [1.00, 1.02]
1.00 [0.99, 1.01]
1.01 [1.00, 1.02]
1.02 [1.00, 1.03]
1.00 [1.00, 1.01]
1.00 [0.99, 1.00]
1.01 [1.00, 1.02]
1.00 [0.99, 1.01]
1.01 [1.00, 1.02]
1.01 [1.00, 1.01]
1.01 [1.00, 1.02]
1.01 [0.99, 1.02]
1.01 [1.00, 1.02]
1.01 [1.00, 1.02]
0.99 [0.98, 1.00]
1.01 [1.00, 1.02]
1.00 [0.99, 1.00]
1.01 [1.00, 1.01]
0.98 [0.97, 0.99]
1.02 [1.01, 1.04]
1.01 [1.00, 1.03]
1.00 [1.00, 1.01]
0.94 0.98 1 1.02 1.06
0.97 0.99 1.01 1.03
0.96 0.98 1 1.02 1.04
Odds Ratio (95%CI)
b
screening
screening2
replication
Summary Estimate
0.88 [0.84, 0.92]
0.91 [0.87, 0.95]
0.89 [0.82, 0.96]
0.89 [0.87, 0.92]
0.88 [0.84, 0.92]
0.89 [0.85, 0.94]
0.87 [0.80, 0.95]
0.88 [0.86, 0.91]
1.11 [1.07, 1.16]
1.07 [1.02, 1.12]
1.04 [0.96, 1.12]
1.08 [1.05, 1.11]
0.8 0.85 0.9 0.95 1
0.75 0.8 0.85 0.9 0.95
0.95 1 1.05 1.15
Odds Ratio (95%CI)

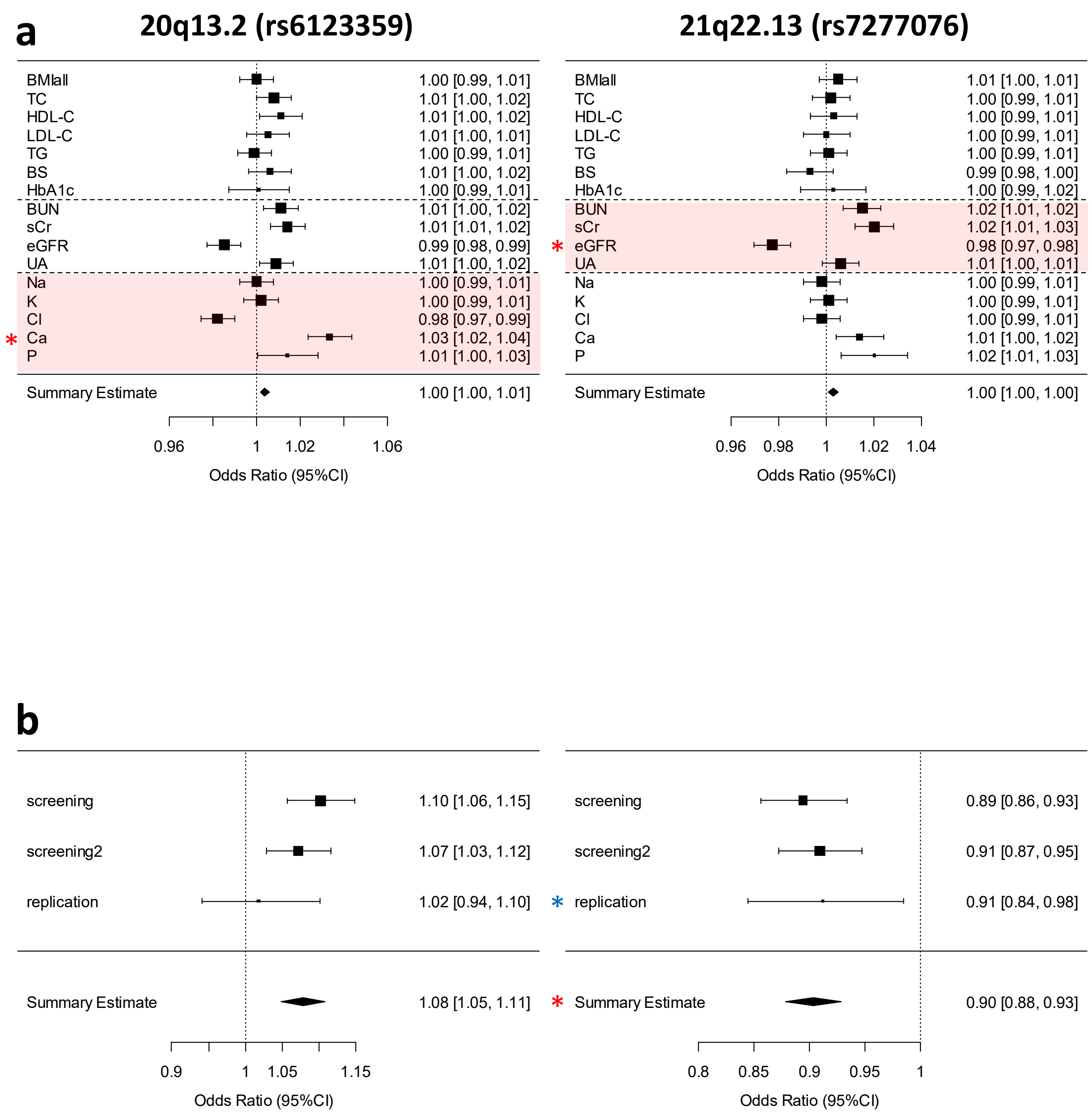


**Supplemental Figure 4 Forest plots for risk variants in the 17 nephrolithiasis risk loci.**
(**a**) Plots show the association estimates (odds ratios) and 95% confidence intervals for GWAS of 16 quantitative traits presented as bars. (**b**) Plots show the association estimates (odds ratios) and 95% confidence intervals for the screening stage and replication study presented as bars. The association estimate and confidence interval for the meta-analysis combining screening and replication result are shown as a diamond.

#### Supplemental Figure 5

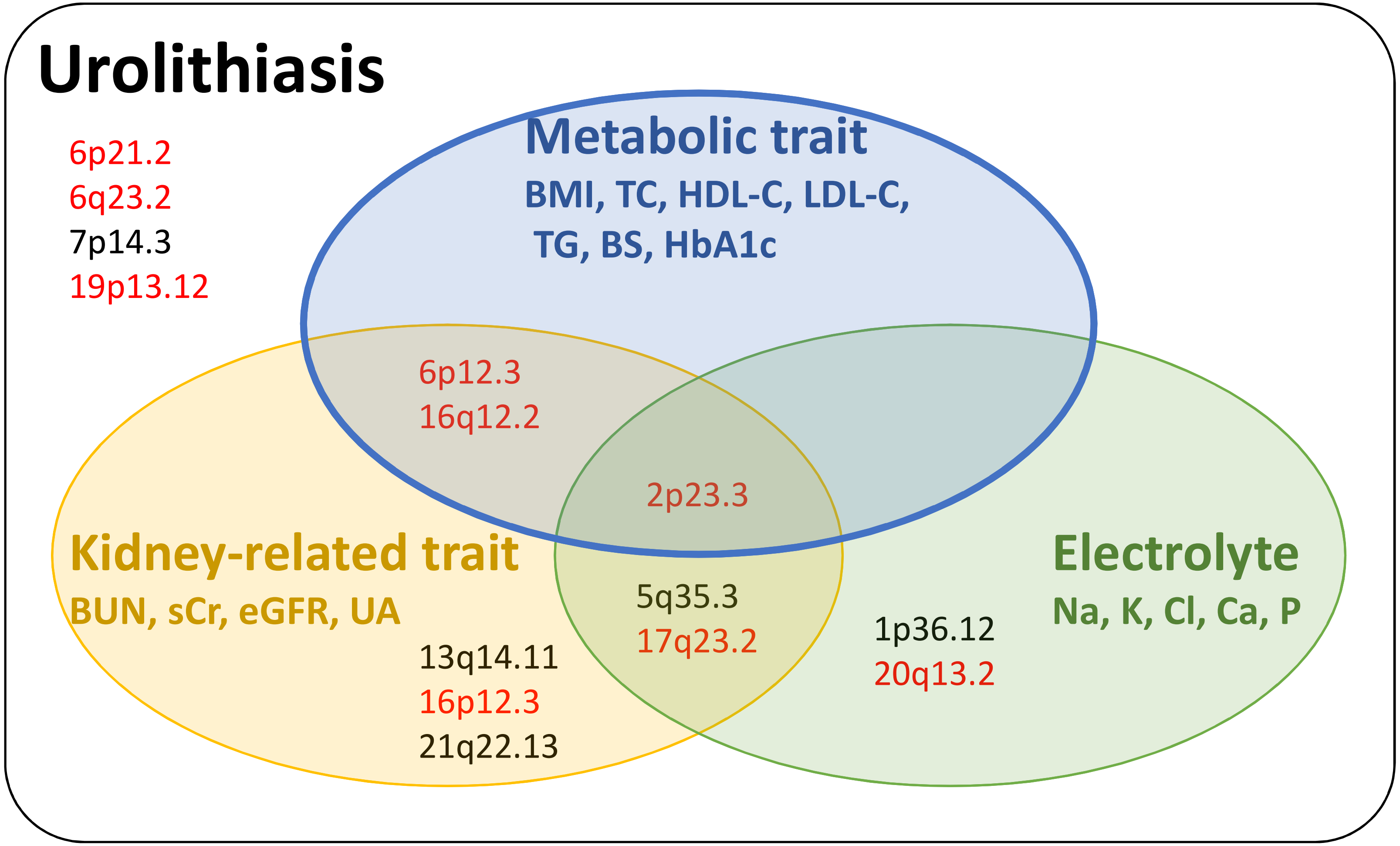


**Supplemental Figure 5 Summary of pleiotropy analysis.**
Venn diagram showing 14 regions that are associated with 16 quantitative traits in three categories including metabolic, kidney related, and electrolyte.
