## Supplementary data2 for "GWAS identifies nine nephrolithiasis susceptibility loci related with metabolic metabolic and crystallization pathways"

### Supplemental Figure 6

## 1p36.12 (rs6667242)

a

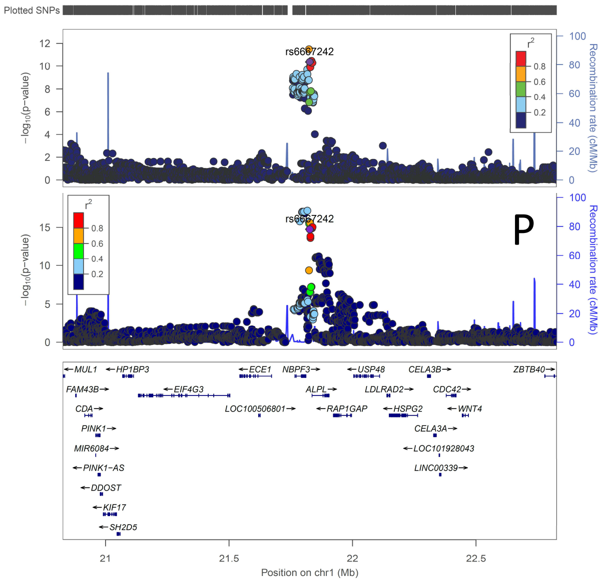


b

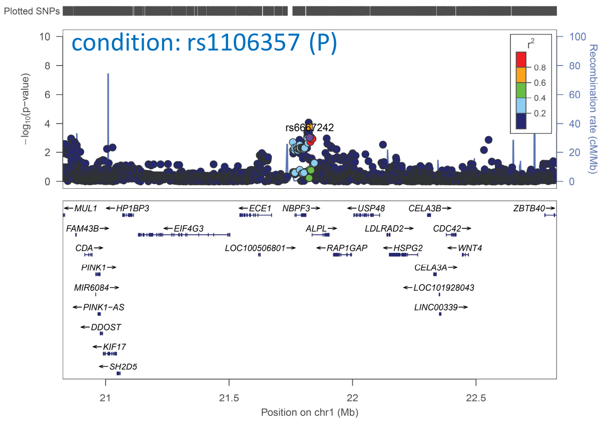

## 2p23.3 (rs1260326)

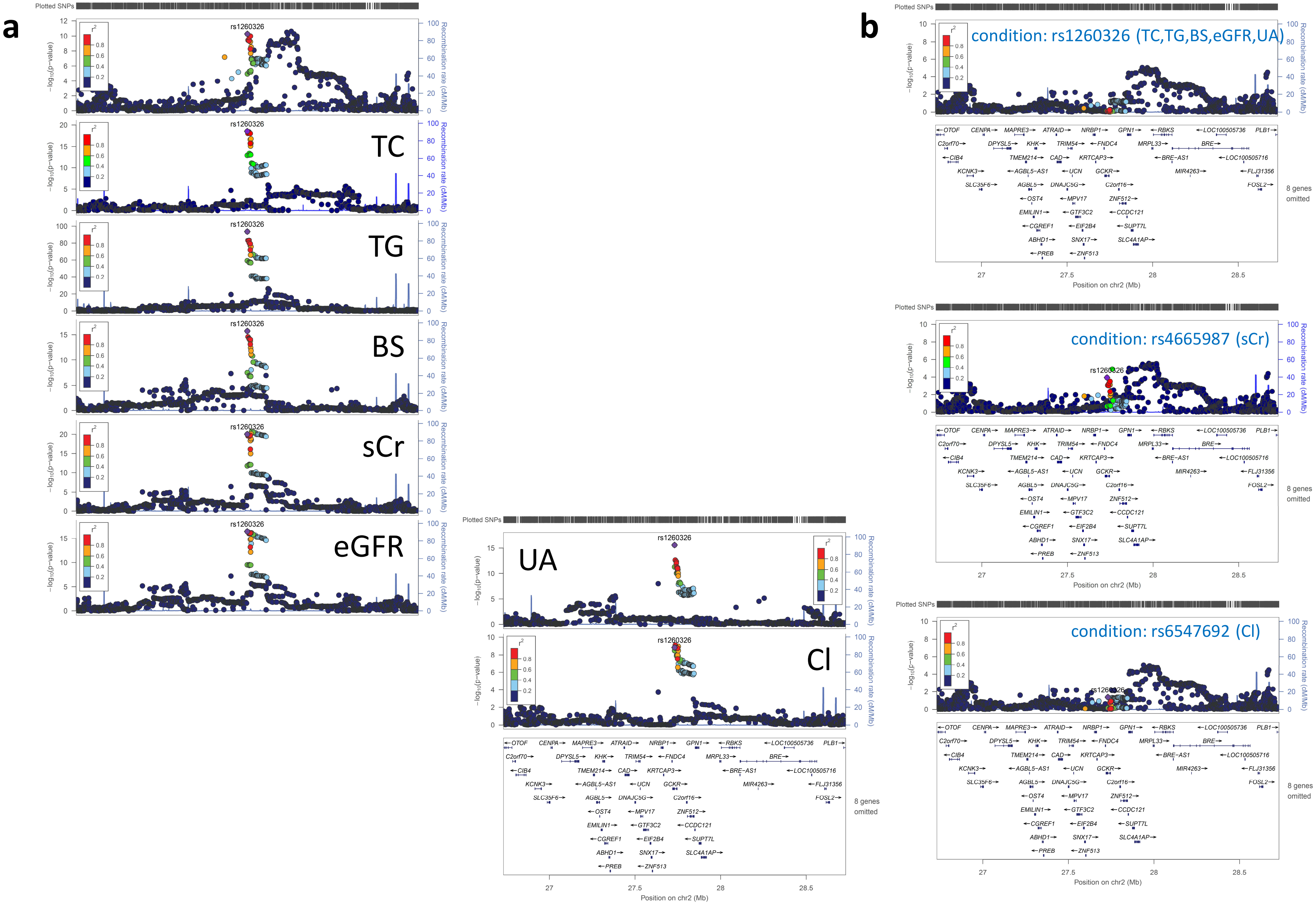

# 5q35.3 (rs11746443)

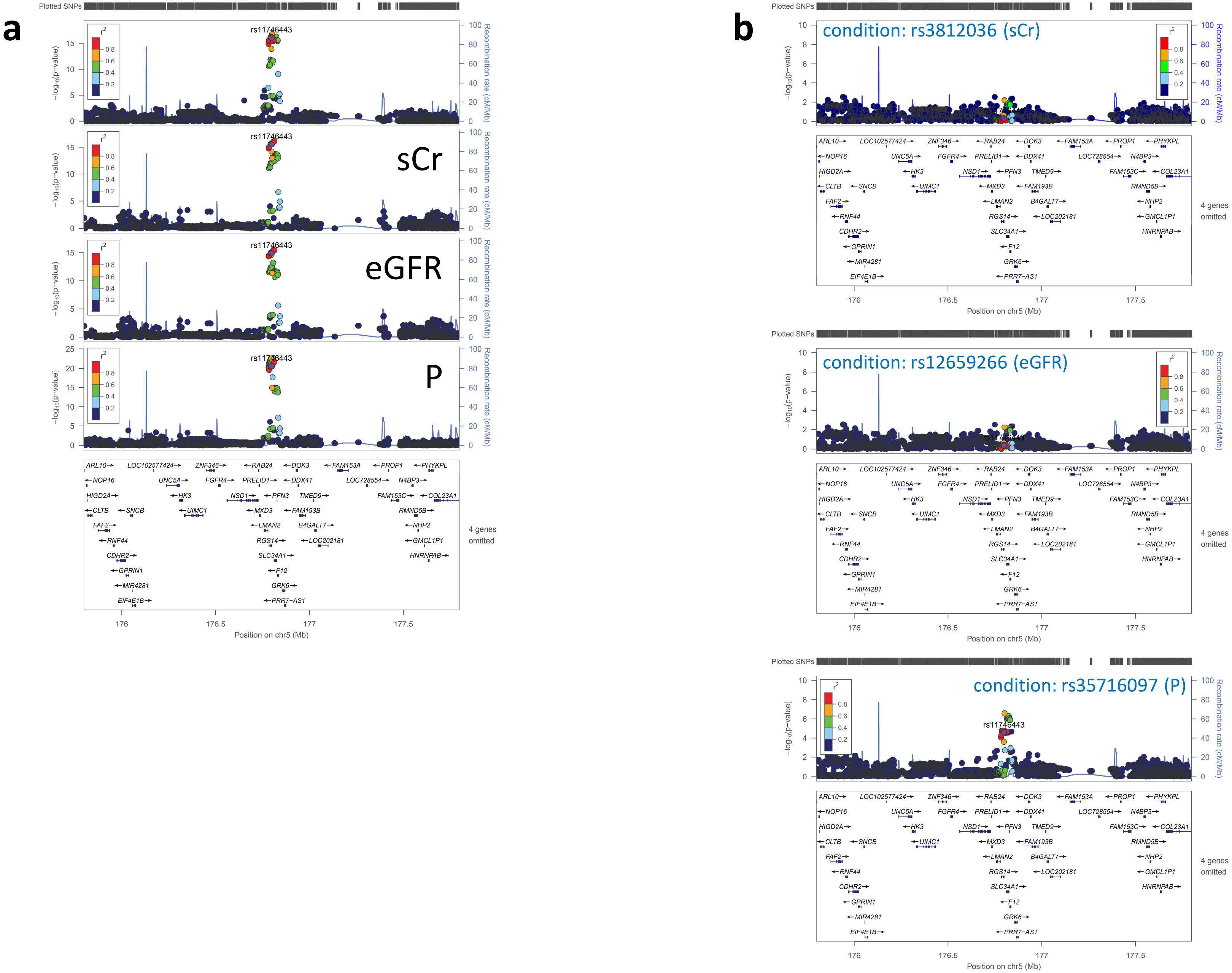

## 6p12.3 (rs3798519)

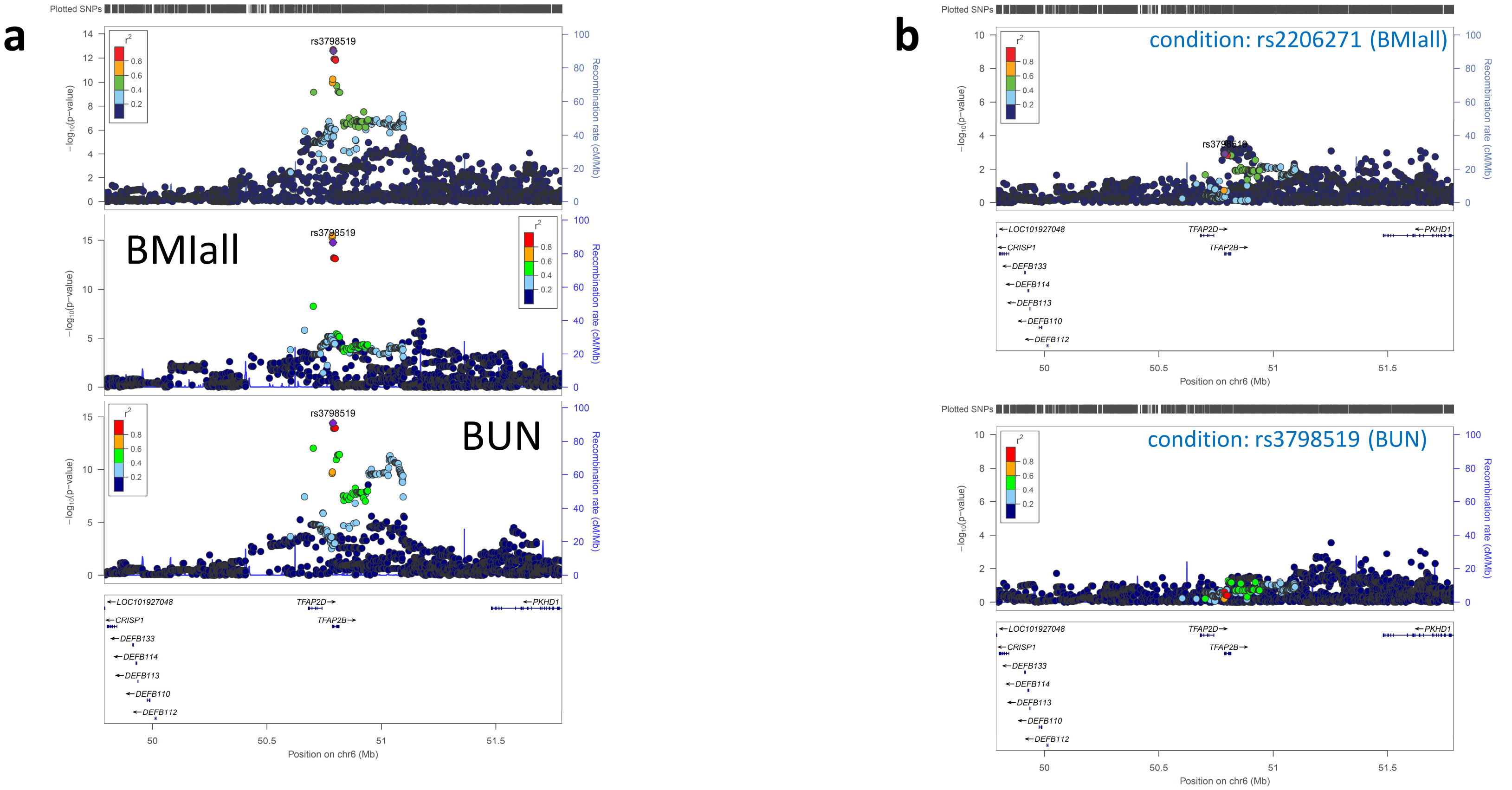

## 13q14.11 (rs7328064)

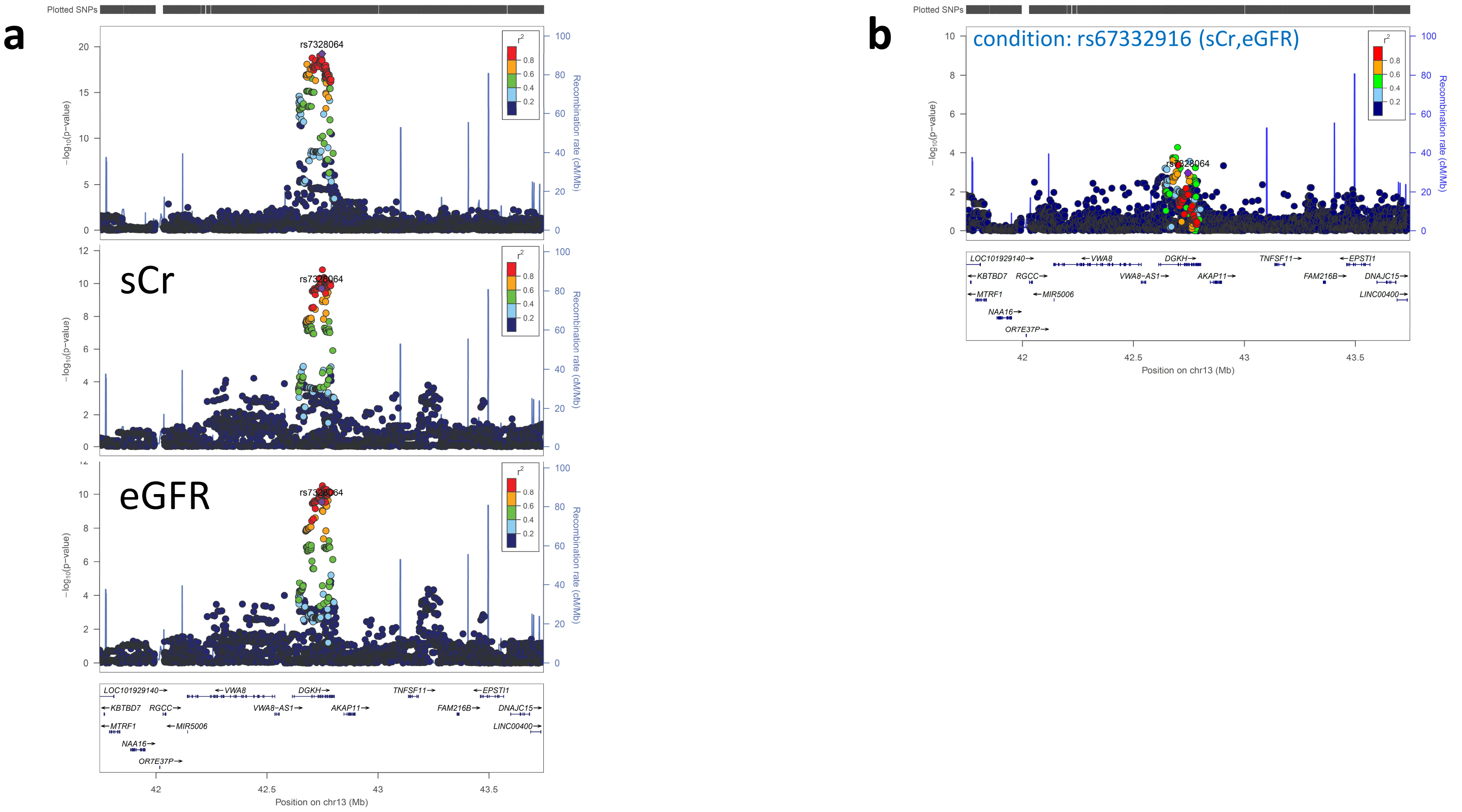

## 16p12.3 (rs35747824)

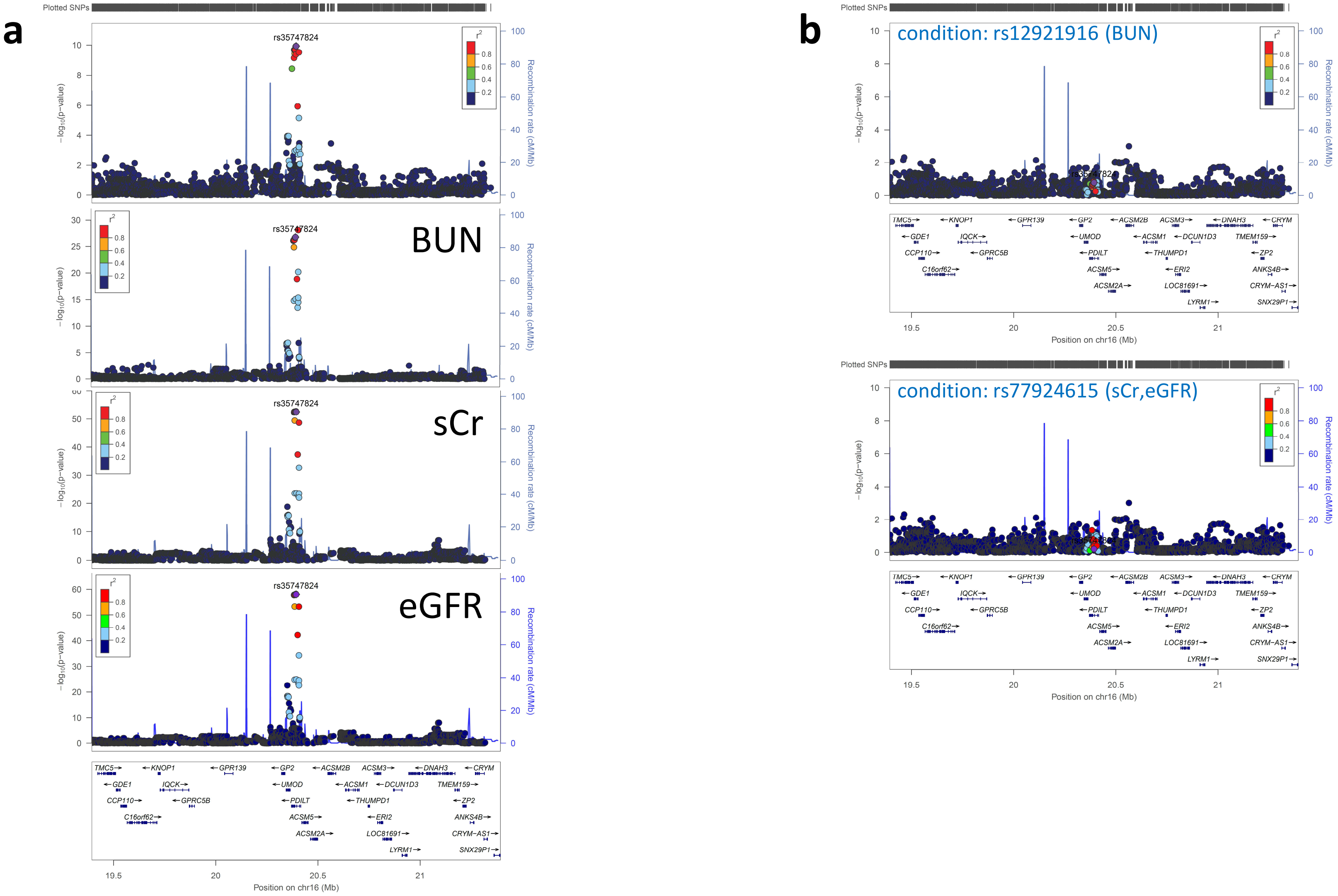

## 16q12.2 (rs7206790)

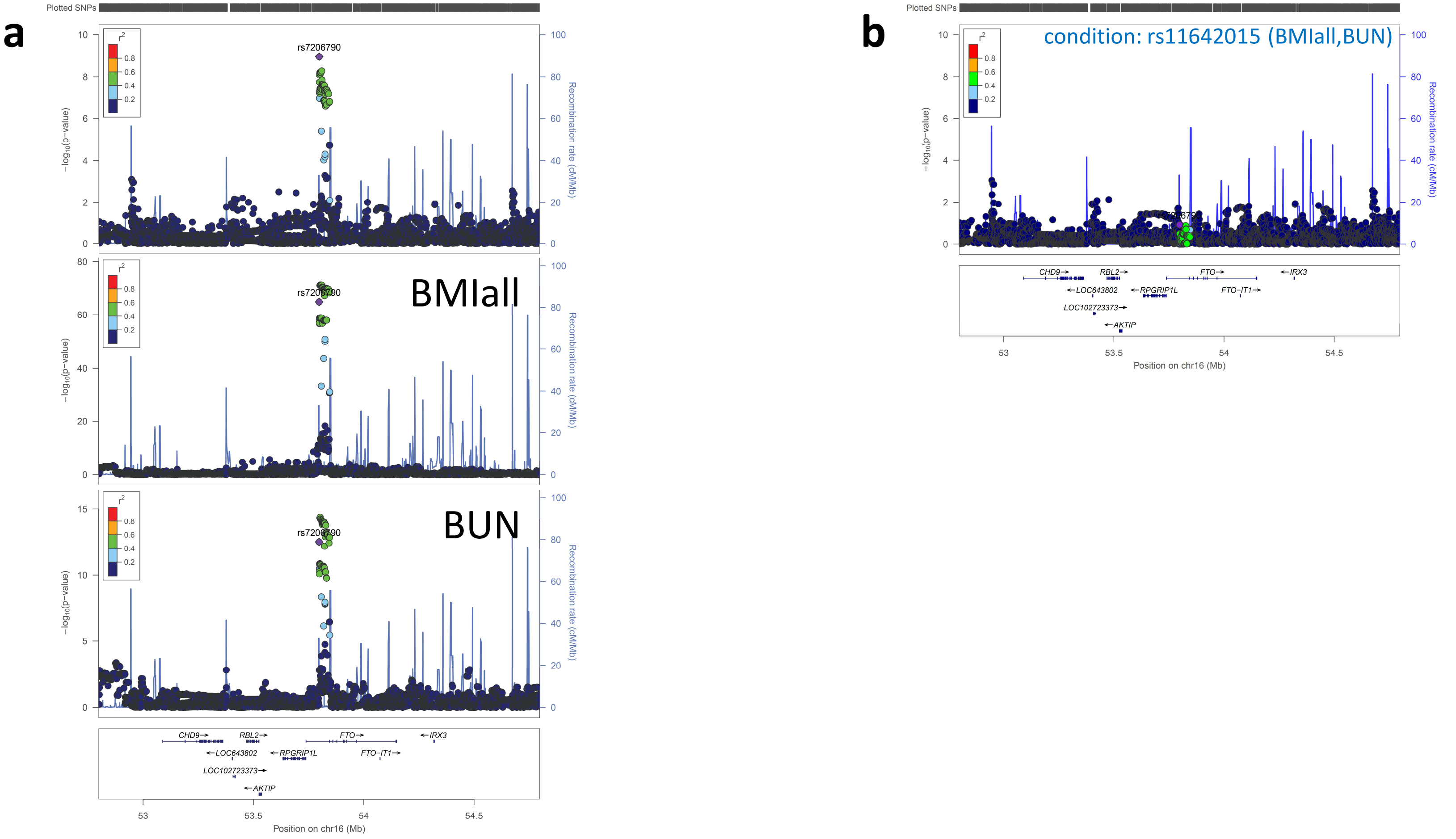

## 17q23.2 (rs2286526)

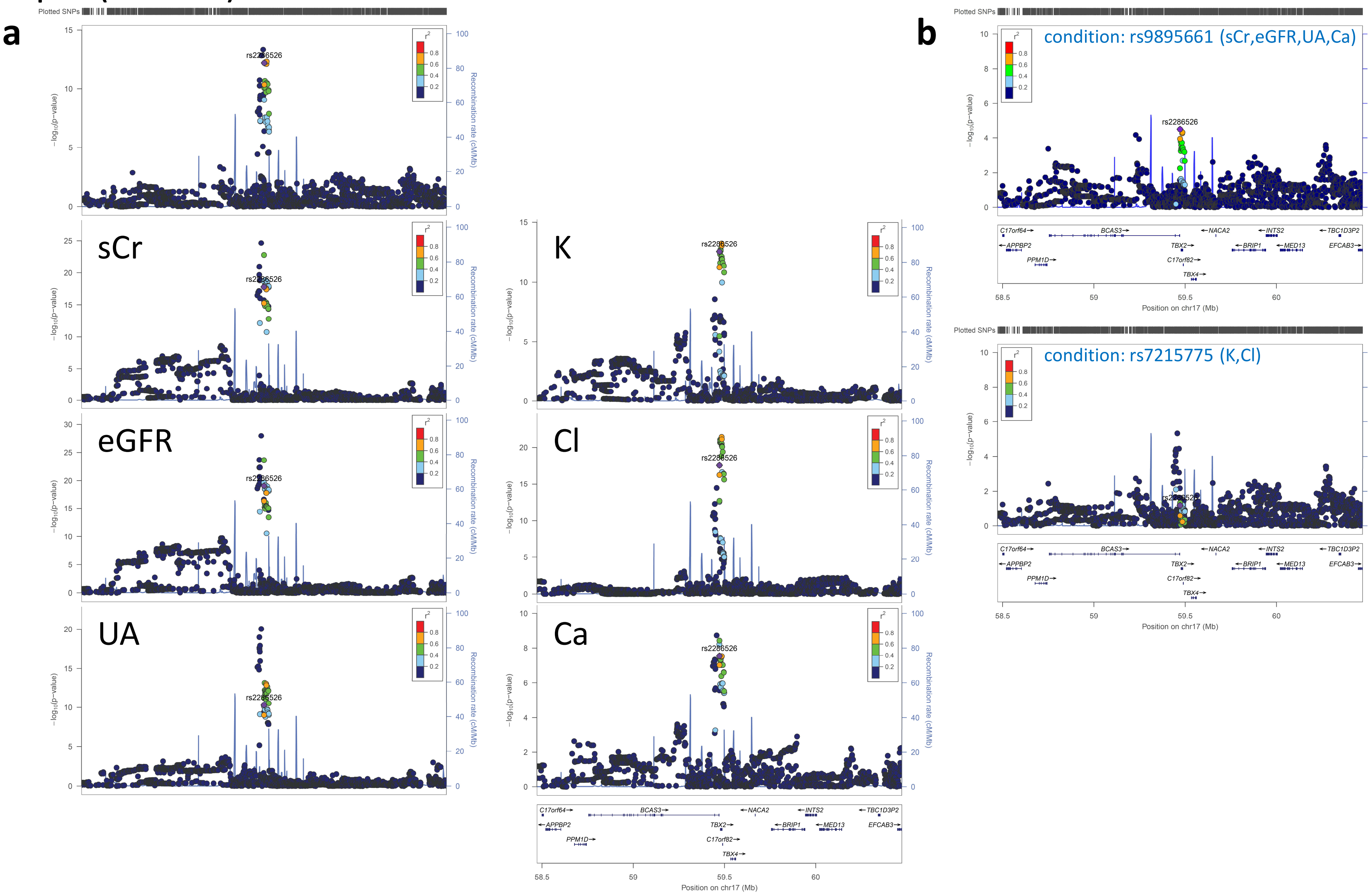

## 20q13.2 (rs6123359)

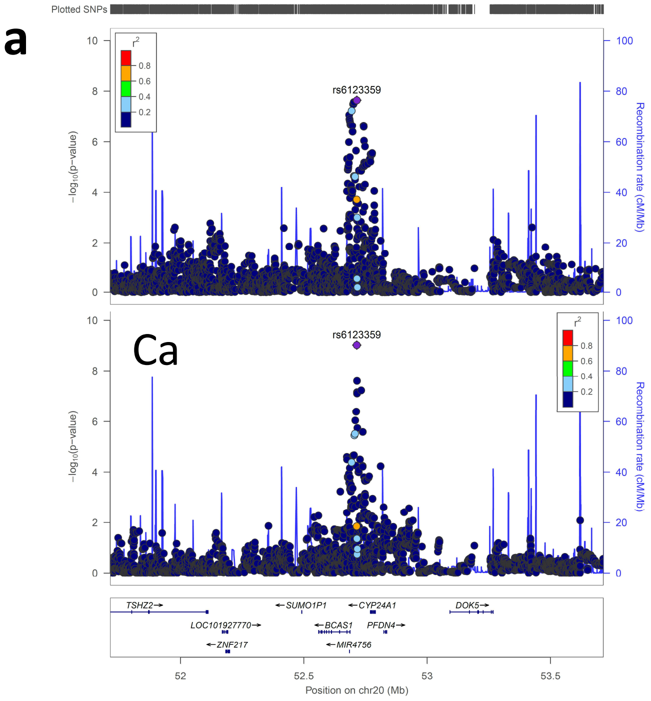


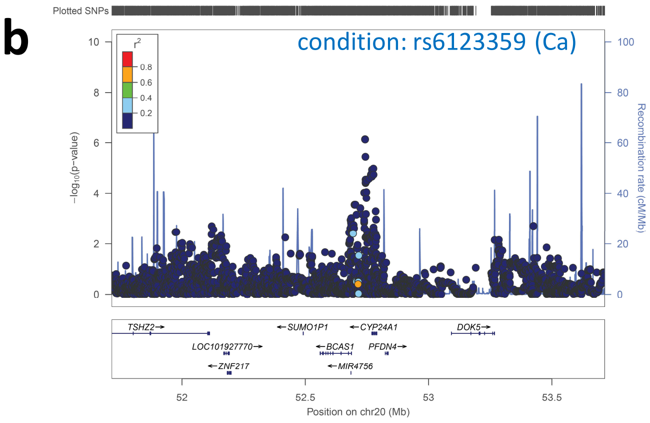

## 21q22.13 (rs7277076)

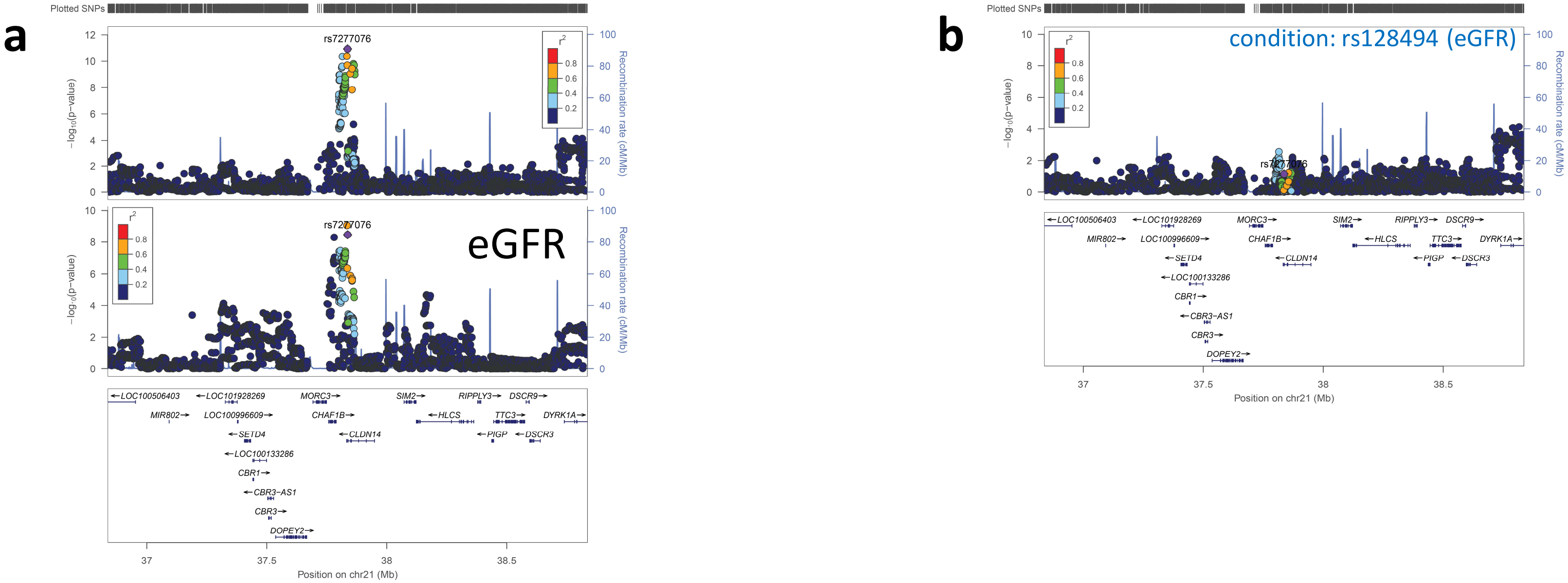


**Supplemental Figure 6 Regional plots of 10 pleiotropic loci before and after condition.**

(**a**) The $-\log_{10}$ *P* values from the meta-analysis of screening 1 and screening 2 (top panel) and GWAS of 16 quantitative traits are shown. (**b**) Regional plots after condition. Estimated recombination rates (from 1000 Genomes) are plotted in blue. The SNPs are color-coded to reflect their correlation with genotyped SNPs. Pairwise r2 values are from 1000 Genomes East Asian data (March 2012 release). Genes, position of exons and direction of transcription from UCSC genome browser (genome.ucsc.edu) are noted. Plots were generated using LocusZoom (http://csg.sph.umich.edu/locuszoom).

### Supplemental Figure 7

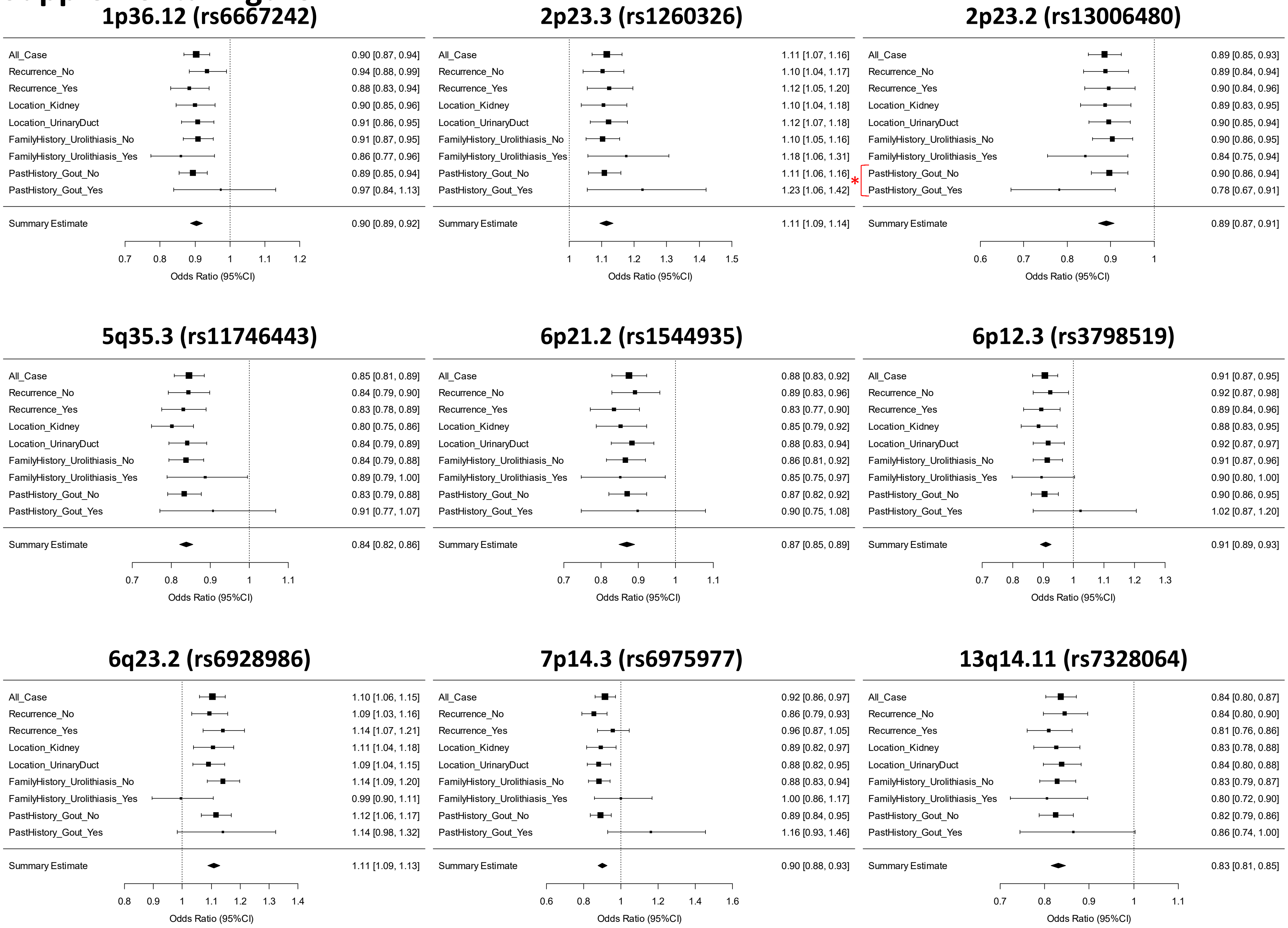

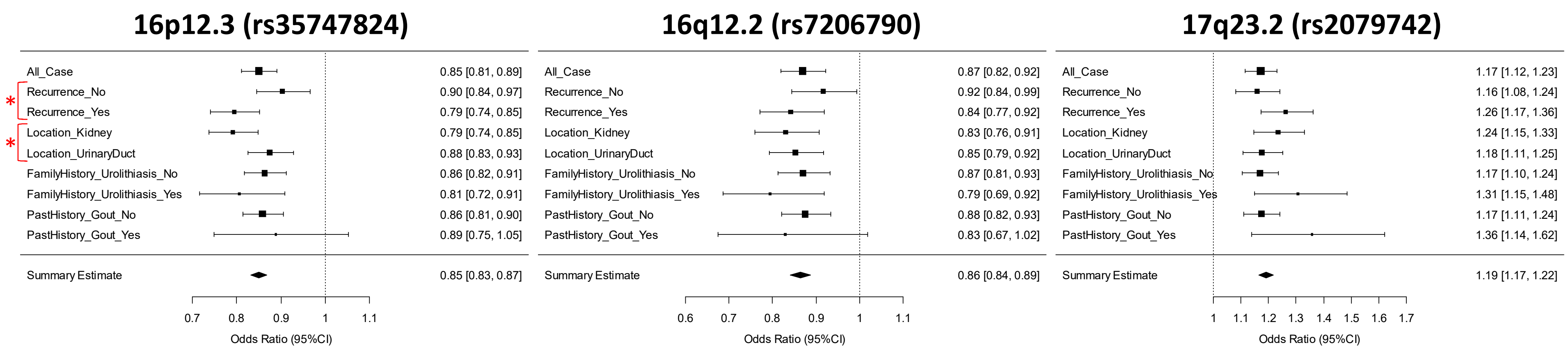


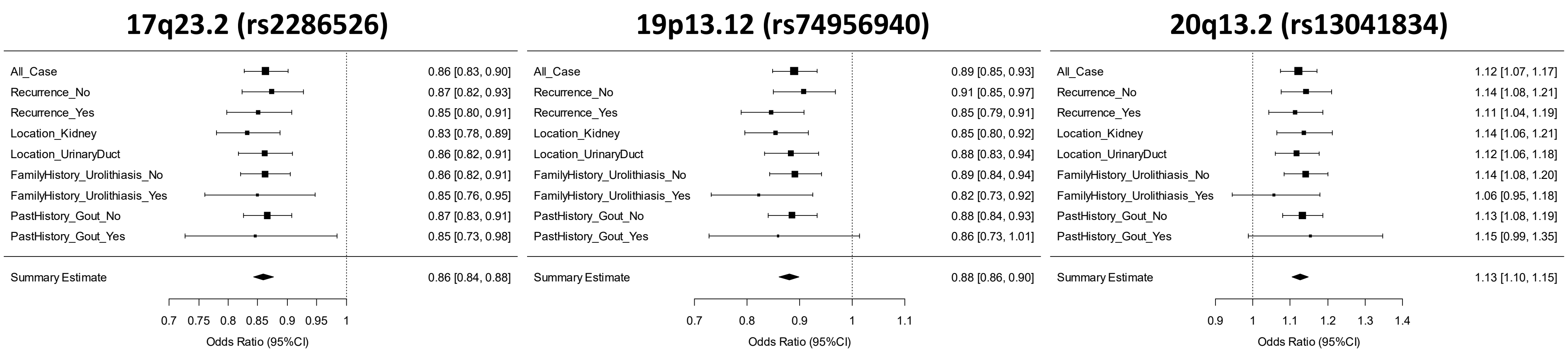


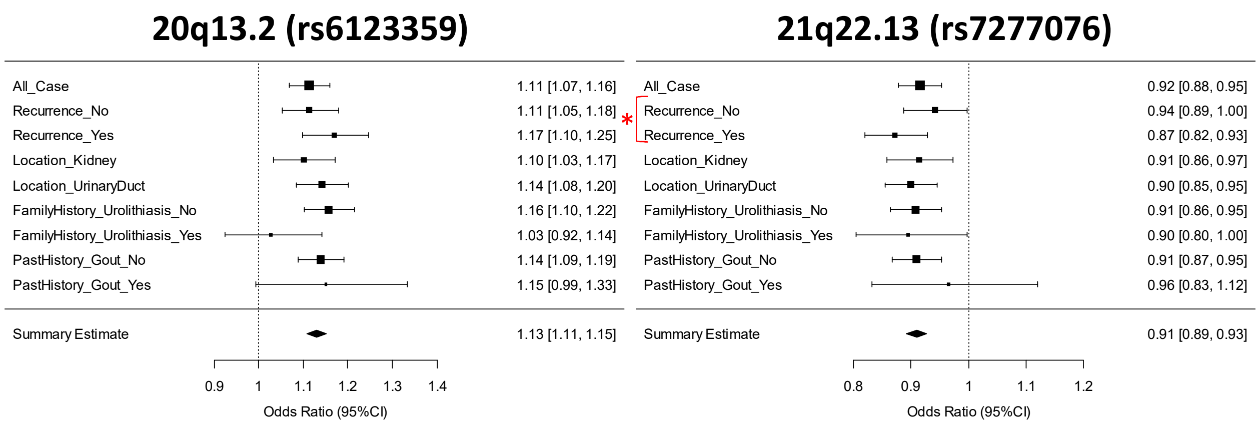


**Supplemental Figure 7 Forest plots for risk variants in the 17 nephrolithiasis risk loci.** Plots show the association estimates (odds ratios) and 95% confidence intervals for subgroup analysis presented as bars. *P<0.05.

### Supplemental Figure 8

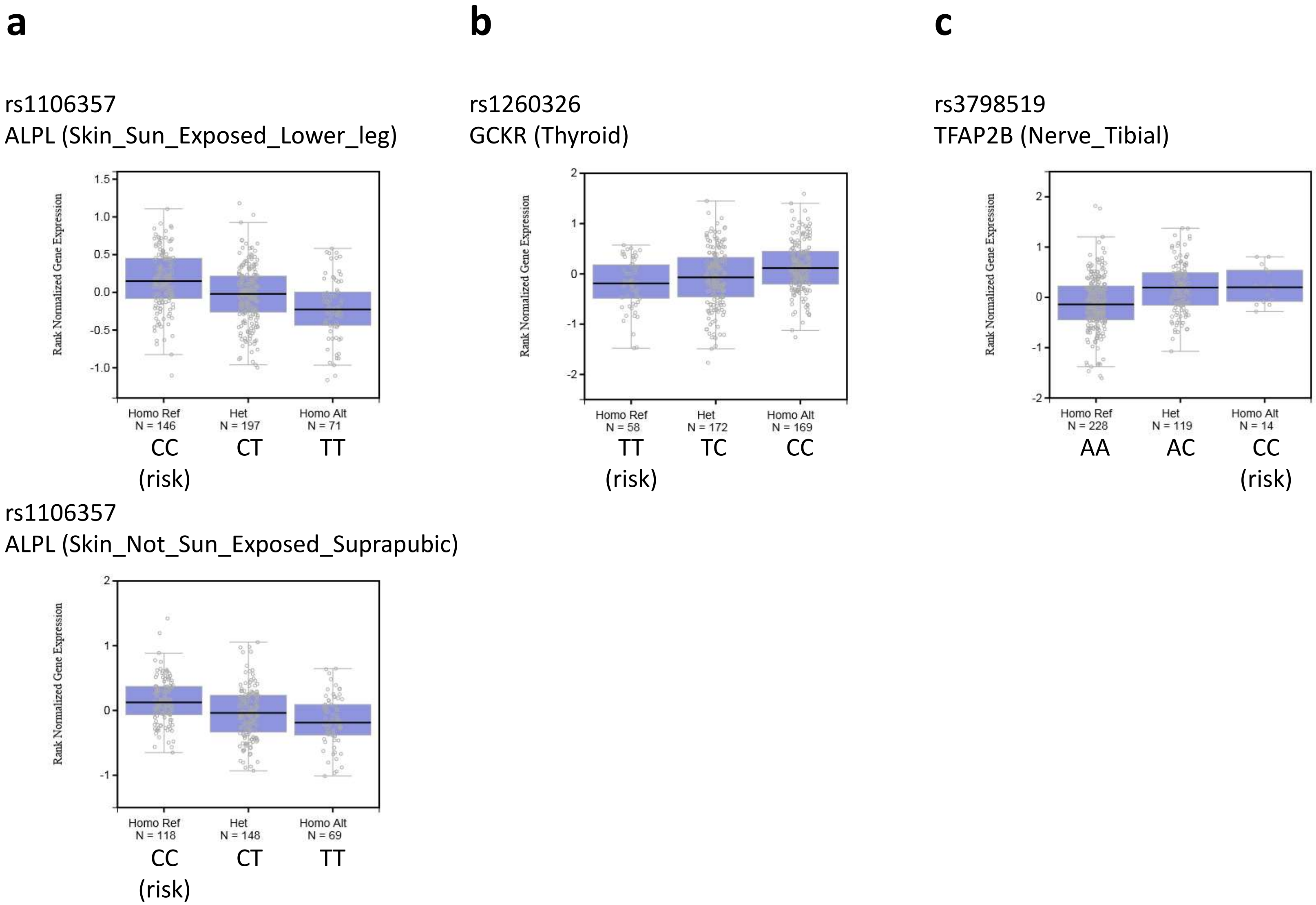


**Supplemental Figure 8 eQTL analysis.** (**a**) Association of rs1106357 with *ALPL* expression in skins. (**b**) Association of rs1260326 with *GCKR* expression in thyroid. (**c**) Association of rs3798519 with *TFAP2B* expression in nerve. Data were derived from GTEx portal (https://www.gtexportal.org/home/).

### Supplemental Figure 9

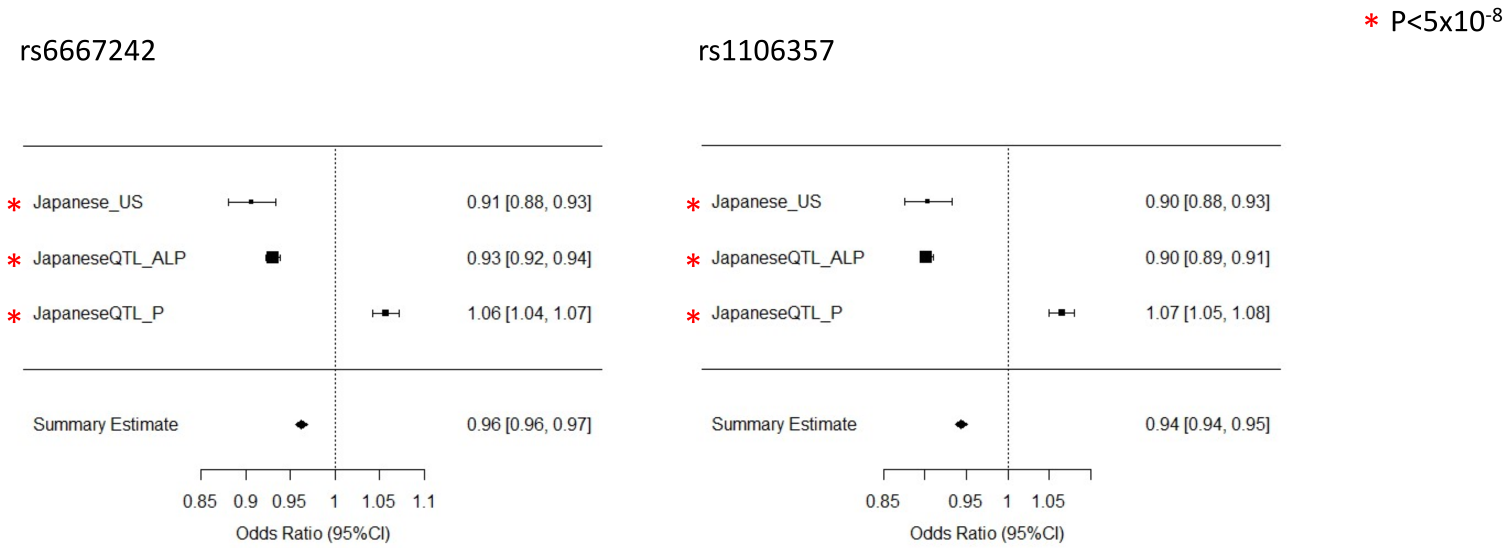


**Supplemental Figure 9 Forest plots for risk variants of rs6667242 and rs1106357.** Plots show the association estimates (odds ratios) and 95% confidence intervals for the meta-analysis of screening stages (top) and GWAS of ALP (middle) and GWAS of P (bottom) presented as bars.
